# The zoo of the gene networks capable of pattern formation by extracellular signaling

**DOI:** 10.1101/2025.05.06.652477

**Authors:** Kevin Martinez-Anhom, Isaac Salazar-Ciudad

## Abstract

A fundamental question of developmental biology is pattern formation, or how cells with specific gene expression end up in specific locations in the body to form tissues, organs and, overall, functional anatomy. Pattern formation involves communication through extracellular signals and complex intracellular gene networks integrating these signals to determine cell responses (e.g., further signaling, cell division, cell differentiation, etc.). In this article we address two question: 1) Are there any logical or mathematical principles determining which gene network topologies can lead to pattern formation by cell signaling over space in multicellular systems? 2) Can gene network topologies be classified into a small number of classes that entail similar dynamics and pattern transformation capacities?

We combine logical arguments and mathematical proofs to show that, despite the large amount of formally possible gene network topologies, all gene network topologies capable of pattern formation fall into only three fundamental classes and their combinations. Additionally, we show that gene networks within each class share the same logic on how they lead to pattern formation and hence, lead to similar patterns. We characterize the main features of each class. This zoo includes the complete gene networks that, to the best of our knowledge, have been experimentally reported to lead to pattern formation as well as other gene networks that have not yet been found experimentally.

**Significance Statement:** Pattern formation is a central problem in developmental biology, yet the principles linking gene regulatory network topology to spatial patterning in multicellular systems are not fully understood. This work identifies logical and mathematical principles that determine which gene regulatory network topologies can give rise to spatial pattern formation through extracellular signaling. The work further shows that these topologies can be systematically classified into a small number of fundamental classes associated with distinct dynamical behaviors. Despite the vast number of *a priori* possible gene network configurations, the gene networks capable of stationary pattern formation fall into just three fundamental topological classes and their combinations. Gene networks within each class implement a common patterning logic and consequently generate analogous pattern transformations, revealing a unifying organizational framework underlying biological pattern formation.

## Introduction

Development is the process by which the intricate complexity of multicellular organisms is constructed from a single fertilized egg or some simple vegetative structure (Fusco and Minelli, 2023). Not many other natural processes lead to so much complexity in such a short time. Development entails a natural process of pattern formation: specific cells and cell types end up in specific positions in space (i.e., anatomy). This process of pattern formation can be seen as the generation of spatial information (i.e., information about where each cell and cell type are) from previous information (e.g., information within the fertilized egg). This latter information includes the DNA but also spatial information in the form of compartments with different proteins and RNAs in different spatial locations within the oocyte. In most species, development cannot proceed if this latter spatial information is removed experimentally (Gilbert and Barresi, 2023). This initial spatial information within the oocyte arises from spatial asymmetries in the mother’s gonads or from the environment (Gilbert and Barresi, 2023). In this sense, pattern formation in development does not usually start from a spatially homogeneous initial condition but from an initial condition that has some simple spatial heterogeneities. In that sense, we use the term pattern transformation instead of pattern formation (Salazar-Ciudad *et al*., 2003).

Development can be seen as a sequence of transformations between initial developmental patterns and latter developmental patterns (what we call *resulting patterns*) over developmental time. By developmental pattern, or simply pattern, we mean a specific distribution of cell types in space or a specific distribution of gene product concentrations over space. For example, the earliest such patterns would be the zygote and the latest the adult. In most animals, early patterns arise from the division of the fertilized egg into different cells that, thus, inherit different parts of the fertilized egg and different proteins and RNAs (Gilbert and Barresi, 2023). Some of these latter molecules act as transcription factors that lead to the expression of further genes, i.e., the transcription and translation of genes and the eventual synthesis of their gene products (Gilbert and Barresi, 2023). Other gene products are secreted and diffuse in the extracellular space and bind to specific receptors in distant cells (Gilbert and Barresi, 2023). Here, we call these molecules extracellular signals but in the literature they are also called morphogens, growth factors, paracrine factors, etc. (Gilbert and Barresi, 2023).

Cells can respond to extracellular signals by changing gene expression and, often, by secreting additional extracellular signals and regulating cell behaviors such as cell division, cell contraction, cell adhesion, cell death, etc. (Salazar-Ciudad *et al*. 2003; Gilbert and Barresi, 2023). The former type of response leads to further changes in gene expression over cells (i.e., further pattern transformations), while the latter leads to cell movement and, consequently, to changes in the distribution of cells and gene expression over space (i.e., further pattern transformations) (Salazar-Ciudad *et al*. 2003).

How cells respond to extracellular signals depends on the signal receptors and signal transduction pathways they express (Gilbert and Barresi, 2023). Signal transduction pathways are actually networks of molecular interactions that integrate incoming signals to determine cell responses. Here, we use the term gene network to refer to these networks of interactions, even if they also include molecules that are not gene products (Gilbert and Barresi, 2023).

A fundamental question in developmental biology is how pattern transformations occur. In animals, pattern transformation involves gene networks, signaling by extracellular signals and the regulation of cell behaviors (e.g. cell division, cell contraction, cell adhesion) and the mechanical properties of cells and tissues (Salazar-Ciudad, et al., 2003). As a preliminary step to understand pattern transformation by these processes, we restrict ourselves to the pattern transformations occurring through gene networks and extracellular signaling alone (not including membrane-tethered signals or mechano-transduction). We ask:

1. Which are the gene networks topologies that can lead to pattern transformation?
2. Can we identify all these topologies and classify them into classes leading to similar pattern transformations?

Among pattern transformations we are only interested in those that are non-trivial. By non-trivial pattern transformations we mean transformations in which:

P1. The resulting pattern is stationary and heterogeneous in space.

P2. There is at least one gene product that, in the resulting pattern, has a new spatial distribution of stationary concentration maxima and minima over space (i.e. critical points of the concentration distribution over space, see S1 of the Appendix). By new we mean that no gene product had this spatial distribution of concentration maxima and minima in the initial pattern (see Fig.1). Thus, non-trivial pattern transformations include the emergence of new maxima or minima, the disappearance of existing ones, or their spatial displacement from one location to another. This excludes the simple widening of already existing concentration maxima, since, in this case, the position of the maxima does not change (see S1 in the appendix for further details). We only consider resulting developmental patterns (and thus new concentration maxima and minima) that are stable in time. We say that a gene product has been patterned if the spatial distribution of its maxima and minima is new (i.e. not present in any gene product in the initial pattern).

**Figure 1.**
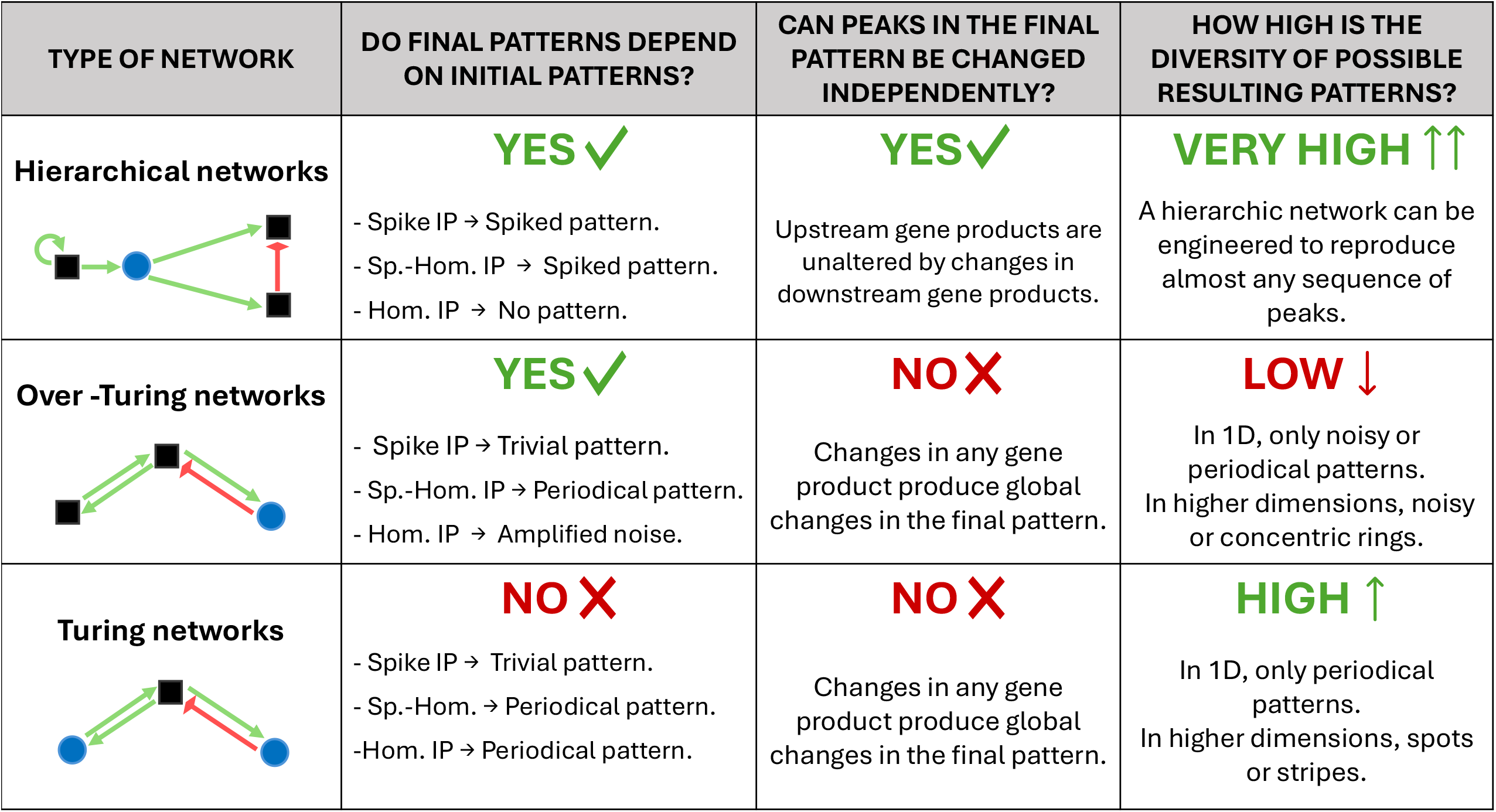
Pattern transformations. The figure shows examples of pattern transformations in 1D **(A-D)** and in 2D **(E-H).** Panels represent different patterns (i.e., different spatial distributions of some gene product concentration *g* (*t, x*)). (**A**) and **(E)** are the initial patterns being transformed into the resulting patterns in **(B-D)** and **(F-H)**. The x-axis is cell position along a 1D array of cells and the y-axis is the concentration of a gene product. The transformation from **(A)** (resp., **(E)**) into **(B)** (resp. **(F)**) is trivial because the resulting pattern is homogeneous. The transformation from **(A)** (resp., **(E)**) to **(C)** (resp., **(G)**) is trivial because the concentration maximum is at the same spatial position in both patterns. The transformation from **(A)** (resp., **(E)**) to **(D)** (resp., **(H)**) is non-trivial because the resulting pattern is heterogeneous and the pattern in **(D)** has maxima that were not present in **(A)**. The blue, grey and pink fillings in **(A-D)** correspond to the sign of the derivative and are meant to highlight the maxima and minima. In **(E-H)** the colors represent gene product concentration.

There are several previous theoretical studies exploring how gene products can be wired to lead to pattern transformations through cell signaling (Salazar-Ciudad *et al*., 2000, 2001; Cotterel and Sharpe, 2010; Marcon *et al*., 2016; Zheng *et al*., 2016; Jimenez *et al*., 2017; Diego *et al*., 2018 ; Leyshon *et al*., 2021). Some of these studies are restricted to networks of three gene products (Cotterel and Sharpe, 2010; Zheng *et al*., 2016) while others are restricted to studying a specific class of gene network topology (Marcon *et al*., 2016; Zheng *et al*., 2016; Jimenez *et al*., 2017; Rand *et al*., 2021; Leyshon *et al*., 2021). In a previous study, we addressed the same questions considered here and identified two classes of gene network topologies capable of pattern transformation (Salazar-Ciudad, 2000). However, this previous study did not show why the identified topology classes are the only possible ones. In addition, due to its purely numerical approach, this previous study failed to identify one class of gene network topologies capable of non-trivial pattern formation. Here we take a more general analytical mathematical approach to identify and characterize all possible classes, regardless of the number of gene products they entail.

It is worth noting that there is an abundant theoretical literature on the topic of gene networks in cell biology (e.g., metabolism, gene regulation for basic cellular functions, etc.). However, the topic of this article is fundamentally different from those. Here we are not dealing with gene networks within a single cell, but with systems of cells with identical gene networks that are coupled through extracellular signaling. Specifically, we are interested in those genes networks leading to non-trivial pattern transformation (which is an inherently spatial question).

In this study, we consider three broad types of initial patterns (i.e., initial conditions): homogeneous-with-noise initial patterns, spike initial patterns and combined spike-homogeneous initial patterns. In a homogeneous-with-noise initial pattern, some gene products have zero concentration everywhere while others have the same non-zero concentration everywhere except for some small random fluctuations due to noise at the molecular level. In a spike initial pattern, the concentration of all gene products is zero everywhere except in a region (i.e. the spike region made of a small number of cells) where some gene products have the same non-zero concentration. In the combined spike-homogeneous initial pattern, some gene products have zero concentration everywhere while others have the same non-zero concentration everywhere except in the spike, where the concentration is larger (see Fig. 2). In the discussion we also consider spike initial patterns with small random fluctuations occurring homogeneously over space. Any other initial pattern can be constructed by combining spikes of different heights (i.e. different gene product concentrations) at different positions. We then study how gene product interactions can be wired into networks with different topologies to transform these initial patterns into other (i.e. resulting patterns) non-trivial ones.

**Figure 2.**
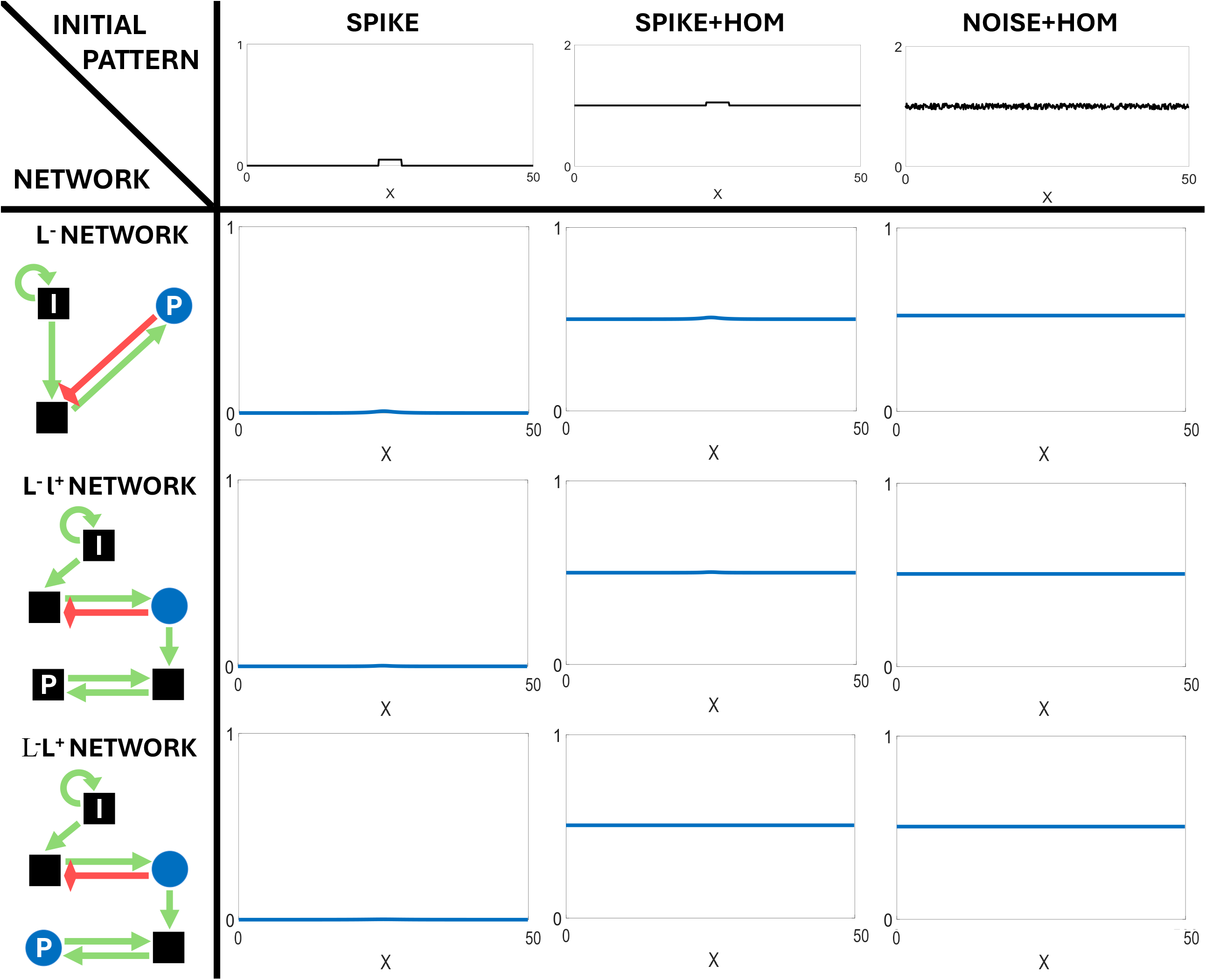
The model transforms initial patterns (A), into resulting patterns (C), through a set of equations implementing gene networks with extracellular signaling (B). The article considers three initial patterns **(A)**: spike initial pattern (left), combined spike-homogeneous initial patterns (middle); and homogeneous-with-noise initial patterns, with small white noise (right). **(B)** Diagram of an example gene network. Black squares represent intracellular gene products, blue circles extracellular signals. Green arrows stand for positive regulations, while red arrows for negative regulations. Weights of the network are given by the *J* matrix; while its topology by the *T* matrix; *D* represents the diffusion coefficients and *M* the degradation rates (see equation (1)). **(C)** Example resulting patterns for each initial pattern in **(A)** under the gene network in **(B)**. For each initial pattern we draw the most possible complex resulting pattern given the gene network topology in **(B)**.

It is worth noting that these three basic initial patterns correspond to spatially discontinuous functions: in homogeneous-with-noise initial patterns, white noise is discontinuous by definition; in spike and combined spike-homogeneous initial patterns, there is a concentration discontinuity between cells on the edge of the spike and nearby cells outside the spike. However, once extracellular signal diffusion begins, these sharp boundaries are smoothed into differentiable gradients, where critical points can be properly defined (e.g., at the center of the initial spike).

## The model

This article considers a set of *N*_*c*_ cells, each with an identical gene network of *N*_*g*_ gene products. For the purpose of clarity we explain our results as if cells have simple spatial arrangements (e.g., a line in 1D, a square lattice in 2D, etc.) and we later discuss them for more complex arrangements of cells in space. We refer to such spatial arrangement as the domain of our system. We consider two types of gene products: intracellular gene products and extracellular signals. The extracellular signals are gene products that are secreted by cells and diffuse in the extracellular space while intracellular gene products are gene products that are not. Even though we only use the general term gene product, our model applies, in principle, to any molecule whose concentration changes as a consequence of the concentration of other molecules in cells.

We consider that extracellular signals diffuse in the extracellular space immediately apical to cells and, therefore, that diffusion occurs in a space with the same spatial dimension than the arrangement of cells. In a 2D lattice of cells, for example, we consider that diffusion occurs in a 2D extracellular space immediately apical to cells. We do not consider any spatial information at scales smaller than a cell (i.e. cells are considered as points). Thus, we treat the extracellular space as an arrangement of contiguous regions, one per cell, and a single concentration value per gene product and region (i.e. concentration variations within each region are not considered, see Fig.3). We consider systems made of many small cells in which diffusion can be approximated by classical continuous equations.

**Figure 3.**
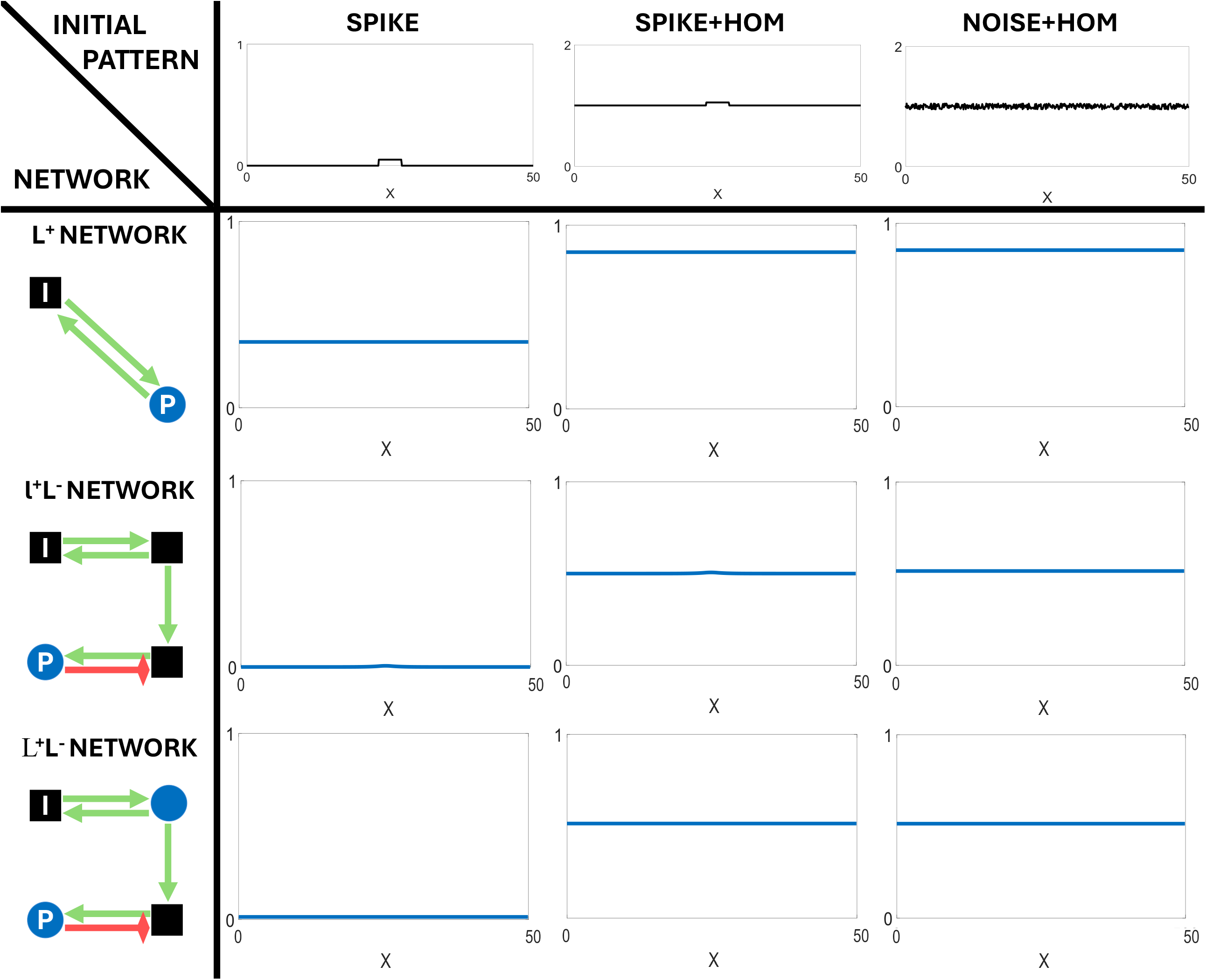
Schema of the elements and spatial relationships in the model. Gray squares are cells. Each white square represents the extracellular space just apical of each cell. Small black squares represent intracellular gene products. Blue circles represent extracellular diffusible signals. Dashed lines represent the diffusion of the extracellular signals over extracellular space. Full green lines represent positive regulation between gene products. Red full lines represent inhibitory regulation between gene products.

For intracellular gene products, *g*_*i*_ (*t, x*) denotes the concentration of gene product *i* inside cell at position *x* at time *t* ≥ 0. For extracellular signals, *g*_*i*_ (*t, x*) denotes the concentration of gene product *i* (i.e., signal) in the region of the extracellular space immediately apical to the cell at position *x*.

The dynamics of gene product concentrations in each cell in our model obey the following system of *N*_*g*_ reaction-diffusion equations (Murray 2002),

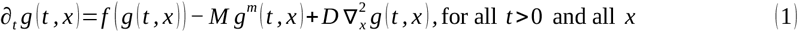

With initial pattern (i.e. initial condition),

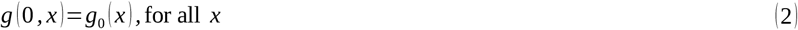

And boundary conditions,

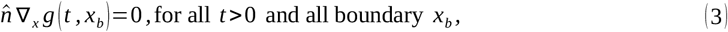

where ∂_*t*_ is the time derivative ; *x* is space in (*n*=1, 2, 3 dimensions) ; 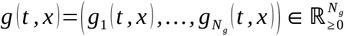 is the vector of gene product concentrations (from gene 1 to *N*_*g*_) at the cell at time *t* and position 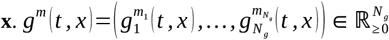 is a vector of powers of the gene product concentrations and ***m*** is a vector of positive integer exponents; 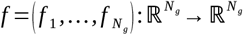 governs the interactions between gene products and determines how much of each gene product *i* is being synthesized in a given instant of time. *f* is, thus, a function that takes a vector of concentrations as input and returns another vector of concentrations as output ; ∇^2^ denotes the Laplace operator (i.e., 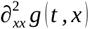) in 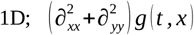 in 2D; in 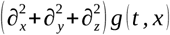in 3D). 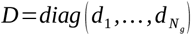 is an *N*_*g*_ *× N*_*g*_-diagonal matrix with real positive diffusion coefficients for each gene product along its diagonal. By definition extracellular signals have *d*_*i*_ >0 while for intracellular gene products *d*_*i*_=0. Similarly, 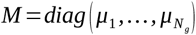 is a *N*_*g*_ *× N*_*g*_-diagonal matrix with the real degradation coefficients for each gene product along its diagonal, *μ*_*i*_ ≥ 0 . The degradation coefficient of a gene product is the default rate at which it would be degraded by proteases if no other factors affect its degradation. Notice that equation (1) has *N*_*g*_ components and applies to each cell. For the cases of extracellular signals, i.e. their concentration in a cell can affect that in distant cells through the diffusion term (last term in (1)).

In (2), *g*_0_ (*x*) represents the initial pattern (i.e. initial spatial condition). In (3), 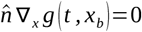 are the boundary conditions of the system (zero flux boundary conditions). ∇ denotes the nabla operator (i.e., ∂_*x*_ *g* (*t, x*) in 1D; (∂_*x*_ +∂_*y*_) *g* (*t, x*) in 2D; and (∂_*x*_ +∂_*y*_ +∂_*z*_) *g* (*t, x*) in 3D.); and 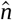 is a unit vector normal to the boundary of the cell arrangement.

The last term in (1) is Fick’s second law of diffusion (Fick, 19855) describing the contribution of extracellular diffusion to changes in the concentration of extracellular signals. Our model does not consider membrane-bound signals (see Salazar-Ciudad *et al*., 2000 for a similar study that does).

The first two terms in system (1) can be understood as a reaction term. *f* defines a directed graph via its Jacobian matrix ***J*** (i.e., the matrix of first derivatives of *f* with respect to each gene product concentration), where each node stands for a given gene product, and each edge describes an interaction between gene products (Fig.2B).

Following the standard terminology in molecular biology, we say that a gene is being expressed in a cell at a give time if its gene product is being produced by that cell (i.e. the first term of (1) is larger than zero for that cell at that time). Notice that, because of the diffusion term, extracellular signals can have positive non-zero concentration around cells that are not expressing them.

We say that gene product *j* directly regulates gene product *k* if the corresponding element of the Jacobian matrix of *f* is non-zero (i.e., *J*_*kj*_ ≠ 0). This regulation is positive if *J*_*kj*_>0 (i.e., *g* _*j*_ increases with *g*_*k*_); or negative if *J*_*kj*_<0 (i.e., *g* _*j*_ decreases with *g*_*k*_). Similarly, we say that *k* is downstream of *j* if there exists a chain of regulations between gene products going from *j* to *k*. Correspondingly, *j* is said to be upstream from *k*. Notice that a gene product can be both upstream and downstream of a set of gene products, thus forming a regulatory loop. We call extracellular loops those regulatory loops in which at least one gene product is an extracellular signal.

Our results are only valid for functions *f* and reactions terms (i.e., *f* (*g*) − *M g*^*m*^) that satisfy some specific requirements. These are very broad biological requirements that are likely to be fulfilled by many developmental gene networks.

### Requirements on the reaction term and f

**R1**. *f* is continuous and continuously differentiable (at least locally), so that its Jacobian matrix can be properly defined. Within each cell, we assume that the number of copies of each gene product and its rate of production are large enough for its concentration to be treated as continuous, as in many other models of pattern formation (Salazar-Ciudad *et al*., 2000, 2001; Cotterel and Sharpe, 2010; Marcon *et al*., 2016; Zheng *et al*., 2016; Jimenez *et al*., 2017; Diego *et al*., 2018 ; Leyshon *et al*., 2021). Notice that the degradation term in (1) is also continuous and continuously differentiable, − *M g*^*m*^ (*t, x*).

**R2**. *f* is such that *f* (*g* (*t, x*)) − *M g*^*m*^ (*t, x*) is non-linear in at least one of its components. In other words, at least for one gene product, the dependence of concentration on that of other gene products (or its own) is not linear. This is because it is well known that no non-trivial pattern transformation is possible in purely linear systems (Kondepudi & Prigogine, 2014; Murray, 2002).

**R3**. *f* is an explicit function of ***g***, but not an explicit function of time or any time derivative of ***g***. This means that the regulation of a gene product at a given moment depends only on the concentration of other gene products at that given moment, and not explicitly on past concentrations or their rate of change over time.

**R4**. *f* is such that ***g*** is always non-negative and bounded. This is because ***g*** is a vector of concentrations and concentrations cannot be negative. Similarly, cells are finite and, hence, cannot produce an infinite amount of gene products. Thus, *f* should be such that gene product concentrations never go to infinity.

**R5**. *f* is monotonously increasing or decreasing with respect to each gene product concentration. In other words, the sign of gene products interactions cannot change with the concentration of the interacting gene products (when everything else is kept constant). Thus, if a gene product *k* activates another gene product *j*, then *k* is an activator of *j*, even if its concentration changes. The same applies to inhibitory interactions (i.e. the sign of the regulation of *k* by *j* does not change with the concentration of *j*). In other words, the partial derivative of *f* with respect to each gene product *j*, should have the same sign for any value of *g*_*j*_,

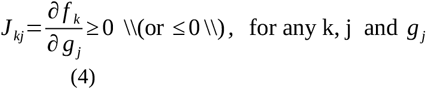

where *f*_*k*_ is the output of *f* regarding gene product *k* (i.e., the k-th component of *f*), and all gene product concentrations, except for *g* _*j*_, are kept constant for the evaluation of *J*_*kj*_. There are some known examples of particular gene product interactions that do not fulfill this requirement (Nelson *et al*., 2021) but, in general, the sign of the regulation of a gene product by another does not change in any complex way with the concentration of either gene product. Even in the reported cases where this happens, sign changes only occur once or twice (Nelson *et al*., 2021). Requirement R5 is simply a way to consider that pattern formation does not just arise from very simple gene networks with very complex interactions between each pair of genes (i.e. changes in the sign of the regulation depending on the concentration of the regulator) but from networks of relatively simple interactions between gene products.

Through this article when we refer to the “the parameters” we mean the parameters described in (1) (i.e. its parameters, ***M, D*** and ***J***) and to any additional parameters a specific *f* may have. We consider that these parameters do not vary over time or space.

In this article we define the topology matrix *T* of a gene network as the matrix whose elements are − 1, 1 or 0 depending on the sign of the corresponding entry in the Jacobian matrix (i.e., *T* =*sgn* (*J*)) (see Fig.2B). The topology of a given gene network is then a description of which gene products interact with which gene products and the sign of that interaction. In contrast, from now on, we will reserve the term gene network for a gene network topology with a specific ***f*** and all its parameter values so that a specific resulting pattern can arise from a specific initial pattern (i.e. a full implementation of equations (1)-(3) for specific ***f*** and specific values of the parameters). In this article a topological class is a set of topologies that fulfill some requirement definable at the level of topology, e.g. they have at least one positive loop. Studying gene network topology is important because for most developmental systems we only have information about *T*, and not so much about ***f*** (Gilbert and Barresi, 2023).

The results are organized into a set of logical arguments (for the least evident of them, we provide mathematical proofs in the Supporting Information). First we present some trivial requirements that the topology of all pattern-transforming gene networks need to fulfill. Secondly, we explain why all gene networks leading to non-trivial pattern transformations can be classified into three topological classes, and their combinations. Thirdly, we explain how the topological class to which a gene network belongs determines which non-trivial pattern transformations it can produce and which ones it cannot produce. In other words, belonging to one of these three classes is a necessary, but not sufficient, condition for a gene network to be able to lead to non-trivial pattern transformations. Among the types of pattern transformations possible from its topological class, a given gene network would produce one or another depending on the exact form of ***f*** and the value of the parameters. However, the topological class to which a gene network belongs indicates which non-trivial pattern transformations it cannot produce, regardless of its parameters and ***f***.

## Results

Any network with more than two gene products can be partitioned into a set of subnetworks but, as we will see, partitioning gene networks based on extracellular signals is especially useful. In this article, we call a *signal subnetwork* to the set of interactions between all gene products downstream of an extracellular signal (including the signal itself). Signal subnetworks, or from now on simply subnetworks, can partially overlap since a gene product can be downstream of several extracellular signals (see Fig.4). An extracellular signal can be upstream of other extracellular signals and, thus, a signal subnetwork can include other signal subnetworks (see Fig. 4).

**Figure 4.**
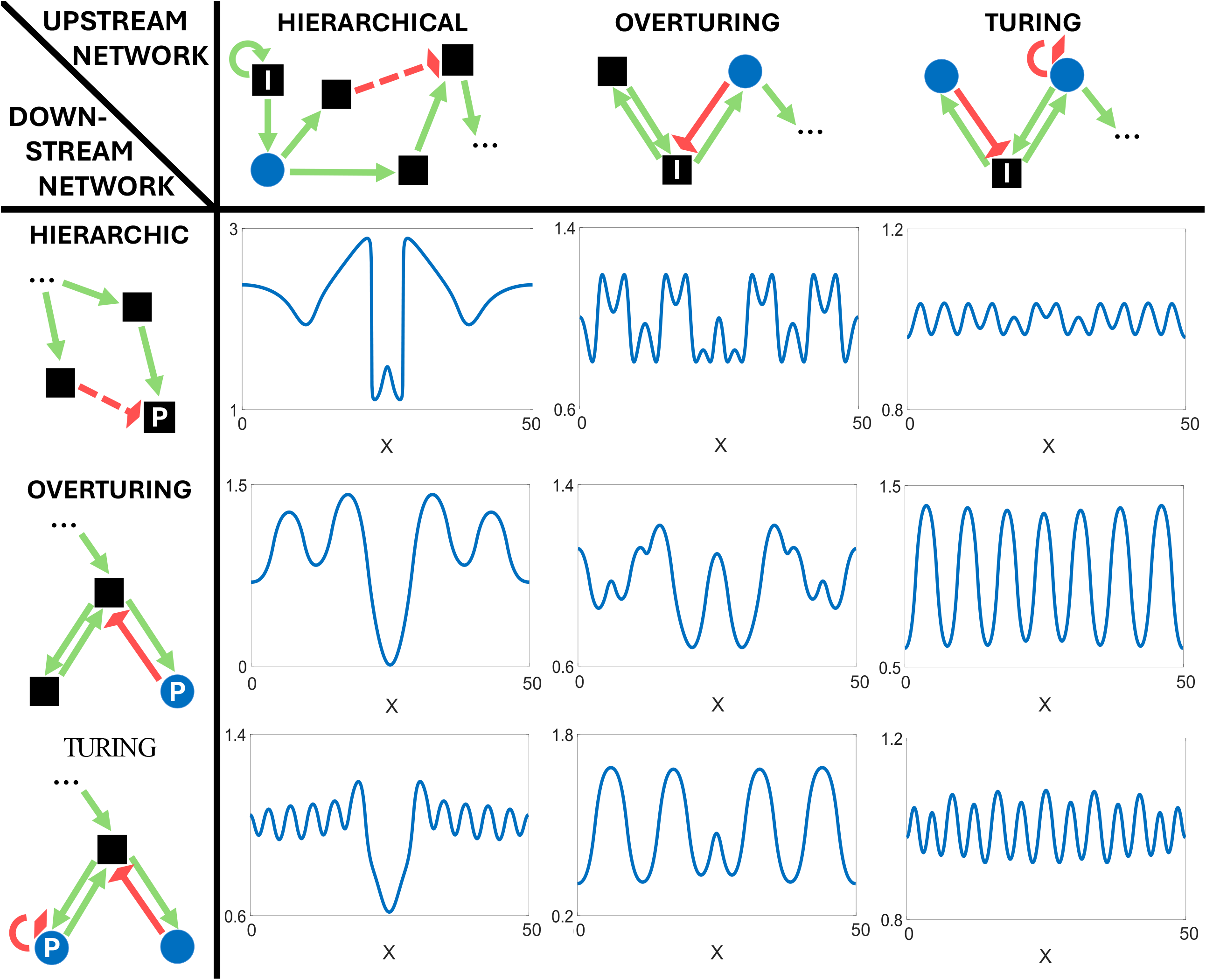
Definition of chains, loops, and signal subnetworks. **(A-D)** Parts of intracellular gene networks **(E-F)** Schema of simple extracellular regulatory loops. **(G-J)** Examples of gene regulatory networks with their signal subnetworks depicted. Small black squares represent intracellular gene products, and blue circles represent extracellular diffusible signals. Green arrows denote positive regulation, and red blunt arrows denote inhibition. Dots (…) indicate any positive chain of gene product interactions. **(A)** A pPositive regulatory chain is a sequence of interactions in which each gene product positively regulates the next and is positively regulated by the previous one in the sequence. A sequence with an even number of inhibitory interactions is also a positive regulatory chain, provided that each inhibited gene product receives a positive input from elsewhere. **(B)** Negative regulatory chain: as in **(A)** but with an odd number of inhibitory interactions. **(C)** A positive intracellular loop is a positive regulatory chain whose first and last gene products coincide and in which all gene products are intracellular. **(D)** Negative intracellular loop: as in **(C)** but with an odd number of inhibitory interactions. **(E)** A positive extracellular loop is a positive regulatory loop in which at least one gene product is an extracellular signal. **(F)** Negative extracellular loop: as in **(E)** but with an odd number of inhibitory interactions. **(G)** An example network in which we have surrounded its signal subnetwork with a grey rectangle. **(H)** Example network with two signal subnetworks. **(I)** Example network with three extracellular signals; each signal subnetwork is enclosed by a rectangle. Note that since signal 3 and 4 are in a loop their signal subnetworks include the same gene products. **(J)** Another example network with its subnetworks indicated.

### Basic requirements on gene networks capable of pattern transformation

In this section we present some simple requirements on the topology of any gene network capable of pattern transformation. These are rather trivial but introducing them here facilitates explaining later results.

**I1**. Any gene network capable of non-trivial pattern transformation must include at least one signal subnetwork whose extracellular signal is either present in the initial pattern or positively downstream of some gene product that is. By ‘present in the initial pattern’, we mean that the corresponding gene product has a non-zero initial concentration somewhere in the domain (i.e., its gene is expressed in the initial pattern). Since we do not consider cell movement, mechano-transduction, and membrane-tethered signaling, non-trivial pattern transformations can only occur through the secretion of extracellular signals, which necessarily requires the activation of at least one signal subnetwork.

**I2**. All gene products are being degraded at all times (second term in (1)). Thus, for a gene product to be present in the resulting pattern, it should receive positive regulation from some gene products. This ultimately requires that all the gene products present in the resulting pattern are either within a self-activatory loop or downstream of it (otherwise, their concentration will decay to zero over time). This broad case includes genes that are constitutively expressed, meaning that they are always produced by cells regardless of other gene products (this can be viewed as these genes being within a self-activatory loop of their own or downstream of one).

Considering I1 and I2, it follows that all gene networks capable of non-trivial pattern transformation should contain a positive regulatory loop that is upstream of the patterned gene products and downstream of gene products present in the initial pattern. These loops can be extracellular or intracellular (i.e., include extracellular signals or not).

### Gene network classification

In this section we explain that all conceivable signal subnetworks can be classified into just three classes based on how cells respond to extracellular signals in terms of secreting, or not, the same extracellular signals (see Fig.4). Let *A* be the most upstream extracellular signal of a given signal subnetwork. Then, there are trivially only two options, either *A* is downstream of itself (directly or indirectly), or it is not. In the latter case, we say that the subnetwork is hierarchical, or class H, while in the former case we say that the subnetwork is emergent (Salazar-Ciudad *et al*., 2000). Notice that this exhausts all possibilities: an extracellular signal is either downstream of itself or not, there are no other possibilities. Thus, all signal subnetworks can be classified as hierarchical, emergent or a combination of them (for subnetworks composed of other subnetworks).

The emergent class can be further divided into the L^+^ class, if *A* is positively downstream (and upstream) of itself, or into the class L^-^, if *A* is negatively downstream (and upstream) of itself. This gives three topological classes: H, L^-^ and L^+^.

H subnetworks can contain regulatory loops as long as these do not include an extracellular signal (because in this case the loop would be extracellular, and the subnetwork, emergent). In fact, because of I2, any H network leading to pattern transformation must contain a positive intracellular loop (we label these as l^+^).

A gene network topology can be composed of several signal subnetworks. We classify whole gene networks topologies in the same way as their composing subnetworks. For example, a gene network topology with only H subnetworks is an H gene network, while a gene network topology with only L^+^ subnetworks is an L^+^ network. Composite gene networks topologies are labeled according to their composing subnetworks, independently of the number of subnetworks of each class they contain (see Fig.5). For example, an H L^+^ gene network can contain one, or more, of H subnetworks, and one, or more, L^+^ subnetworks.

**Figure 5.**
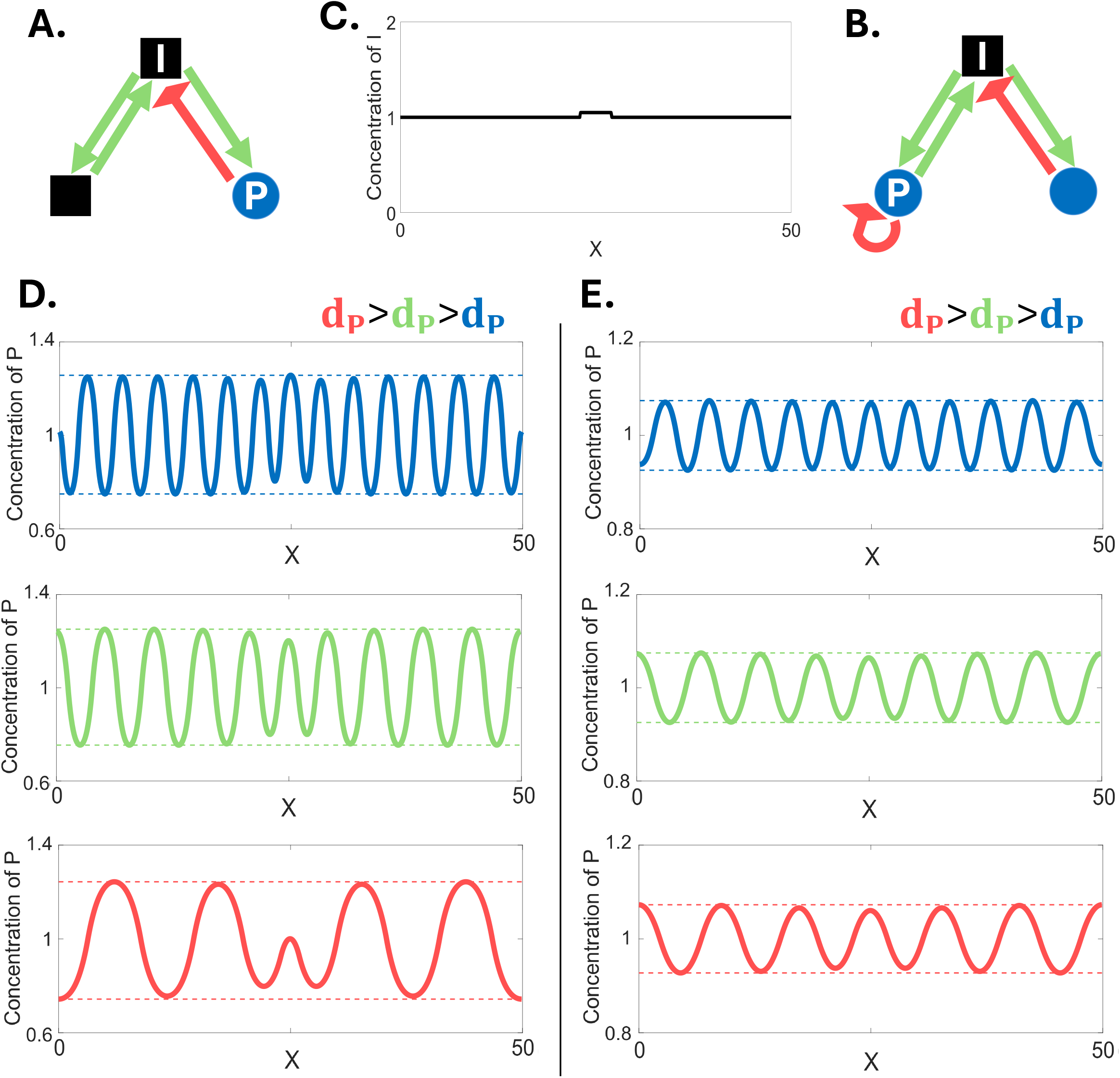
Subnetwork and network classification. **(A)** Schema representing each class of subnetwork. In a H signal subnetworks, the extracellular signal is not downstream of itself. In a signal L^+^ subnetworks, the extracellular signal is both upstream and downstream of itself, forming a positive extracellular loop (which may include any number of gene products). In a L^−^ signal subnetwork, the extracellular signal is negatively upstream and downstream of itself. The box labeled ‘rest of subnetwork’ represents any gene network provided that the most upstream extracellular signal is not negatively upstream of itself (i.e., no negative loop leads back to it). **(B) (B)** Example of a H gene network containing two H subnetworks. **(C)** Two examples of L^-^ gene networks, one with on L^-^ subnetwork (left) and one with two L^-^ subnetworks (right). **(D)** Example of a L^+^ gene network with two L^+^ subnetworks. **(E)** Two examples of L^+^L^-^ gene networks. Each of them has a L^+^ subnetwork and a L^-^ subnetwork. **(F)** Example of a H gene network. **(G)** Example of a gene network with a L^+^ subnetwork, a L^-^ subnetwork and a H subnetwork. Colors of arrows and gene products as in figure 2 and 4.

Our classification into classes H, L^+^ and L^-^ exhausts all possible topologies for gene networks. Within each topological class, however, there are gene networks that can lead to non-trivial pattern transformations and gene networks that cannot (e.g. depending on the parameters and the exact ***f***). In the rest of this article we first identify further topological requirements that gene networks of each class must satisfy in order to lead to non-trivial pattern transformations. Then, we study the types of resulting patterns possible from each topological class. These types are described in very broad terms. In other words, we do not specify which exact resulting patterns arise from each gene network but some general features and commonalities among the resulting patterns possible from the gene networks within each class. We do the same for the gene networks combining different classes of subnetworks.

#### Linear stability analysis

In this section, we introduce the mathematical apparatus that allows us to link the topology of a gene network to its ability to produce non-trivial pattern transformation: the dispersion relation. We will treat each initial pattern as a perturbation of an otherwise spatially homogeneous steady state, and we will study the conditions under which such perturbation grows in time. If initial perturbations do not grow, one recovers the original homogeneous steady state and, thus, there is no pattern transformation. Even when these perturbations grow, one may recover a different homogeneous steady state as a resulting pattern, i.e. no non-trivial pattern transformation either. Thus, instability against perturbations is a necessary but no sufficient condition for pattern transformation.

To analyze whether a perturbation will grow we will use the standard linear stability analysis of the dynamical system defined by our model equations (1)-(3). In this study we focus in those ***f*** functions for which equation (1) to (3) admit a steady state homogeneous solution *g* (*t, x*) ≡ *g* * (i.e. a homogeneous steady state) given by *f* (*g* *) − *M* (*g* *)^*m*^=0. Hence, let 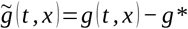 denote a small concentration perturbation from the homogeneous steady state *g* *. Then, plugging 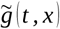 back into (1)-(3) shows that the evolution of such perturbation in space and time is given by the linearized reaction-diffusion equations,

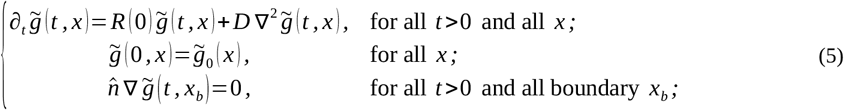

where 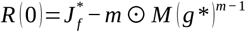 is the reaction-diffusion matrix in the absence of diffusion; and 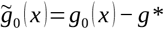 represents the initial perturbation. In the reaction diffusion matrix, 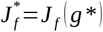 is the Jacobian matrix of *f* (i.e. the matrix of derivatives of *f* (*g*) with respect to each gene product) evaluated at the steady state *g* *, and 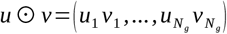 denotes component-by-component vector multiplication.

Each initial pattern can be described as a different small initial perturbation 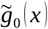 around an otherwise homogeneous steady state *g* *. In homogeneous-with-noise initial patterns, *g* *>0 and 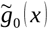 is small white noise. In spike initial patterns, *g* *=0 and 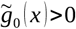 only for positions *x* such that ||*x* − *x*_*c*_||< *L*, where *x*_*c*_ is the center of the spike and *L* its width. In the combined spike-homogeneous initial pattern, *g* *>0 and, again, 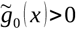 for positions *x* in the spike such that ||*x* − *x*_*c*_||< *L*. In both cases, the initial concentration in the spike must be small enough for linearization (5) to hold (see the appendix for the case of larger spikes).

Over time, the initial perturbation can either decay (i.e., 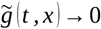 as *t* →∞) or grow away from the original homogeneous steady state. In the former case, we say that 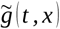 is a stable perturbation; in the latter, that 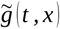 is an unstable perturbation. Since we are interested in the gene networks that can lead to pattern transformations through extracellular signaling, we are interested in the gene networks in which perturbations are unstable when there is extracellular signal diffusion (i.e., when some elements of ***D*** are non-zero).

Solutions to (5) can be found using the spectral decomposition of the Laplacian ∇^2^ with zero-flux boundary conditions (Murray, 2003). In this sense, let *W* _*k*_ (*x*) be a zero-flux eigenmode of the Laplacian operator in our domain, that is, a non-trivial solution (i.e., *W* _*k*_ (*x*) ≠ 0 for at least one *x*) of the following equations,

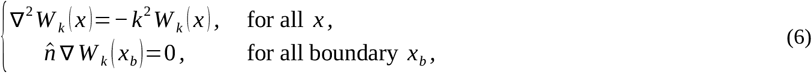

for some *k* ∈ℝ^*n*^, with *n*=1, 2 or 3. We denote by *σ* (∇^2^) the countable set of all *k* ∈ℝ^*n*^ for which equation (6) admits a non-trivial solution (Baker *et al*., 2008), and we say that *k* ∈*σ* (∇^2^) are the wavenumbers of the corresponding eigenmode *W* _*k*_ (*x*). This is a slight abuse of notation based on the fact that, in simpler spatial domains (e.g., a 2D square), eigenmodes *W* _*k*_ (*x*) are trigonometric functions for which each entry in *k* counts the number of maxima and minima that *W* _*k*_ (*x*) displays along each spatial dimension.

Eigenmodes *W* _*k*_ (*x*) can be seen as a generalization of classical Fourier modes and hence, we can now consider the series expansion of 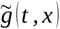 in terms of *W* _*k*_ (*x*), that is,

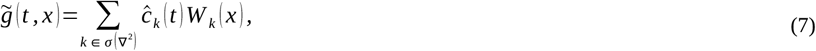

where each 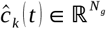 is given by,

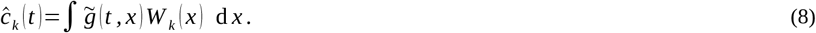

If we now plug series expansion (7) into system (5) and use the linearity of the equations to split the sum into terms for each *k*, we get,

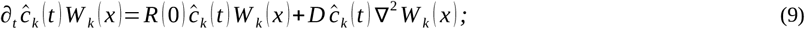

and then, if we use (6) to solve the space derivatives, and eliminate alike terms from each side of the equations (i.e., *W* _*k*_ (*x*)), we get that the time evolution of each *c*^ _*k*_ (*t*) is given by the equation,

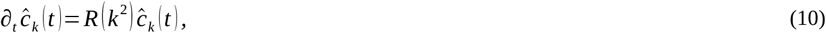

where *R* (*k*^2^)=*R* (0)+*k*^2^ *D*=*J*_*f*_ (*g* *) −*m*⊙*M* (*g* *)^*m*−1^ is the so-called reaction-diffusion matrix. Solutions to equation (10) are given by,

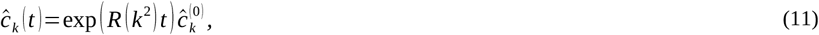

where exp (*R* (*k*^2^) *t*) is a matrix exponential; and each 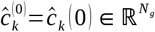 is computed as in (8), changing 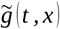 by the corresponding initial perturbation 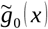.

At this point, we can plug (11) back into (7) and thus, we get that solutions 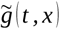 to the linearized reaction-diffusion equations (5) can be written as a series expansion of the form,

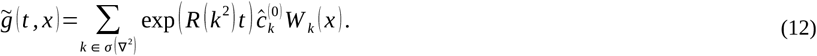

However, for our linear stability analysis, we are only interested in the long-term behavior of (12), namely whether 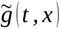 grows over time or eventually decays back to 0. In this sense, the long-term behavior of exp (*R* (*k*^2^) *t*) is dominated by the eigenvalue of *R* (*k*^2^) with the greatest real part and then, we can reduce expression (12) to,

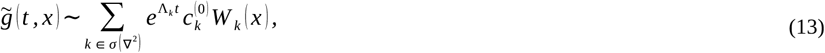

where Λ_*k*_ ∈ℂ is the eigenvalue of *R* (*k*^2^) with the greatest real part, and 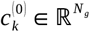 is the spectral projection of 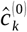 onto the corresponding eigenspace with eigenvalue Λ_*k*_ .

The different eigenvalues of *R* (*k*^2^) vary with *k*, and this variation is given by the so-called dispersion relation. By definition, the dispersion relation corresponds to the characteristic equation of the reaction-diffusion matrix *R* (*k*^2^) (Murray, 2003; Baker *et al*., 2008), that is,

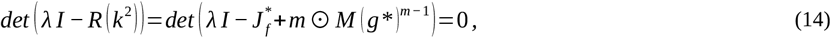

where *I* denotes the *N*_*g*_ *× N*_*g*_ identity matrix. Then, Λ_*k*_ ∈ℂ in the exponent of (13) corresponds to the solution of (14) with the greatest real part for each given *k* (i.e., Λ_*k*_=*λ*_*i*_ such that Re (*λ*_*i*_) ≥ Re (*λ* _*j*_) for any other solution *λ* _*j*_ of (14)). Accordingly, we will refer to Λ_*k*_ as the principal branch of the dispersion relation.

The principal branch in of the dispersion relation, (13), determines whether a perturbation is stable or unstable. If Re (Λ_*k*_)<0 for some *k* ∈*σ* (∇^2^), then 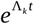 decays to 0 as *t* →∞ and we say that *k* (resp. *W* _*k*_ (*x*)) is a stable wavenumber (resp., eigenmode). Conversely, if Re (Λ_*k*_)>0 for some *k* ∈*σ* (∇^2^), then 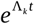 grows as *t* →∞ and we say that *k* is an unstable wavenumber (resp., eigenmode). Hence, perturbation 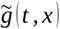 is stable if all wavenumbers *k* ∈*σ* (∇^2^) are stable, and is unstable if there exists at least one unstable wavenumber *k* ∈*σ* (∇^2^) for which the principal branch has a positive real part. Note that, in accordance with requirement R4, the unbounded exponential growth of unstable eigenmodes in (13) will be halted, (13), eventually, by the higher order terms in (1) that we disregard in the linear approximation (6).

Each gene network has a 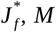 and *D* matrices, and each such set of matrices defines a different reaction-diffusion matrix *R* (*k*^2^). This means that each gene network has and associated dispersion relation that is given by equation (14). In this sense, a gene network can only produce pattern transformations if its associated dispersion relation has at least one unstable wavenumber *k* ∈*σ* (∇^2^) for which Re (Λ_*k*_)>0. Moreover, each initial perturbation 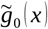 can be described as a series expansion in terms of wavemodes *W* _*k*_ (*x*) (e.g., a homogeneous-with-noise initial pattern will result in a series with many eigenmodes *W* _*k*_ (*x*) and many of them with large wavenumbers *k*). In this sense, each gene network can be seen as leading, or not, to pattern transformation by specifically destabilizing some of these wavenumbers while stabilizing others. The later means that some perturbations, the ones with the destabilized wavenumbers, grow over time until some non-linearities in *f* (<u>R2</u> and R4) may preclude their further growth. This means that the resulting patterns, can be characterized, to a large extent, by these destabilized eigenmodes (Murray, 2003), even though some other wavenumbers may be destabilized by nonlinear effects (e.g., resonant eigenmodes (Castelino *et al*., 2020)).

The principal branch (i.e. Λ_*k*_) allows to classify all the pattern transformations arising from gene networks into two mutually exclusive types: those with a finite amount of unstable wavenumbers and those with an infinite amount of unstable wavenumbers.

We say that a gene network is **RD-unstable of the first kind** if, for some choice of the parameters, its associated dispersion relation yields a finite amount of unstable wavenumbers (see Fig.6A-B). In this case, the resulting patterns, if heterogeneous, are periodic since series expansion (13) reduces to a sum of finitely many unstable eigenmodes, and eigenmodes are trigonometric functions for the vast majority of simple 1, 2 and 3-dimensional domains considered for pattern transformations in development (Baker et al., 2008). Notice that, since none of the initial patterns we consider are periodic, the emergence of these periodic resulting patterns constitutes a non-trivial pattern transformation (i.e. new concentration peaks and valleys will arise in some places).

**Figure 6.**
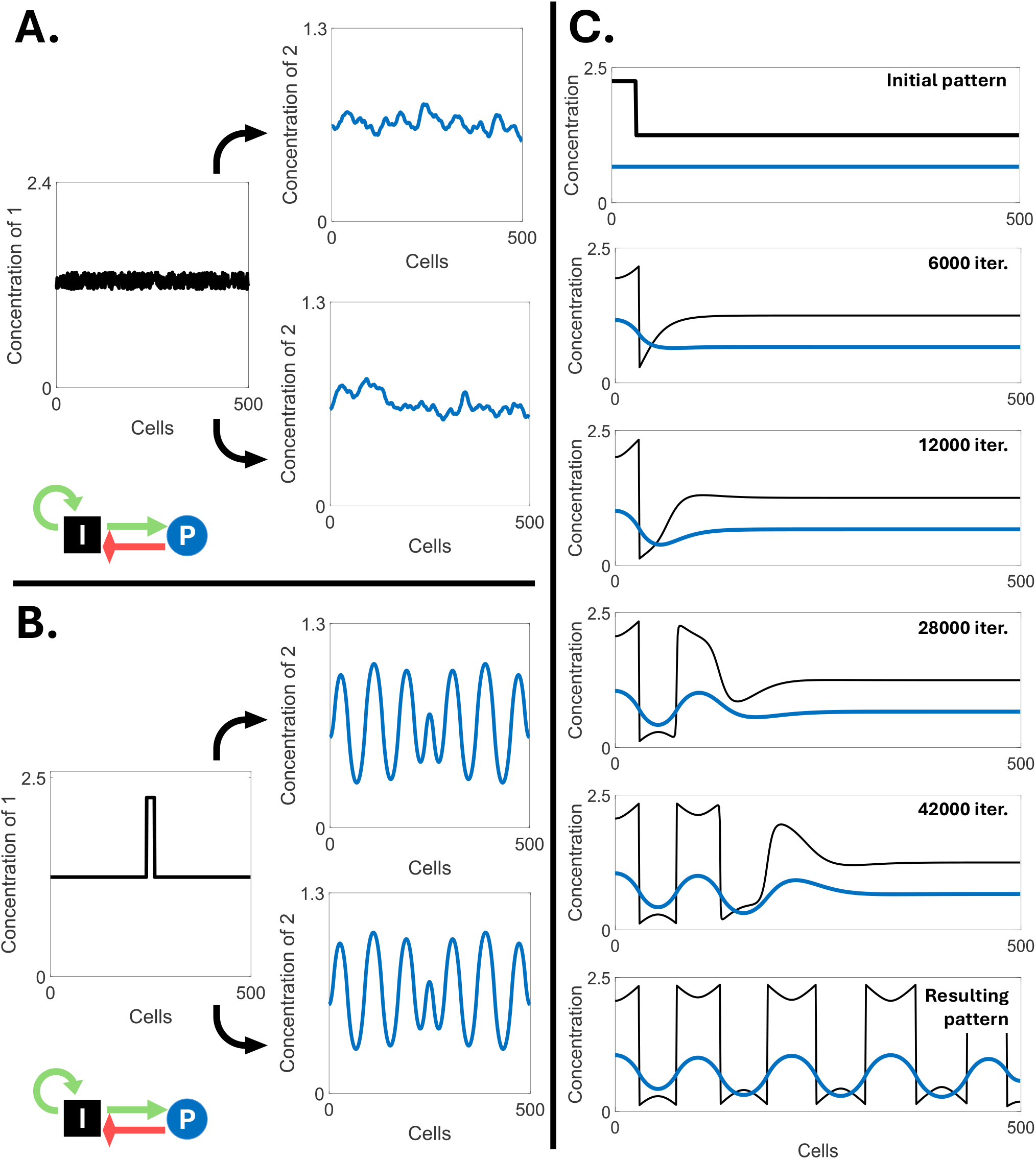
Instances of principal branches of dispersion relations. **(A-B)** Dispersion relations of RD-unstable gene networks of the first kind (i.e. there is only a number of wavenumbers, x-axis, for the which there are eigenvalues with a positive real part). The real part of the principal branch of a dispersion relation can diverge to −∞ for large wavenumbers **(A)**; it can converge to a negative finite value **(B). (C-D)** Dispersion relation of RD-unstable gene networks of the second kind (i.e. there is an infinite number of wavenumbers with eigenvalues with a positive real part). The real part of the principal branch of a dispersion relation cannot diverge to +∞ (Klika *et al*., 2012).

We say that a gene network is **RD-unstable of the second kind** if, for some choice of the parameters, its associated dispersion relation yields an infinite amount of unstable wavenumbers (see Fig.6). Previous research (Klika *et al*., 2012) has shown that the dispersion relation never goes to positive infinity (see Fig.6E) and that gene networks that are RD-unstable of the second kind have a dispersion relation that saturates to a positive value and, thus, have an infinite queue of large unstable wavenumbers (see Fig.6C-D). Since there is an infinite number of unstable wavenumbers, the resulting patterns, if heterogeneous, can have an infinite sum of eigenmodes in (9). There exist both periodic and non periodic functions that admit such an infinite series representations (Stein & Shakarchi, 2003). Nevertheless, as we later explain, these gene networks lead to three broad types of heterogeneous resulting patterns: periodic (but not trigonometric) patterns, noisy patterns, and radially symmetric multi-peaked patterns. Which of these can arise depends on the initial pattern and the topological class of the gene network.

#### Positive regulatory loops determine the kind of RD-instability

For a gene network to be RD-unstable it needs to have one or more positive regulatory loops. If such loops are all extracellular, then the gene network can only be RD-unstable of the first kind or RD-stable ; and if such loops are all intracellular, then the gene network can only be RD-unstable of the second kind or RD-stable. In here, we provide a proof of these two statements for the particular case of gene networks with only one positive loop and linear degradation, while in section S2 of the appendix we give a general prove for gene networks with any number of loops.

In the square matrix 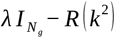 in (13), each off-diagonal entry corresponds to the interaction between two different gene products, while each diagonal entry contains the variable *λ*, the degradation rate of a given gene product, its diffusion coefficient and, perhaps, some self-regulation term (i.e., *J* _*ii*_ ≠ 0). By the Leibniz formula for determinants (Strang, 2016), each term in 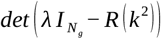 (i.e., the characteristic polynomial of the reaction-diffusion matrix) is the product of *N*_*g*_ entries of different rows and columns (i.e. different permutations). This means that each term of the characteristic polynomial of *R* (*k*^2^) contains only one entry per gene product (i.e., row or column) and hence, only the terms that correspond to regulatory loops are non-zero (since all other terms contain at least one zero element in the product), except for the product of all diagonal entries. The characteristic polynomial reduces, thus, to a sum of loop terms. This implies that, if a gene network contains no loops, the characteristic polynomial of *R* (*k*^2^) has just one term: the product of the elements of its diagonal. Namely, the dispersion relation of the network reads,

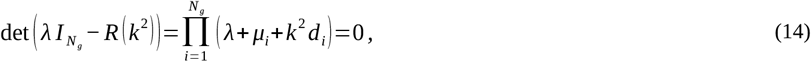

and, thus, its eigenvalues are given by,

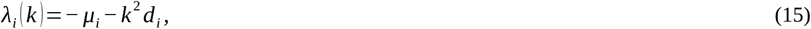

for all *i*=1, …, *N*_*g*_. In this case, all eigenvalues are negative and, thus, all wavenumbers are stable: no pattern transformation is therefore possible. This proves that, without regulatory loops in the network, pattern transformations are not possible.

Let us now consider the case of a gene network with only one loop, which we assume to contain *N*_*l*_>1 gene products. Without loss of generality, gene products can be relabeled so that the submatrix of 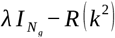 of the loop contains all non-zero elements under the diagonal and in the top-right entry. Namely,

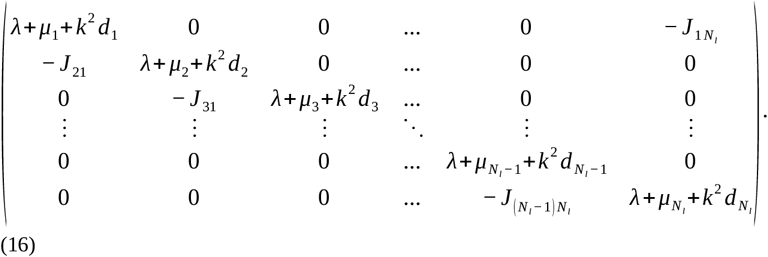

Similarly, given that there exists no other loop in the network, we can relabel the remaining gene products so that the rest of the matrix is upper triangular (Bang-Jensen, 2008). Thus, the characteristic polynomial has two terms: the product of all diagonal entries, and the product of the non-diagonal entries in the loop times the remaining *N*_*g*_ − *N*_*l*_ diagonal entries. The corresponding dispersion relation reads,

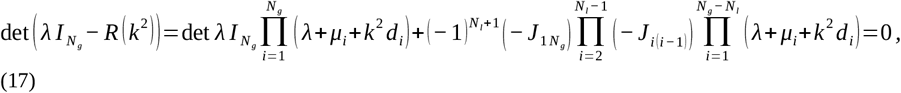

where the minus sign before each *J*_*ij*_ comes from (13) and the factor 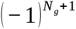 comes from the Leibniz formula for the case of a loop like (16). A (− 1) factor can be extracted from each Jacobian entry to obtain a 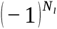 factor that then simplifies (17) to,

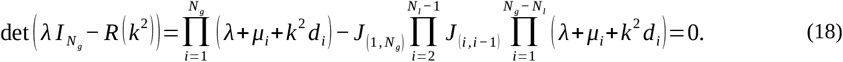

Similarly, the factor (*λ*+ *μ*_i_ +*k*^2^ *d*_*i*_) is found *N*_*g*_ times in the left term and *N*_*g*_ − *N*_*l*_ times in the right term and so, (18) can be further simplified into,

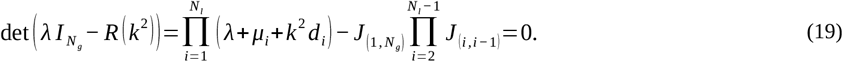

All terms in the first product of (19) are positive and, thus, its expansion is an *N*_*g*_- polynomial in *λ* with positive coefficients,

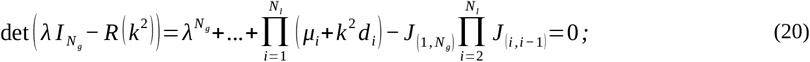

and then, the characteristic polynomial has a negative constant term if:

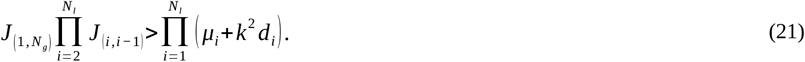

Notice condition (21) only holds if the regulatory loop is positive (i.e. all regulations are positive or there is an even number of negative ones). If condition (21) holds for some ***k***, it follows that the characteristic polynomial takes a negative value for *λ*=0. Moreover, since the term with the largest degree is positive, the characteristic polynomial becomes positive as *λ*→∞. This implies that the polynomial has to cross the *λ*-axis at least at one positive *λ* and such positive root is a positive eigenvalue of *R* (*k*^2^) for any *k* satisfying (21). In other words, we have proven that the dispersion relation of the gene network with a positive regulatory loop can yield unstable wavenumbers and, thus, the network can be RD-unstable depending on the parameters.

If the positive regulatory loop is extracellular, then at least one of the diffusion coefficients of its gene products is non-zero (i.e., *d*_*i*_ ≠ 0 for some *i* in (21)). Then, given any specific choice of the parameters (i.e., specific values for *J*_*ij*_) condition (21) will not hold for large wavenumbers *k* because the right-hand side simply grows unbounded as *k*^2^ →∞. This means that condition (21) can only hold, at most, for a finite amount of small wavenumbers *k*. Thus, gene networks that have a single positive loop and in which this loop is extracellular, can only be RD-unstable of the first kind, or not RD-unstable at all. A general proof for the case of gene networks with several positive extracellular loops, and in fact for any gene network topology, is given in section S2 of the appendix.

If the positive loop is intracellular, then the diffusion coefficients of all its gene products are zero (i.e., *d*_*i*_=0 for all *i* in (21)). Consequently, condition (21) reduces to,

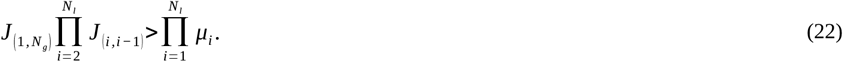

In this case, the existence of positive eigenvalues does not depend on any specific wavenumber *k*. This means that, whenever the parameters allow (22) to hold, all wavenumbers *k* are unstable (i.e., there are infinitely many unstable wavenumbers). In other words, gene network that have a single intracellular positive loop can only be RD-unstable of the second kind, or not RD-unstable. Again, a general proof for the case of gene networks with several positive intracellular loops is given in section S2 of the appendix.

As we have seen, at least one positive regulatory loop is a necessary for a gene network to lead to pattern transformation. As we will see, for non-trivial pattern transformations to occur, negative regulation is also required (otherwise, the resulting pattern can only be homogeneous). Given that positive loops are required, and that there are only three classes of signal subnetworks, it follows that the gene networks leading to pattern transformation can only be: gene networks with positive intracellular loops and intracellular negative regulation (i.e., H networks); gene networks with positive intracellular loops and negative extracellular loops (i.e., *L*^*-*^ gene networks with positive intracellular loops l^+^); gene networks with extracellular positive loops and extracellular negative loops (i.e., emergent L^+^L^-^ gene networks); or a combination of any of the above. As we will see, each of these classes of gene networks leads to qualitatively different types of resulting patterns and the dispersion relation, together with some topological considerations, can be used to shed some light into the types of resulting patterns possible.

### Hierarchical gene networks

By definition, hierarchical networks do not have extracellular loops and, due to requirement *I2*, hierarchical gene networks leading to non-trivial pattern transformations include at least one positive intracellular loop. As we saw in the previous section, this means that these networks are RD-unstable of the second kind.

#### Pattern transformations from spike initial patterns in H networks with a single extracellular signal

Let us begin by considering H gene networks consisting of a single H subnetwork with a single extracellular signal. We call these H^0^ networks. From a spike initial pattern, it is only the cells in the spike that, by definition, express the extracellular signal. This implies that it is only these cells that are secreting the extracellular signal. Thus, the concentration of the extracellular signal in all other cells (i.e., cells outside the spike) is fully governed by signal diffusion from this source and its degradation. Indeed, if we consider the set of cells either on the right-hand side of the spike (computations are equivalent for the left-hand side), we can solve equation (1) assuming linear degradation in the signal to show that its concentration forms an exponential gradient around the spike,

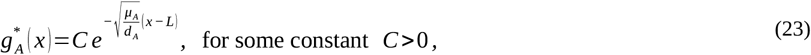

where *A* is the extracellular signal in the H^0^ network, 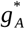 is its concentration in the resulting pattern, *x* is the distance to the border of the spike and *L* is half the width of the spike. In 2D, the signal concentration gradient decays as 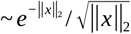, where ||*x*||_2_ denotes euclidean norm; and in 3D, as 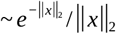 (Sommerfeld, 1949; Stakgold & Holst, 2011). In all cases signal concentration has a radially decaying spatial distribution centered on the spike. In section S3 of the appendix we calculate *C*.

In the spatial distribution arising from (23), each cell at each side of the spike is at a unique distance from the spike and, thus, experiences a unique concentration of *A*. In that sense, there is a unique correspondence between the concentration of *A* and the distance to the spike. This correspondence has been used in the literature (Wolpert, 1968) to propose that, with a single extracellular signal, any pattern transformation is possible (see Fig.7). The only thing that would be required is that cells interpret the signal concentration in a different way for each different resulting pattern attained (Capek and Müller, 2019; Sharpe, 2019). However, what would that interpretation be, or which would be its underlying gene network, was not specified by this author (Horder, 2001). Allegedly, any interpretation should rely on specific gene networks. A common idea for such interpretation is that cells express specific genes if they receive *A* at concentrations beyond some threshold value that is different for different genes. Then, different genes may become expressed at different distances from the spike and, thus, a non-trivial pattern transformation would occur (Capek and Müller, 2019; Sharpe, 2019).

**Figure 7.**
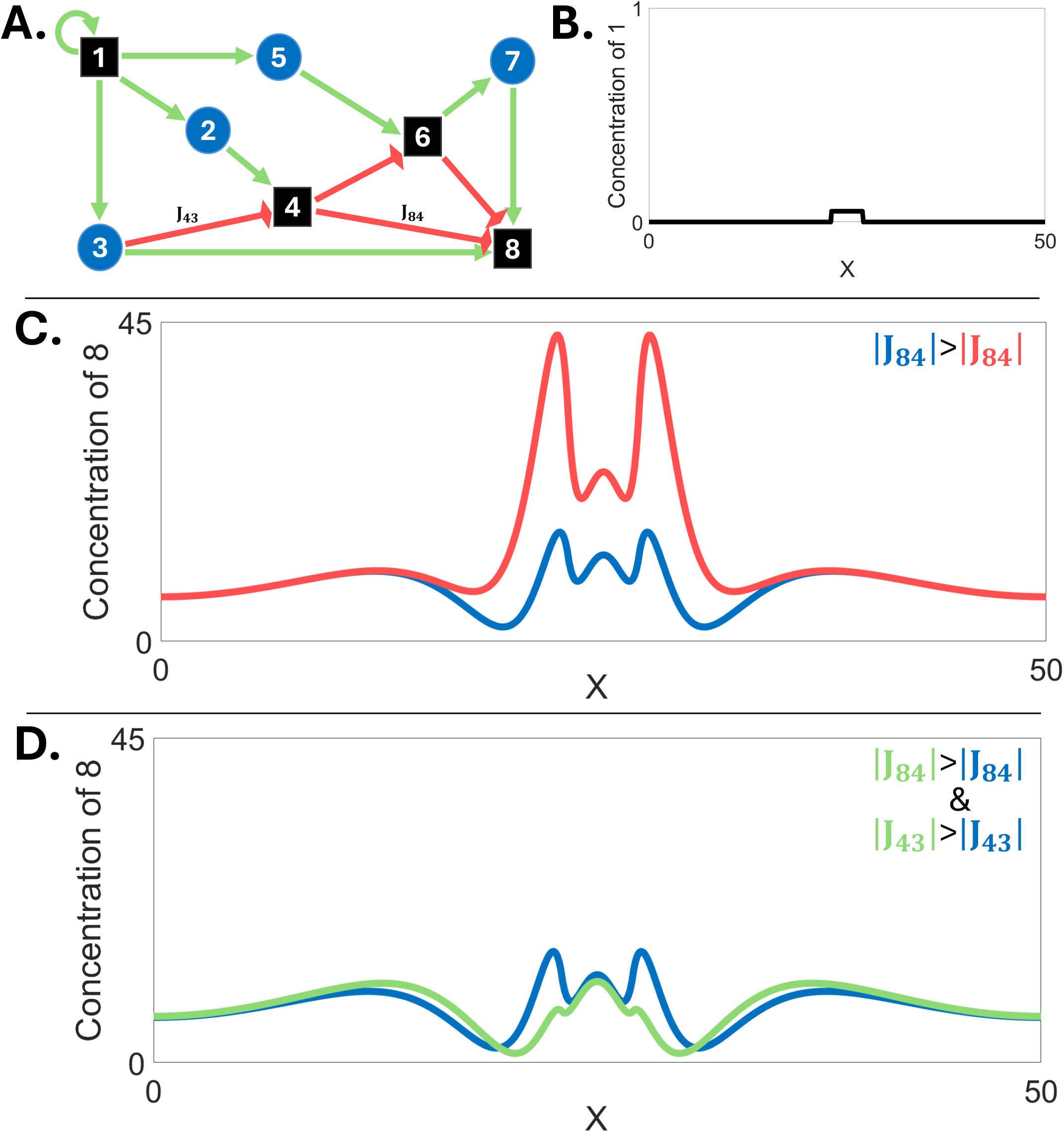
If function *f* did not satisfy R5, then a two gene network could lead to any resulting pattern in which concentrations are finite. **(A)** Gene network topology. An extracellular signal activating an intracellular gene product. 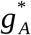 stands for the concentration of the extracelllular signal in the steady state and 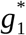 for that of the intracellular gene product. **(B)** The extracellular signal is secreted from the boundary of an array of cells (x-axis), leading to a gradient in the concentration of the concentration of the signal. Labels c1 to c6 are shown to facilitate the interpretation of the figures but we assume *f* can be understood as continuous. **(C)** Different example functions *f* are plotted. In the x-axis we represent the different concentrations of the signal and in the y-axis the concentration of the intracellular gene product arising from each concentration of the signal. Notice that none of the example functions *f* satisfy R5, i.e. the function is not monotonously growing with the concentration of the signal **(D)** The resulting patterns arising from each *f* . By choosing an adequate *f*, any resulting pattern can be achieved.

There are, however, many other ways in which H^0^ networks can lead to non-trivial pattern transformations. In the following, we explain some fundamental requirements that H^0^ networks need to fulfill in order to lead to non-trivial pattern transformations and which are their possible non-trivial pattern transformations. This new requirements should be added on top of requirements R1-R5 to enable non-trivial pattern transformations in H^0^ networks.

##### Requirement RH1

For non-trivial pattern transformations in H^0^ networks, at least one gene product that is positively downstream of the extracellular signal *A* (let us say gene product *k*) must inhibit at least one gene product that is also positively downstream of *A* (let us say gene product *j*).

Requirement (RH1) has been previously discussed by other authors (Monteanu *et al*., 2014), especially for networks with three gene products, but we include it and expanded it here for completeness (see Fig.8). This requirement can be split into two parts: 1) there needs to be at least two gene products that are positively downstream of *A*; and, 2) one of them has to inhibit the other. Part 1 is simply requirement I1 applied to H^0^ networks: any gene product undergoing a non-trivial pattern transformation must be positively downstream of an extracellular signal and in H^0^ networks, this is *A* because it is the only signal there is.

**Figure 8.**
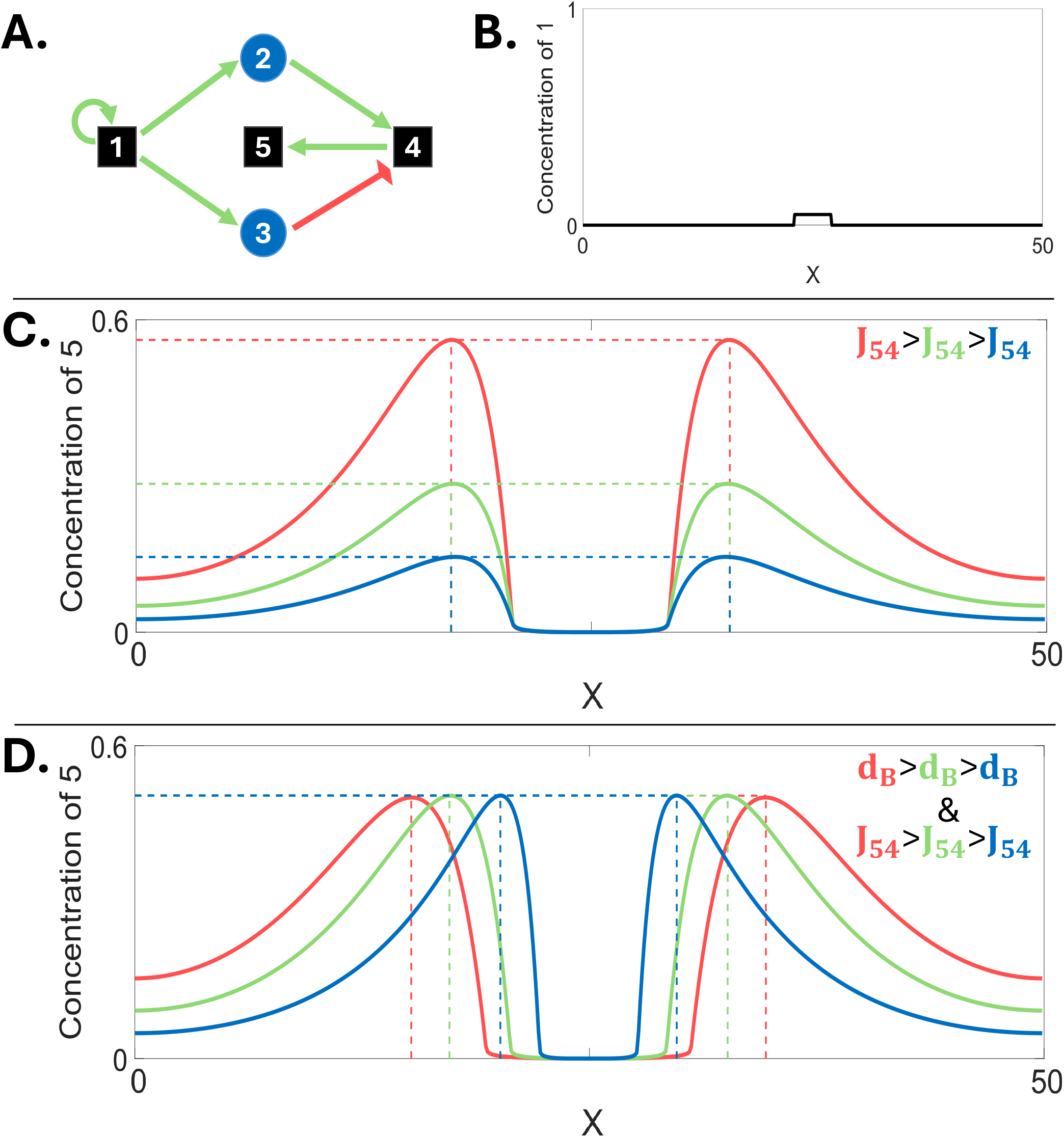
**Diagram showing the topological requirements for pattern transformation in hierarchical gene networks** using example gene networks. (A) Requirement (RH2a): the positive regulation of 3 by *A* (discontinuous green arrow) has a different non-linearity than that of 4 by *A*. (B) Requirement (RH2b): the negative regulation of 5 by 3 (discontinuous red arrow) has a different non-linearity than that of 5 by 4. (C) Requirement (RH2c): two extracellular signals downstream of 1 and upstream of 4. The signals have different concentration profiles over space because either they have different diffusion coefficient or different intrinsic degradation rates. Each of these requirements lead to the formation of two new concentration peaks in the most downstream gene product. Network colors and shapes as in Figure 2. Simulations were run using a Forward-Euler algorithm on the Maini-Miura model (see S6 above for parameter values). Notice that in **(A)** gene product 3 is an example of whatcall gene product *j* in the main text, while in **(B)** gene product *j* corresponds to gene product 5.

To understand part 2, let us first consider H^0^ networks in which part 1 holds but part 2 does not i.e., *j* and *k* are both positively downstream of *A*, but *j* is not inhibited by *k*. In that case, according to requirement R5 on *f*, both *k* and *j* will increase their concentration when that of *A* increases, and decrease their concentration when that of *A* decreases. Since the concentration of *A* decreases away from the spike (23), it follows that the concentrations of *k* and *j* also decrease with the distance to the spike and thus, *k* and *j* have their concentration peak in the same place as *A* and, thus, no non-trivial pattern transformation occurs. In fact, the same will occur for any gene products that, in a H^0^ network, are positively, and only positively, downstream of *A*.

Let us now consider the case in which part 1 and part 2 of RH1 hold. In that case, there is at least one gene product, *k*, that is downstream from *A* and inhibits another gene product, *j*, that is also downstream from *A*. In this case, the concentration of *j* can become very small where that of *k* is large (i.e., in the spike) and thus, form a concentration valley centered in the spike. This valley will in turn be flanked by two concentration peaks of *j* at each side (see Fig. 9). In the 2D and 3D cases there would be a central basin and a ridge surrounding it (see Fig.9-10). However, the emergence of these concentration peaks and valleys is only possible if function *f* fulfills at least one of the two following additional requirements (see Fig.8):

**Figure 9.**
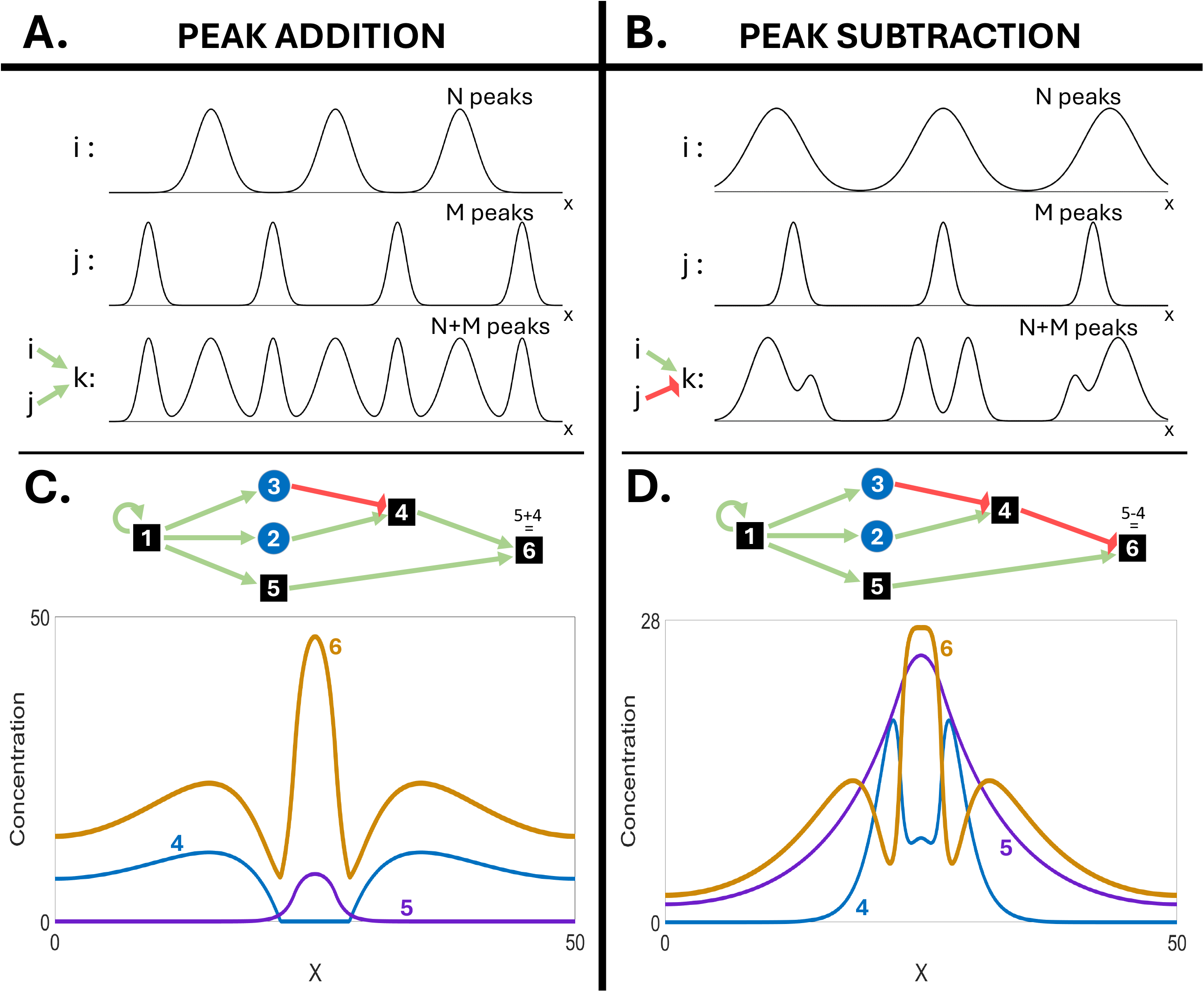
Classes of gene networks capable of pattern formation and their resulting patterns in 1D. The first column depicts simple examples of each class of gene network topology capable of non-trivial pattern transformation. The upper row shows the three initial patterns. Intermediate panels show each type of possible resulting pattern arising from each combination of initial pattern and gene network topologies. The network topologies in the left column are only simple ones, for a more detailed description of the possible ones check the main text. For the over-Turing topology we chose to represent a intracellular loop with two gene products (but we could have chosen one). Note that some of the pattern transformations are trivial (over-Turing and Turing on spike initial pattern). H network can lead to non-trivial pattern transformations from homogeneous-with-noise initial patterns. Network colors and shapes as in Figure 2, except that *P* stands for the gene product plotted as resulting pattern while *I* stands for a gene product present in the initial pattern. Simulations were run using a Forward-Euler algorithm on the Maini-Miura model for *f* (see S6 in the appendix for parameter values).

##### Requirement RH2a

function *f* is such that the proportion between the activations of *k* and *j* by *A* changes with the concentration of *A* (i.e., *f* _*k*_ (*g* _*A*_, …)/ *f* _*j*_ (*g* _*A*_, …) is not a constant function of *g* _*A*_). In other words, the functional form of the activation of *k* by *A* (i.e., *f* _*k*_ (*g* _*A*_, …)) has a different non-linear part than the functional form of the activation of *j* by *A* (i.e., *f* _*j*_ (*g* _*A*_, …)).

##### Requirement RH2b

The inhibition of *j* by *k* is non-linear in the sense that it is disproportionately larger for high concentrations of *k* than for low concentrations of *k*.

Both requirements imply an inhibition of *j* that is disproportionately different for different values of *g* _*A*_ and, thus, for different distances to the spike. In RH2a, this disproportionality arises from the different non-linear parts of the activations of *j* and *k* by *A*, while in RH2b, it arises from an intrinsic non-linearity in the inhibition of *j* by *k* (see Fig.8).

To better understand these requirements lets consider what happens if neither RH2a nor RH2b hold and all relevant interactions are linear. In this case, gene products *k* and *j* have concentrations that are, in each cell, proportional to those of *A* (and, thus, proportional to each other). In this case, if *k* inhibits *j* linearly, then *g* _*j*_ decreases proportionally to *g*_*k*_ in each cell. Consequently, *g* _*j*_ remains proportional to *A* over space but it is simply expressed at a lower overall level (i.e., no non-trivial pattern transformation can occur).

If the inhibition of *j* by *k* is disproportionately higher when *g*_*k*_ is large i.e. non-linear inhibition, then *g* _*j*_ would be disproportionately smaller where *g*_*k*_ is large (i.e. around the spike) than where *g*_*k*_ is small (i.e. away from the spike). This can lead to the formation of a valley of *g* _*j*_ in and around the spike and, consequently, to the formation of two peaks of concentration around it (in two or higher dimensions that would be a basin and a ridge surrounding it, see Fig.10).

**Figure 10.**
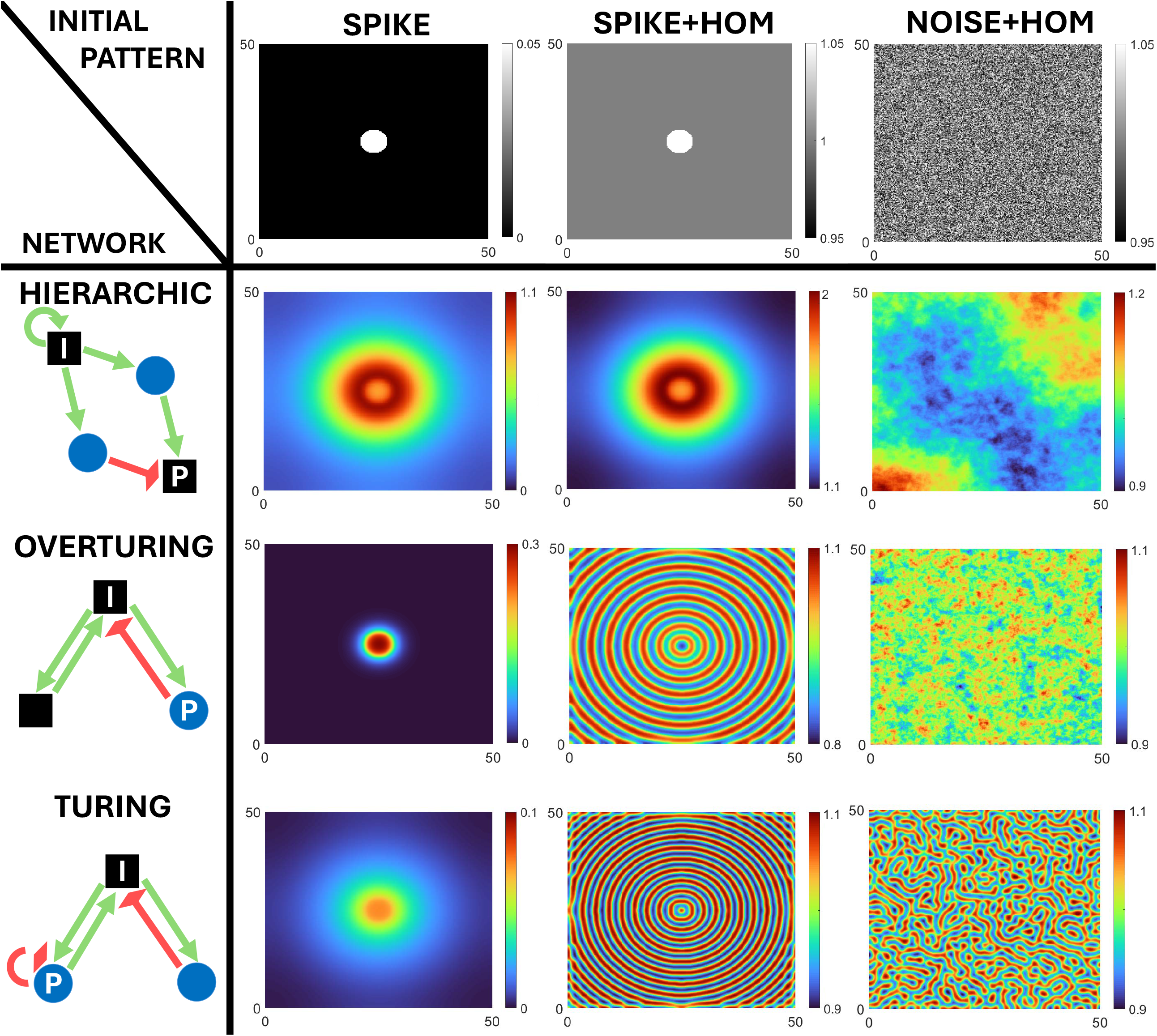
Classes of gene networks capable of pattern formation and their resulting patterns in 2D. The first column depicts simple examples of each class of gene network topology capable of non-trivial pattern transformation. The upper row shows the three initial patterns. Intermediate panels show each type of possible resulting pattern arising from each combination of initial pattern and gene network topologies. In the case of spike and spike-homogeneous initial patterns, the resulting patterns correspond to the revolution of the patterns in Figure 8 around the center of the spike given the radial symmetry of equation (1) and the spike. In homogeneous-with-noise initial patterns, the white noise breaks such symmetry. In the case of Turing networks acting on homogeneous-with-noise initial pattern other types of periodical resulting patterns (e.g., dots or stripes) have been reported in the literature (Murray, 2002). Network colors and shapes as in Figure 8, Simulations were run using a Forward-Euler algorithm on the Maini-Miura model for ***f*** (see section S6 in the appendix for parameter values).

To illustrate these arguments lets us consider an example gene network and *f* in which signal *A* directly activates gene products *j* and *k*, and *k* directly inhibits *j* in such a way that *j* and *k* are regulated by no other gene product in the network. In *H*^*0*^ networks the concentration of *A* is given by equation (23) and the dynamics of *j* and *k* are given by (24).

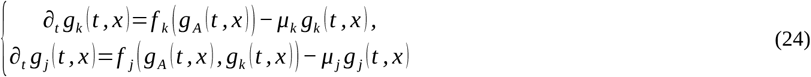

Where we assume, for simplicity, that both *j* and *k* suffer linear degradation (i.e., *m*_*j*_=*m*_*k*_ =1). Now, assume that regulations *f* _*k*_ (*g* _*A*_) and *f* _*j*_ (*g* _*A*_, *g*_*k*_) are both linear. Then, system (24) reads,

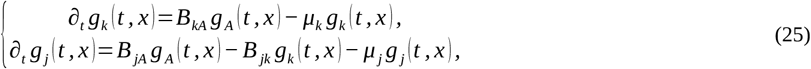

where *B*_*kA*_, *B* _*jA*_, *B* _*jk*_ >0 are real positive constants. Given that we are only interested in stationary resulting patterns, we can take time derivatives in (25) to zero and then we find that the concentrations of *k* and *j* in the resulting pattern are given by,

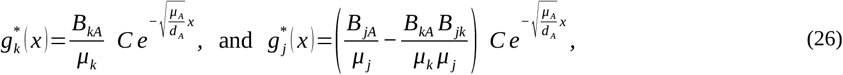

where we have used the gradient shape of 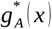 from (23). Then, as we can see in (26), both concentrations have negative exponential distributions in space with the same exponential decay but different heights (i.e., different leading coefficients). Moreover, equation (26) also shows that there is a single concentration peak for *A, j* and *k*, and that these peaks are all centered at the same position (i.e. the center of the spike) and, thus, there is no non-trivial pattern transformation. This proves that no non-trivial pattern transformations are possible if all regulations are linear (requirement R2).

Now, let us consider the case in which all regulations remain linear except for the inhibition of *j* by *k* (as requested by RH2b). Then, system (24) becomes,

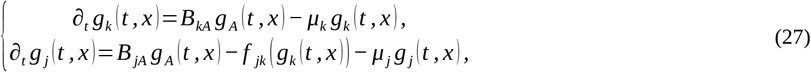

where *f* _*jk*_ is a non-linear inhibition of *j* by *k*. Hence, taking time derivatives in (27) equal to zero, the resulting pattern concentrations of *j* by *k* are,

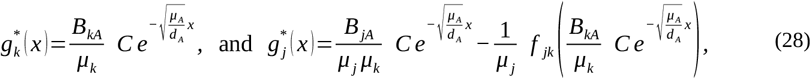

where we have used the gradient shape of 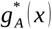 from (23). For gene product *k*, we have the same result as before: a single concentration peak centered in the spike. Things are, however, different for the resulting pattern of gene product *j*. From requirement R5, we know that *f* _*jk*_ is monotonically increasing with *g* _*A*_ and hence, it decreases with *x*. The two terms in 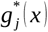 have a maximum at *x*=0 and then, since the second term is negative, the overall value of 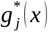 in (28) can be lower at *x*=0 than elsewhere. If this happens, a concentration valley of *j* can form at *x*=0, flanked by two concentration peaks of *j* at each side of it. This implies the formation of new concentration peaks and valleys, and thus, that non-trivial pattern transformations are possible (see Fig.8) when requirement (RH2b) holds.

Next, let us consider the case in which RH2a holds and RH2b does not. In this case, the activation of *k* by *A* at high concentrations of *A* is disproportionately larger than that of *j by A*. As a result, *g*_*k*_ would be disproportionately larger than *g*_*j*_ in, and close to, the spike and not away from it (i.e. *g*_*k*_ and *g*_*j*_ would decay with a different slope from the spike). Then the inhibition of *j* by *k*, even when linear, can lead to the formation of a valley of *g*_*j*_ in and around the spike and, consequently, to the formation of two concentration peaks away from the spike (in two and higher dimensions that would be a concentration ridge and basin, see Fig.10).

Let us illustrate these arguments with a specific example in which regulations *f* _*k*_ (*g* _*A*_) and *f* _*j*_ (*g* _*A*_, *g*_*k*_) are non-linear, but the activations of *k* and *j* by *A* have a different non-linear part in their functional forms (as required by RH2a) In this case, system (24) reads,

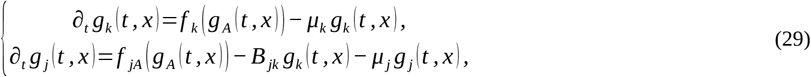

where *f* _*jA*_ (*g* _*A*_) is a non-linear activation of *j* by *A* that is different from that of *f* _*k*_ (*g* _*A*_) (e.g., one is a quadratic monomial and the other, a cubic monomial). We maintain linear inhibition of *j* by *k* because we are considering the case in which RH2a holds but RH2b does not. As in previous cases, if we equal time derivatives in (29) to zero, we get that the stationary resulting patterns of *j* and *k* are given by.

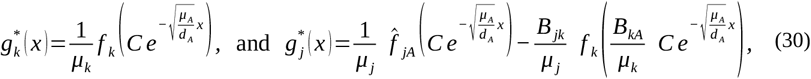

where we have used (23). Requirement R5 ensures that *f* _*k*_ (*g* _*A*_) and *f* _*jA*_ (*g* _*A*_) are both monotonically increasing functions of *g* _*A*_ and consequently, monotonically decreasing functions of *x*. For gene product *k*, this means that its resulting pattern will have the same monotony as 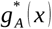 (including its unique maximum at *x*=0), but with a different slope. Similarly, the two terms in 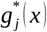 share that same monotony with a maximum at *x*=0 but different slopes (e.g., if 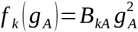 and 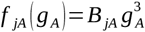, then 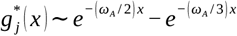, with 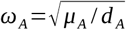. Consequently, since the second term is negative, the overall value of 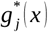 in (30) can be lower at *x*=0 than it is in nearby regions and, if this happens, a concentration valley of *j* can form at *x*=0, flanked by two concentration peaks of *j* at each side of it. Like before, when and where these concentration peaks and valleys form depends on the specific choice of the parameters (again, the parameters choice must be such that 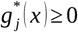 for all *x*). However, if they form, then gene product *j* would undergo a non-trivial pattern transformation as two new concentration peaks emerge at positions where there was no previous concentration peak (see Fig.8). This proves that requirement (RH2b), together with the proper choice of the parameters, enables non-trivial pattern transformations in H^0^ networks.

It is worth mentioning that, although we used a very simple H^0^ network for the illustration of requirements RH2a and RH2b, both requirements apply even if the regulations involved are not direct (e.g., if *k* activates some intermediate gene product *i* and *j* activates *j*). In this case, the different non-linear part in the functional form of regulations can be in any intermediate regulation (e.g., in the activation that *k* exerts on *i*, or the one that *i* exerts on *j*).

#### Pattern transformations from spike initial patterns in general H networks

In general, H networks may have different extracellular signals and these may affect patterning in downstream gene products. These networks can also lead to non-trivial pattern transformations and they do it with a set of less restrictive requirements than H^0^ networks (see Fig.8C). In this case, inhibition does not need to occur between gene products downstream of the same extracellular signal *A*, but can occur between gene products downstream from different extracellular signals secreted from the spike, e.g. signals *A*_1_ and *A*_2_. Then, requirements RH1, RH2a and RH2b are no longer necessary as long as these different extracellular signals have different diffusion coefficients (i.e., 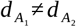), or different degradation coefficients (i.e., 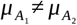). Hence, former additional requirements shall be substituted by the following one:

##### Requirement RH2c

For non-trivial pattern transformations H networks with several extracellular signals, there must exists some gene product *j* that is positively downstream of one extracellular signal *A*_1_ and negatively downstream of another extracellular signal *A*_2_, both secreted from the spike and with different diffusion coefficients (or different degradation rates).

If this occurs, then according to (23), these two signals have concentration gradients with different slopes along x. In other words, some of the extracellular signals have a higher proportion of their concentration close to the spike than others. As in the previous case then, concentration valleys can form around the spike for those gene products that are inhibited by one signal and activated by the other, directly or indirectly (see Fig.8). To illustrate this possibility, let us consider a simple H network where extracellular signal *A*_1_ directly activates an intracellular gene product *j*, while signal *A*_2_ directly inhibits it. Thus, if we assume that regulation 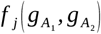 is linear, the concentration dynamics of *j* is given by,

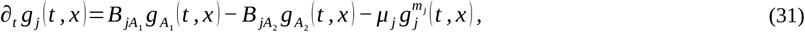

where 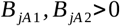 are positive real constants. Then, taking the time derivative in (31) equal to zero, we have that the emerging patter of *j* is given by,

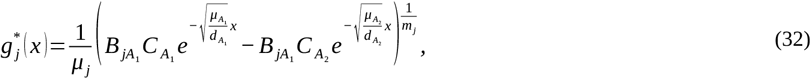

Equation (32) is essentially the difference of two negative exponential functions. Then, if the slope of the first exponential is smaller than that of the second exponential, this subtraction can lead to the formation of concentration valley in 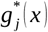 at *x*=0 flanked by two concentration peaks at each side of it (thus leading to non-trivial pattern transformation).

#### Pattern transformations arising from the combination of multiple H subnetworks

Gene products that undergo a non-trivial pattern transformation through an H network can lead to further non-trivial pattern transformations in downstream gene products without the need for activating additional extracellular signals (Scalise & Schulman, 2014). This can occur in gene networks in which gene products with a heterogeneous resulting pattern activate a gene product that is, in turn, inhibited by gene products with another different pattern (see Fig. 11). The resulting pattern in the downstream gene product can then have concentration valleys where its inhibitory gene products have concentration peaks, and concentration peaks where its activating gene products have concentration valleys (we call this peak subtraction).

**Figure 11.**
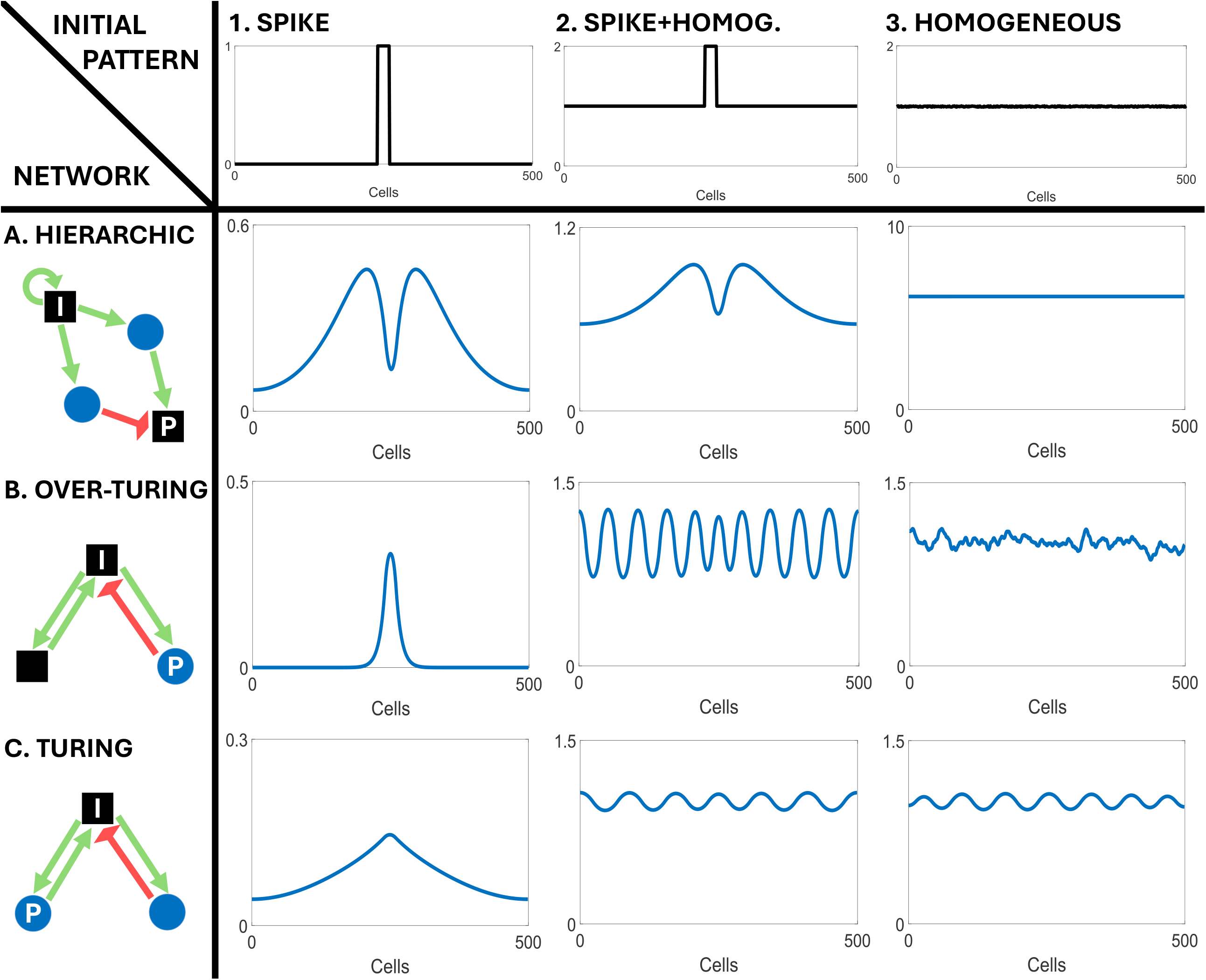
Peak addition and subtraction in the resulting pattern of downstream gene products from hierarchic gene networks. The most upper row shows an idealized pattern of a gene product *i* with several concentration peaks. The middle row that of another gene product, *j*, with several concentration peaks. In **(A)** both gene products positively regulate the same gene product, *k*. As a result gene product *k* has concentration peaks either *i* or *j* have them. In **(B)**, instead, *j* inhibits *k* while *i* activates it. **(C)** shows an example network in which the pattern of two gene products, 4 and 5, are effectively added into that of gene product 6. **(D)** shows and example gene network in which the patterns of gene products 4 is effectively subtracted from that of gene product 5 to give rise to the pattern of gene product 6. Simulations were run using a Forward-Euler algorithm on the Maini-Miura model for *f* (see section S6 in the appendix for parameter values).

Similarly, a gene product that is positively downstream of gene products with different patterns can also develop a new pattern in which concentrations peaks would form wherever any of the upstream gene products have concentration peaks, as long as the concentration peaks of upstream activators are sufficiently distant from one another, (see Fig.11). We call this peak addition.

In both cases, the downstream gene product may acquire new concentration peaks and valleys and, thus, peak addition and subtraction can lead to non-trivial pattern transformations. The total number of peaks in the resulting pattern is less than or equal to the sum of the number of peaks in the resulting pattern of upstream gene products (as explained in section S4 and S5 of the appendix).

#### The ensemble of possible pattern transformations from spike initial patterns in H networks

As we have seen in the previous subsection, H subnetworks can be combined in simple ways to lead to resulting patterns with many concentration peaks and valleys. In this subsection we argue that, with some broad restrictions, for any radially symmetric resulting pattern there is a H network producing it from a spike initial pattern.

Let us first acknowledge that any resulting pattern can be seen as a sequence of concentration peaks of different heights and steepness at different positions (or basins and ridges in higher dimensions). Now, let us consider a simple gene network that we call the diamond network (see Fig.12A). This gene network has an intracellular gene product in the spike (say gene product 1) that self-activates and activates two extracellular signals (say signals 2 and 3) that, in turn, regulate a downstream intracellular gene product (say gene product 4). Gene product 4 is activated by signal 2 and inhibited by signal 3. Finally, the gene network includes an intracellular gene product 5 that is activated by gene product 4.

**Figure 12.**
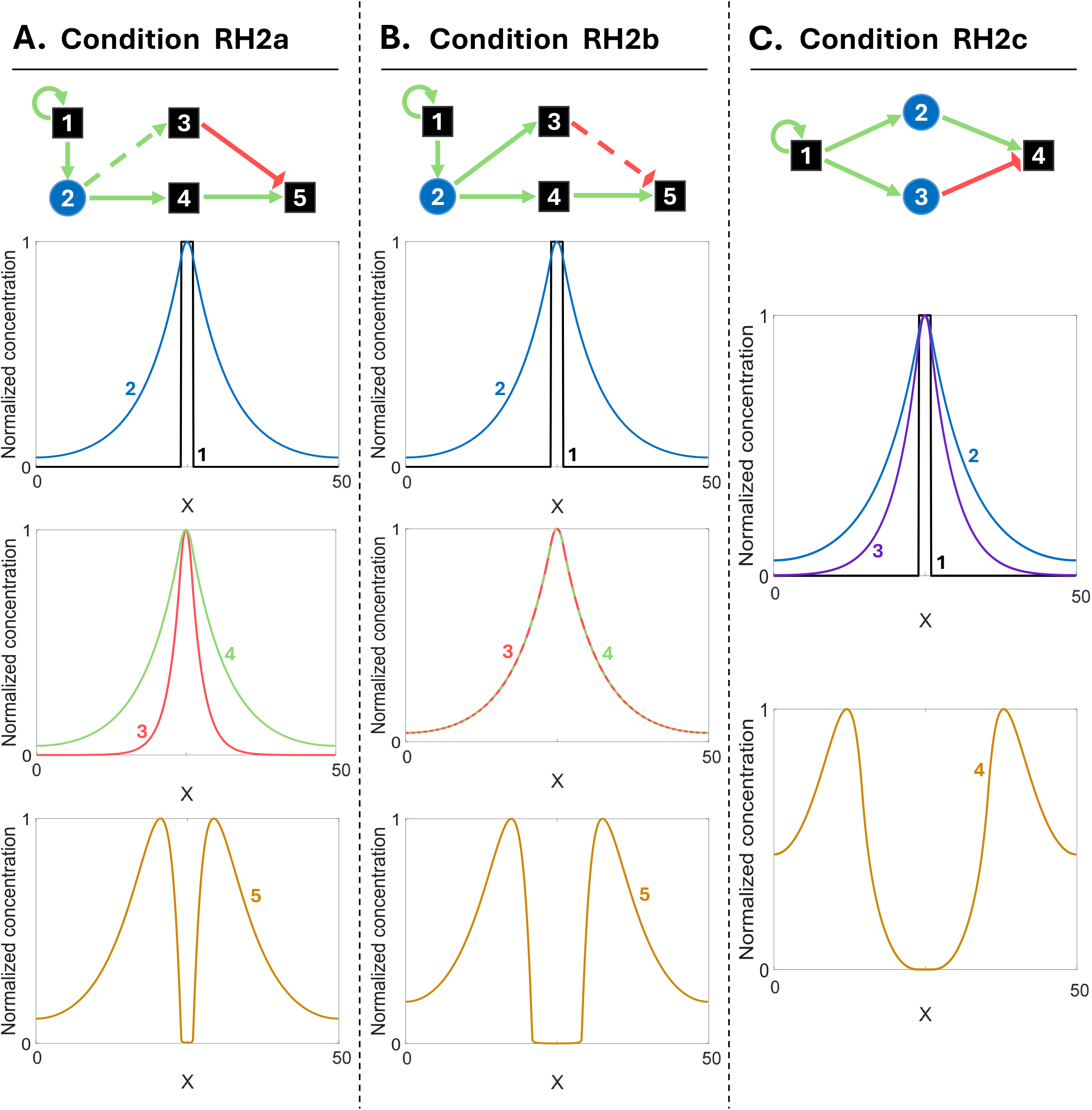
Variational properties of the diamond H network. **(A)** Diamond H network (see section S4 in the appendix). Network colors and shapes as in Figures 2 and 4. **(B)** Initial pattern in gene product 1. **(C-D)** The resulting patterns consist in two symmetric peaks around the initial spike. The height **(C)** and position **(D)** of such peaks can be independently modified by tuning the model parameters. Peak height can be varied by changing a single parameter, that is the activation of gene product 5 by gene product 4 (i.e. *J* _54_). To change the position of the peak while peaking its height constant **(D)** two parameters need to be changed at the same time, the diffusion coefficient of extracellular signal and the activation of gene product 5 by gene product 4. Just changing the diffusion coefficient the position of the peaks will vary but so will their height. Simulations were run using a Forward-Euler algorithm on the Maini-Miura model for *f* (see section S6 in the appendix for parameter values).

As we know from previous subsections, if extracellular signals 2 and 3 have different diffusion coefficients, or degradation rates, the resulting pattern of 4 can exhibit a pair of concentration peaks at a distance from the spike. With some very mild restrictions (specified in S4 of the appendix), this distance can be made larger or smaller by changes in these parameters and the rest of parameters (see S4 in the appendix for a more formal discussion) (see Fig.12). The resulting pattern of gene product 5 will exhibit concentration peaks in the same place as gene product 4. However, by choosing an adequate *f* _5_ (*g*_4_) (while keeping R1 to R5) the concentration peaks of gene product 5 can have different heights and steepness (i.e., different decrease rates from its peak). In the most extreme case (e.g., using a sigmoidal function for *f* _5_ (*g*_4_)), the concentration of gene product 5 can be made to be only appreciable in the cell where it attains its peak and its most immediate neighboring cells (i.e. very narrow concentration peak).

This simple diamond gene network can be combined any number of times to produce any number of peaks in any combination of positions and heights: simply, gene product 1 would activate different diamond gene networks and each such networks would have different parameters that lead to concentration peaks in different locations and with different heights and steepness. Then, the peak addition described above (see Fig.11) can put together all such peaks in the resulting pattern of a gene product downstream of all such diamond networks. Hence, the resulting pattern can produce any array of concentration peaks and valleys (i.e. target resulting pattern) as long as it is symmetric around the spike, it is continuous over space, and peaks and valleys are sufficiently distant from each other (peaks that are very close together may merge; see S4 in the appendix). Moreover, it is worth noting that if the resulting pattern consists of very narrow concentration peaks, function *f* and its parameters may need to be tuned very precisely and thus, patterns made of extremely narrow peaks may not be biologically feasible in practice.

The hierarchical gene networks studied in the previous sections are just simple examples. Since these simple H gene networks can lead to nearly all resulting patterns, one can expect that there are other, more complex, H networks that can also do it.

Finally, it is worth noting that hierarchical gene networks acting on a spike initial pattern tend to have similar variational properties (as studied in Salazar-Ciudad et al., 2000, 2001). Thus, in many H networks, the properties of the different peaks and valleys in a pattern (i.e. their height, position and steepness) are able to vary in respect to those of many other peaks in a pattern (while keeping radial symmetry). In the combination of diamond networks just described, for example, each pair of peaks arises from a specific diamond subnetwork with specific extracellular signals and its own parameters. This implies that it is possible, as we have seen, to regulate the height, steepness, and position of each pair of peaks in the resulting patterns (or, of each ridge in higher dimensions) independently from each other through changes in the parameters of each subnetwork (see Fig. 12-13 for other simple examples). However, not in all H gene networks all peaks can vary independently from each other. Peak addition and subtraction, for example, can lead to some peaks varying together with parameter changes (Salazar-Ciudad *et al*., 2000).

#### Pattern transformations from combined spike-homogeneous initial patterns in H networks

The pattern transformations possible from the combined spike-homogeneous initial patterns are similar to those possible from the spike initial conditions and have the same properties (see Fig.9-10). It suffices to add steady-state concentration 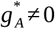 to the signal gradient in (23) and all subsequent arguments in the previous subsections follow similarly.

#### Pattern formations from homogeneous-with-noise initial patterns in H networks

As we have seen in previous section, for pattern transformations to be possible, the homogeneous steady state needs to be unstable to perturbations. This means that the gene network, *f* and the parameters must be such that equations (1) admit at least three equilibrium points (the initial steady state and some others equilibrium points).

Gene networks that admit only two such equilibrium points cannot lead to non-trivial pattern transformations. The reason is that the initial equilibrium point needs to be unstable for pattern transformation to occur and that noise is applied to all cells. If the initial equilibrium is indeed unstable, then noise will lead all cells to change from the initial unstable equilibrium point to the second equilibrium point. In other words, the homogeneous-with-noise initial pattern will transforms into another homogeneous pattern with just different concentrations of the gene products.

Gene networks that admit three or more equilibrium points can lead to pattern transformations. In this case, cells may move to one equilibrium point or another depending on the magnitude of the random perturbation they initially receive. For example, in the case of gene networks that admit three different equilibrium points 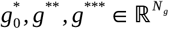 such that 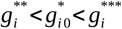 for all gene product *i*, the gene product concentrations in each cell where the initial concentration is below 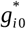 may change to 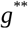. Similarly, any cell where the initial concentration is above 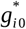 may change to 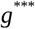. This way cells randomly settle in different equilibrium points (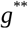 and 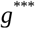 in the example).

If the above gene networks do not include any extracellular signal, the resulting pattern will simply be a random distribution of these equilibrium points among cells. This will be, in other words, a random distribution of concentration peaks and valleys. Each of these peaks and valleys will form where the initial pattern also had, due to noise, tiny peaks and valleys. In that sense, no new concentration peaks and valleys appear and, thus, without signaling, there are no non-trivial pattern transformation (not for H networks nor in any other gene networks).

In the case of H gene networks and homogeneous-with-noise initial patterns there are, by definition, extracellular signals and, thus, concentration peaks and valleys can form where the initial pattern had tiny concentration peaks and valleys (as explained in the previous paragraph) but also at a distance from them, as explained for H gene networks under other initial patterns. In this case, thus, new peaks and valleys can arise and non-trivial patterns transformations are possible. The resulting patterns, however, still consist of random distributions of peaks and valleys (basins and ridges in higher dimensions)(see Fig.9-10). This is, in fact, what the dispersion relation indicates. Hierarchical networks are RD-unstable of the second kind and, thus, have an infinite number of unstable wavenumbers. Except for sampling limitations, the initial pattern with white noise contains all wavenumbers. This means that many, if not most, wavenumbers present in the initial noise can be amplified and, thus, the resulting patterns most likely consist of random distributions of concentration peaks and valleys (see Fig.9-10).

### Emergent gene networks

#### L^+^ gene networks

These are gene networks with one or several L^+^ subnetworks (and no other types of subnetworks). From a spike initial pattern, the signal, or signals, in the L^+^ subnetworks diffuse from the spike across the extracellular space and activate their own production in the cells that receive it. Since these signals promote their own production, the cells receiving them also end up producing and secreting them. As a result, each signal progressively spreads further until it ends up being expressed by all cells (see Fig.9-10). Due to the averaging effect of diffusion and the self-activating nature of L^+^, all cells end up having the same concentration of the signal (even noise is suppressed) and, thus, the only possible resulting patterns are homogeneous (i.e. no non-trivial pattern transformation occurs). The same happens for homogeneous-with-noise or combined spike-homogeneous initial patterns.

#### L^-^ gene networks

By definition, these networks only contain subnetworks with extracellular signals that inhibit their own production, directly or indirectly, in the cells that receive them. According to requirement I2, gene networks leading to non-trivial pattern transformations must include one or several positive loops (i.e. l^+^) positively upstream of the gene products present in the resulting pattern. For L^-^ gene networks, these loops have to be intracellular, otherwise the gene network would be L^+^L^-^ not L^-^. Accordingly, we must only consider the L^-^ networks that include at least one L^-^ subnetwork downstream of at least one l^+^.

#### Pattern transformations in L^-^ subnetworks from spike initial patterns

In the spike initial pattern, only the cells in the spike initially express the genes of the L^-^ subnetwork. Since the secreted signal inhibits itself, the expression of this signal cannot spread beyond the spike (i.e. it will only be secreted from the spike) and, thus, its peak concentration will be in the spike itself (i.e. no non-trivial pattern transformation).

#### Pattern transformations in L^-^ subnetworks from homogeneous-with-noise initial patterns

Lets us first consider the gene networks with a single L^-^ subnetwork and a single l^+^. Among these, let us first consider the case of L^-^ gene networks with a single L^-^ subnetwork positively downstream, but not upstream, of a single l^+^. Since l^+^ is not affected by any extracellular signal, its gene products can only have a homogeneous resulting pattern, or a noisy resulting pattern like the one described for gene networks without extracellular signals (in the “Pattern formations from homogeneous-with-noise initial patterns in H networks” section). Neither case constitutes a non-trivial pattern transformation. By definition, the most upstream extracellular signal, *A*, inhibits itself. Then it can only be expressed in the same place than its positive regulators, l^+^, and thus, these gene networks cannot lead to non-trivial pattern transformations either. The case of a L^-^ subnetwork upstream of an l^+^ but not downstream of it does not fulfill requirement I2 and, thus, cannot lead to pattern transformations either.

The only case left is that of a L^-^ subnetwork that is both downstream and upstream of the l^+^. In the simplest case, l^+^ activates *A* and *A* inhibits some gene product in l^+^, and thus indirectly all of them and itself. We call these networks over-Turing gene networks since, as we will see, they are capable of producing periodic patterns that resemble those of classical Turing gene networks (see Fig.9-10). Since the only positive loop is intracellular, these gene networks are, depending on the parameters, either RD-unstable of the second kind or RD-stable (see Fig.6C-D) (Klika *et al*., 2012).

In each cell, l^+^ activates *A* but *A* diffuses extracellularly and inhibits l^+^ in the cells that receive it and, thus indirectly itself. This means that, indirectly, the l^+^ in the different cells are competing with each other. In other words, by inhibiting the l^+^ of the other cells, the l^+^ in a cell is indirectly activating itself. If the system is made of just two cells, the concentration of the gene products in l^+^ (as well as *A*) can only increase in one cell (the one experiencing a weaker effective inhibition of *A* by l^+^ at the initial time) and decrease in the other (the one subject to a stronger initial inhibition of *A* by l^+^). This implies that the initial difference in the concentration of *A* between cells can only but grow until some steady state may be reached (in accordance with requirement R4). In other words, the initial concentration difference between cells (i.e. the initial noise in the pattern) becomes amplified.

The two-cell example can be used to understand larger systems with homogeneous-with-noise initial patterns. These systems can be understood as being composed of multiple overlapping pairs of cells that, due to noise, have small initial differences in the concentration of the gene products in l^+^ (or any gene product upstream of it). From the arguments explained in the previous paragraph, one can expect that the initial concentration differences between each pair of cells can only but grow. In the case of systems made of many cells, however, each cell is contiguous to more than one cell (e.g., two in 1D) and, thus, it can occur that two contiguous cells both increase their concentration of *A* because they are both contiguous to other cells that, by chance, have smaller initial concentrations of *A*. Which cells end up in concentration peaks or valleys depends on which cells had higher signal concentration in the initial pattern and the inhibition arising from the extracellular signals, as well as on the parameters of the network.

The resulting pattern consists of randomly distributed concentration peaks and valleys of *A* and the gene products in l^+^ (see Fig.9-10). These patterns are very similar to the ones that can arise from H networks acting on homogeneous-with-noise initial patterns (even if the underlying dynamics are quite different). This is, in fact, what the dispersion relation indicates in the case of RD-unstable networks of the second kind: high wavenumbers are unstable and thus, they all can be amplified to form a noisy resulting pattern. Indeed, the resulting pattern can be understood, to some extent, as an amplification of the initial noise in the homogeneous-with-noise pattern (Marcon *et al*., 2016; Diego *et al*., 2018). Because of this, each resulting pattern will be different, with different peaks and valleys in different positions and different distribution of heights between them (see Fig.14A).

**Figure 13.**
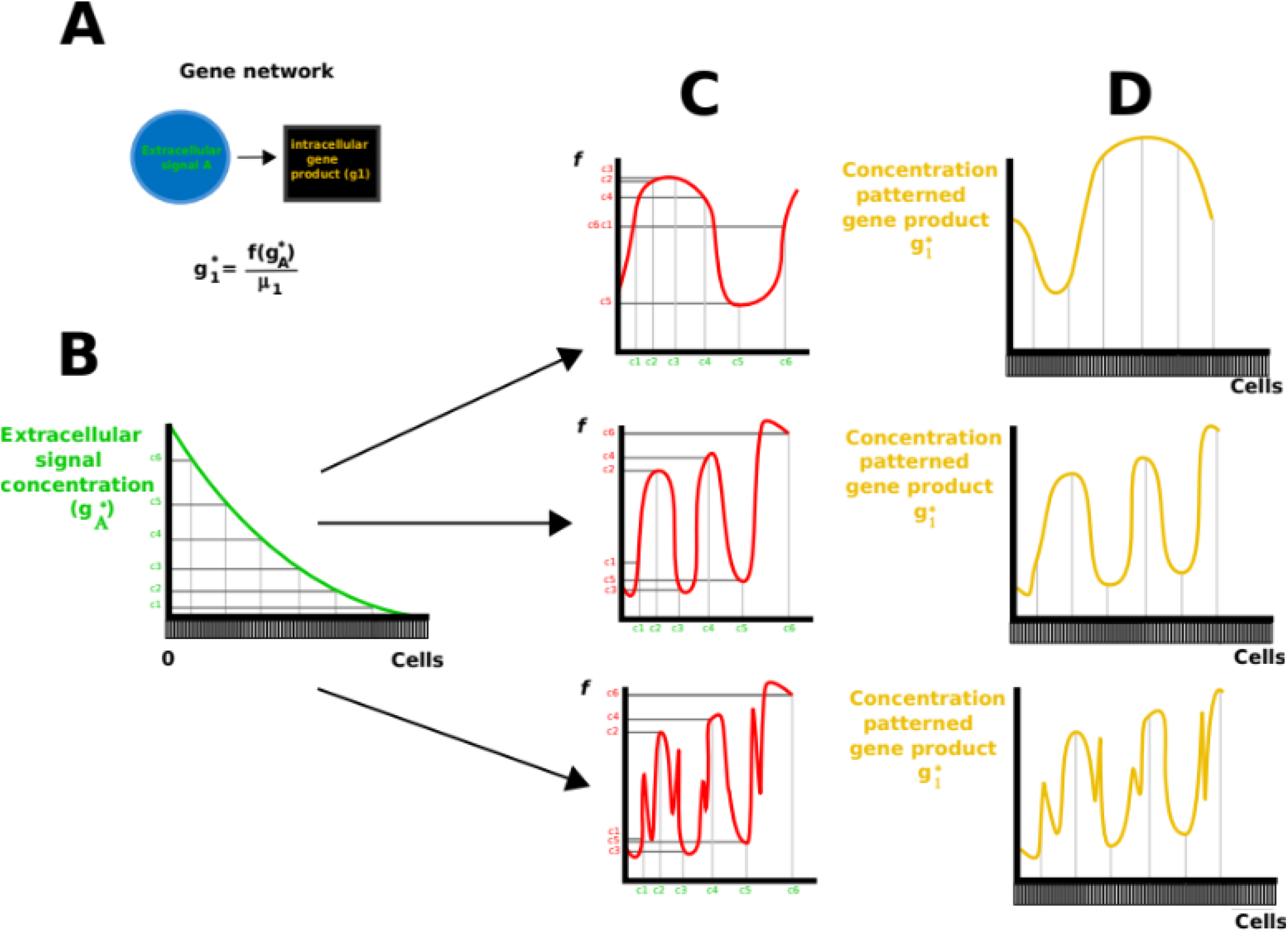
Example H gene network where by changing the parameters it is possible to vary the height of the different concentration peaks. **(A)** Gene network topology. **(B)** Spike initial pattern. **(C-D)** The blue, red and green plots show the concentration of gene product 8 for different combinations of parameter values. As we can see, the height of the concentration peaks of 8 around the initial spike can be independently tuned. Simulations were run using a Forward-Euler algorithm on the Maini-Miura model (see section S6 for parameter values).

**Figure 14.**
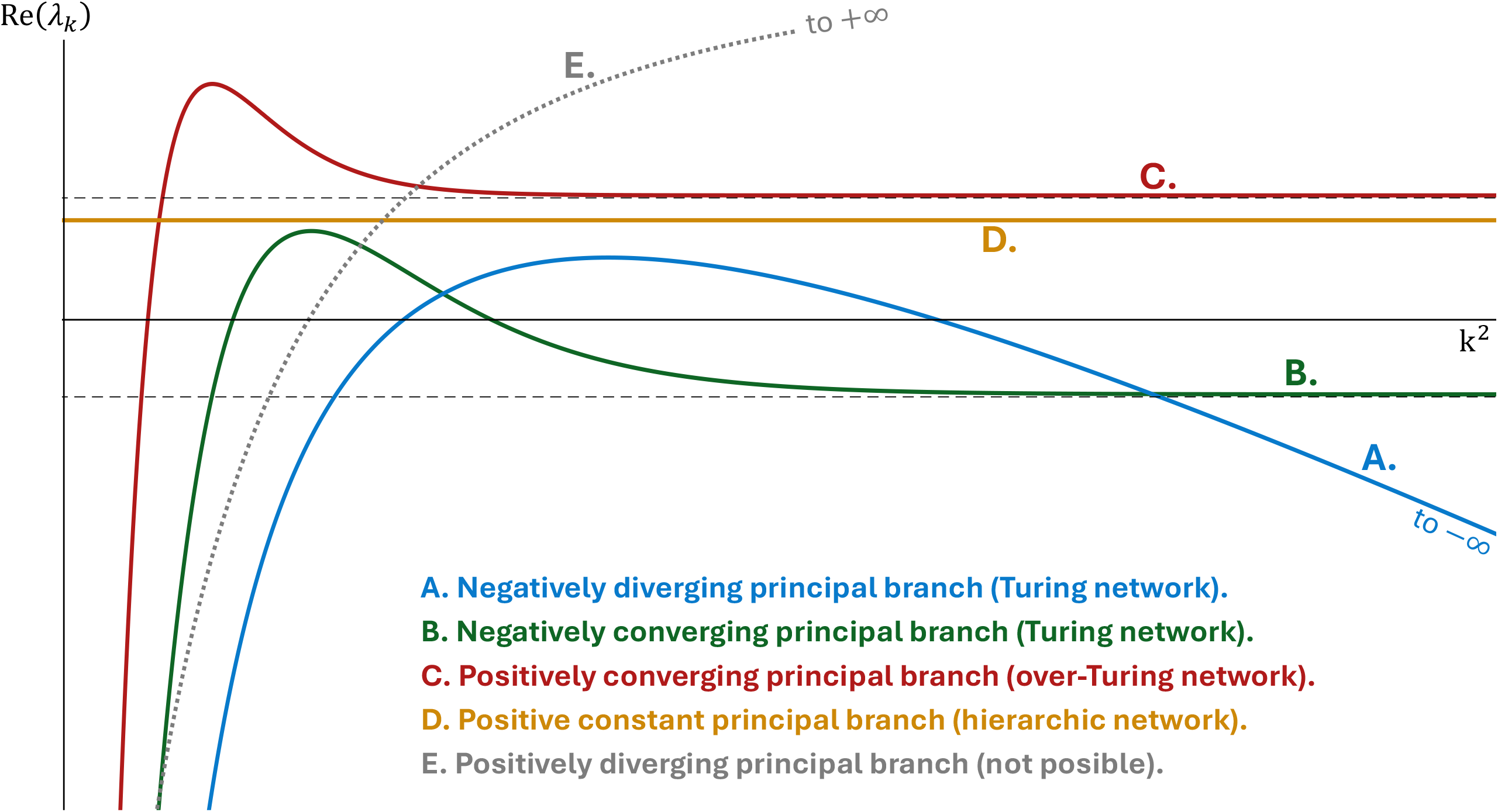
Pattern formation in over-Turing gene networks. **(A)** Noise-homogeneous initial patterns are transformed into amplified noise patterns. Even with the same choices of model parameters, the amplified noise resulting patterns are different every time. **(B)** Spike-homogeneous initial patterns lead to periodical patterns. If the choice of model parameters is the same, the final pattern is the same for each transformation. **(C)** The peaks of concentration formed from a spike-homogeneous initial pattern are produced one by one. The whole process resembles a traveling wave that moves away from the spike and, at the same time, freezes whenever a new peak is formed. Network colors and shapes as in Figure 2. *P*stands for the gene product plotted as resulting pattern while *I* stands for the gene product in the initial pattern. Simulations as in Figure S2 (see section S6 for parameter values).

Let us now consider the ensemble of over-Turing gene networks. This is any gene network combining any number of l^+^ loops and L^-^ subnetworks downstream and upstream of each other (but no L^+^ subnetwork). In all of them it is still the case, by the definition of L^-^ subnetworks, that extracellular signals inhibit their own production in the cells that receive them and, that thus, cells are inhibiting each others’ production of each extracellular signal, directly or indirectly. This means that, as in the simple examples above, any initial difference in the initial concentration of extracellular signals between cells can only increase. In fact, over-Turing networks contain positive intracellular loops and, thus, irrespectively of their complexity, they can only be, depending on the parameters, RD-unstable of the second kind or RD-stable. In the former case, most wavenumbers would be unstable and, since there is an infinite number of them, the resulting patterns should include many of them, i.e. the resulting pattern will likely be a random distribution of peaks and valleys as in the case of the simplest over-Turing networks.

#### Pattern transformations in L^-^ subnetworks from spike-homogeneous initial patterns

As in the previous initial patterns, and for the same reasons, only the over-Turing gene networks can lead to non-trivial pattern transformations. Over-Turing gene networks are known to be able to lead to relatively complex and periodic non-trivial pattern transformations from the spike-homogeneous initial pattern (Marcon *et al*., 2016; Wang *et al*., 2022). By definition, the concentration of *A* starts being higher in the spike than elsewhere. As *A* diffuses, it inhibits itself in the cells around the spike and this leads to the formation of concentration valleys of *A*. However, *A* activates gene products in l^+^ and thus, similar concentration valleys of gene products in l^+^ will form around the spike (see Fig.14B-C). Since l^+^ activates signal *A*, the cells in these valleys will produce less *A* and, in turn, cells beyond the valleys will receive a smaller amount of it. As a consequence, there is less inhibition of the gene products in l^+^ in such cells, and the concentration of *A* and the gene products in l^+^ will increase in those cells. This increase in the concentration of *A* leads to the formation of a concentration valley further away, and these valleys to further peaks, and so on. This process continues until the whole domain is occupied by a sequence of concentration peaks and valleys of *A* and the gene products in l^+^. These have very similar heights, shapes and distances between them (see Fig.14), with some variation in those closer to the spike or the boundary of the domain. In 2D there are no concentration peaks but concentric rings of high and low gene product concentration centered around the spike. In 3D there are concentric spherical shells of low and high concentration. We refer to these patterns as frozen-wave patterns as they result from the action of a patterning wavefront that moves away of the spike leaving a stationary wave-like pattern behind. As in the case of H networks, these frozen-wave patterns are symmetric around the initial spike but, in contrast to H networks, they extend over the whole system.

The formation of these frozen-wave resulting patterns can also be understood as a consequence of the fact that, in over-Turing networks, any initial difference in the concentration of *A* between contiguous cells can only increase (as we explained in the previous subsection). In this case, the initial concentration differences are between the cells at the margin of the spike and the cells just outside of it in every radial direction. These differences can only but grow over time and thus, the concentration of *A* and gene products in l^+^ will decrease in cells just outside of the spike. This, in turn, leads to a concentration difference between the cells just outside the spike and cells further away, in which the concentration of *A* and gene products in l^+^ will increase again. This process continues leading to an alternation over space of concentration peaks and valleys (or concentric concentration rings and shells in higher dimensions) or, in other words, to the frozen-wave pattern.

The frozen-wave patterns cannot arise from spike initial patterns. This is because concentration valleys cannot form where gene product concentrations are zero (i.e. around the spike), since gene product concentration cannot be negative.

Changes in the parameters of over-Turing gene networks (e.g., diffusion coefficients) can lead to changes in the number, height or width of concentration peaks and valleys in the resulting pattern. However, contrary to what happens for H gene networks, concentration peaks and valleys in over-Turing networks cannot change independently from each other (see Fig.15). This lack of variational independence in the properties of peaks and valleys is easy to understand from the fact that over-Turing gene networks produce many different concentration peaks from a small number of interactions in the gene network and thus, no specific part of the network is responsible for any specific concentration peak (as it can happen in H gene networks). This implies that the ensemble of pattern transformations attainable by these networks is less diverse than the ensemble of pattern transformations attainable by H gene networks. In other words, the over-Turing gene networks can produce patterns with equally sized and shaped peaks, while the H networks can also produce patterns with concentration peaks and valleys with different characteristics (different heights and shapes, Salazar-Ciudad *et al*., 2000, 2001).

**Figure 15.**
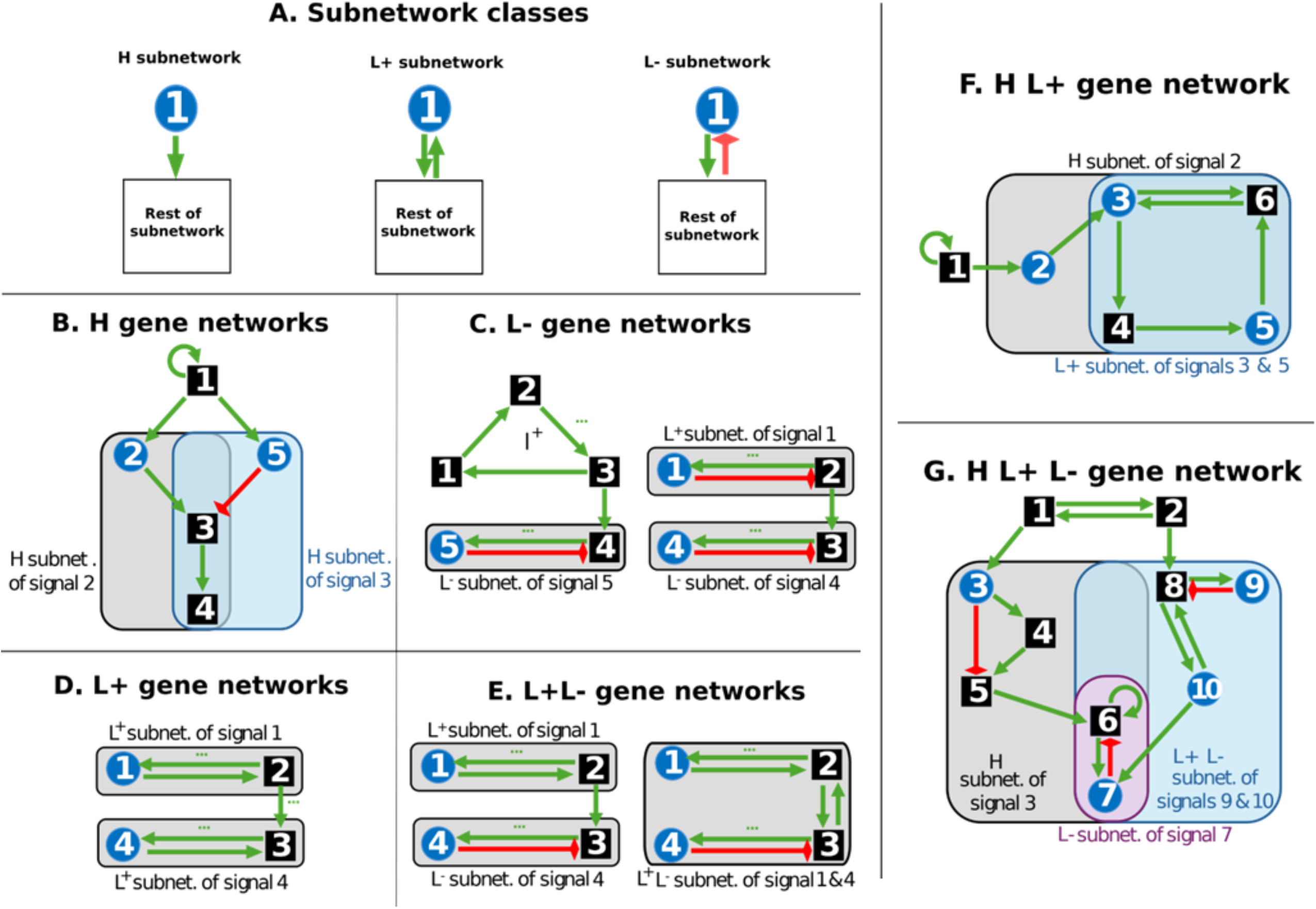
Variational properties of emergent networks. **(A-B)** An over-Turing (left) and Turing (right) gene networks produce a very similar periodical pattern from a spike-homogeneous initial pattern. A change in any model parameter (such as the diffusivity of an extracellular signal) alters the height and position of all peaks at the same time. **(C)** Spike initial pattern. **(D-E).** Network colors and shapes as in Figure S1. *P* stands for the gene product plotted as resulting pattern while *I* stands for the gene product in the initial pattern. Simulations as in Figure S3 (see section S6 for parameter values).

### Gene networks combining different classes of subnetworks

Subnetworks can be combined in parallel, in series, or both. By parallel combinations we mean that different subnetworks are upstream of the same set of gene products (while being downstream of some gene product in the initial pattern, requirement I1). Since in this case the different subnetworks do not regulate each other, the resulting patterns in downstream gene products can have concentration peaks and valleys wherever their upstream gene products have them (just as described for H gene networks above, see Fig.11).

By in series combinations we mean that some subnetworks are upstream of others (i.e., the signal of the first subnetwork is upstream of the signal of the other). In this case we are interested in the resulting patterns of the downstream subnetwork (since the pattern transformations in the upstream subnetworks are unaffected by the downstream gene network and have already been described in previous sections).

Subnetworks can be combined in series positively or negatively (i.e., the signal of the upstream subnetwork is negatively upstream of the signal of the other subnetwork). In the case of negative regulation, however, some gene products in the initial pattern must positively regulate the downstream subnetworks independently of the upstream subnetwork, otherwise the genes in the downstream subnetwork only receive negative regulation and, thus, cannot be expressed (i.e. requirement I1). In this case, the resulting pattern of the downstream network would be the one it would produce on its own, but with that of the other subnetwork subtracted from it, as we described for H subnetworks (see Fig. 11). The case of positive in series combinations of subnetworks is described in the next section. Naturally, several subnetworks can be combined in series to form a chain in which each subnetwork is upstream of the next one.

Subnetworks can also be combined in series but forming a loop (i.e., the signal of each subnetwork is upstream and downstream of the signals of other subnetworks). However, if an H subnetwork is combined forming a loop with any other class of subnetwork, then the signal of the H subnetwork is indirectly regulating itself and is, thus, an emergent subnetwork. In this sense, combining subnetworks in a loop leads to emergent subnetworks.

#### Pattern transformations in the combination of H and L^+^ and H and L^-^ subnetworks

As we have seen, all gene networks capable of pattern transformation can be RD-unstable of the first kind or RD-unstable of the second kind. Thus, regardless of how sub-networks are combined, there are only two main types of resulting patterns. H and L^+^ subnetworks can only be RD-stable or RD-unstable of the second kind (since the only positive loop is extracellular). Beyond that we can acquire a more detailed intuition of the possible pattern transformations through some simple considerations on the pattern transformations we have described for the H and L^+^ subnetworks individually.

Gene networks composed of L^+^ subnetworks upstream of H subnetworks are equivalent to H subnetworks acting on a homogeneous initial pattern without noise. This is because, as explained in the L^+^ section, these subnetworks lead to homogeneous resulting patterns whichever the initial pattern and, in the process, eliminate noise. Thus, the downstream H subnetworks effectively receive and input pattern that is homogeneous (i.e. without noise) and, thus, cannot lead to pattern transformations.

In gene networks in which H subnetworks are upstream of L^+^ subnetworks no pattern transformations are possible downstream of the H subnetworks because the L^+^ subnetworks homogenize any input patterns that the H subnetworks may convey.

Gene networks composed of L^-^ subnetworks upstream of H subnetworks can only lead to non-trivial pattern transformations in the case the L^-^ subnetworks are over-Turing, for the reasons explained in the L^-^ subnetworks section. The same applies to gene networks composed of H subnetworks upstream of L^-^ subnetworks. In this case the H subnetwork can be seen as providing the L^-^ subnetworks with a pattern but as we have seen L^-^ subnetworks cannot lead to non-trivial pattern transformations unless they are over-Turing, regardless of the initial pattern they act on.

In the case of H subnetworks upstream of over-Turing subnetworks and combined spike-homogeneous initial patterns, the frozen-wave pattern that would normally spread from the spike perturbation can spread, instead, from the peaks and valleys produced by the H subnetwork. Thus, the resulting is still a frozen-wave pattern but locally modified by the H subnetwork so that the peaks close to the initial spike can vary in size (this effect keeping radial symmetry around the spike) (see Fig.16). If the over-Turing subnetworks are upstream of H subnetworks, then the radially symmetric patterns that are attainable by the latter can be repeated around each concentration peak of the amplified noise pattern. When acting on a homogeneous-with-noise initial pattern, the only possible non-trivial pattern transformations are the noisy resulting patterns reported for both over-Turing and H networks (since both kinds of networks produce similar patterns on their own anyway).

**Figure 16.**
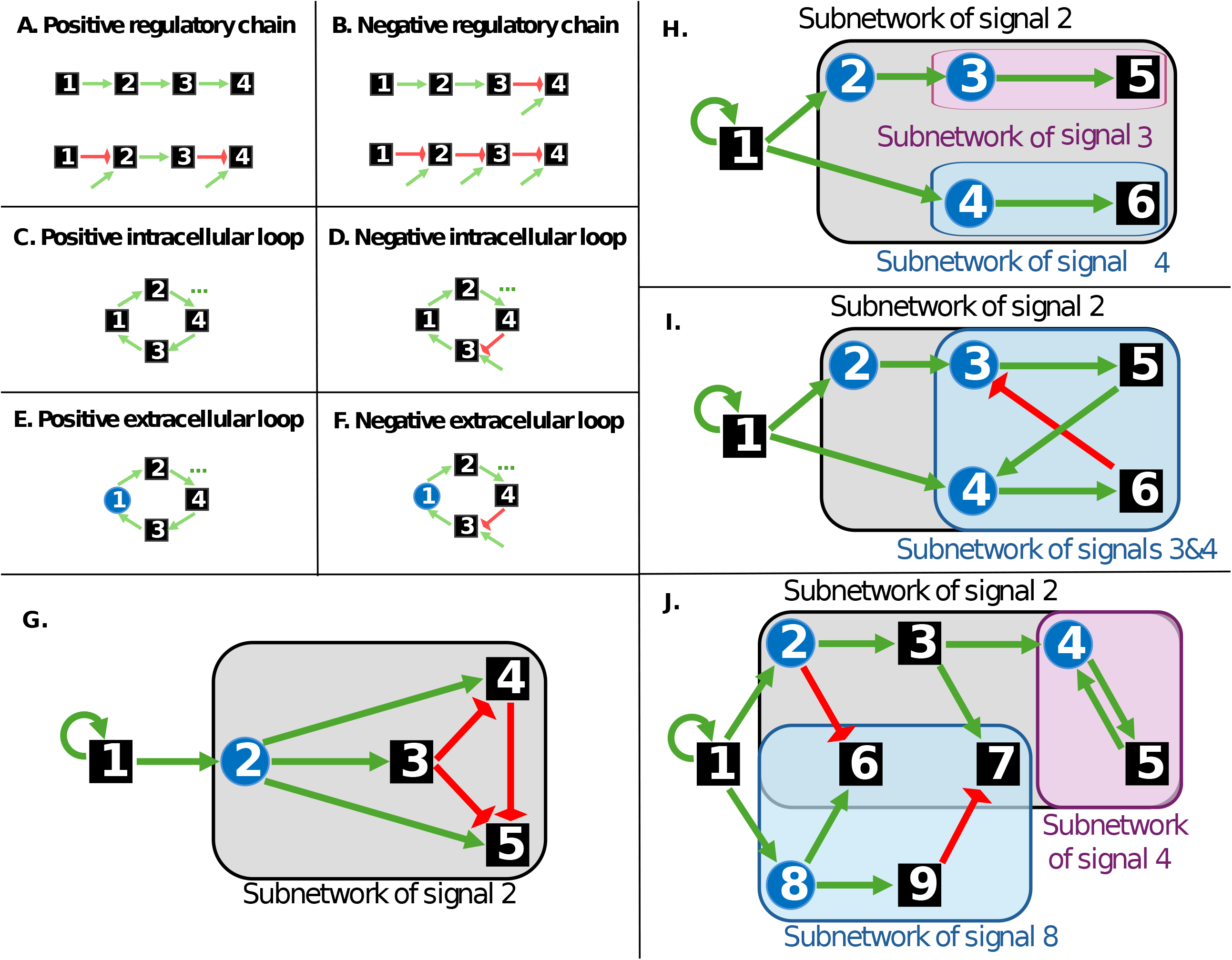
The combinations of hierarchical, over-Turing and Turing networks are and their resulting patterns. The first row depicts simple examples of each class of gene network topology acting as the upstream subnetwork of the combination in series. The first column shows simple examples of each the same gene network topologies acting as the downstream subnetwork of an in series combination. Intermediate panels show each type of possible resulting pattern arising from a spike-homogeneous initial pattern from each combination. Note that all pattern transformations are non-trivial. Network colors and shapes as in Figure 2. Dashed arrows represent the different, and strictly non-linear, regulations in the H^0^ networks. *P* stands for the gene product plotted as resulting pattern while *I* stands for the gene product in the initial pattern. Simulations were run using a Forward-Euler algorithm on the Maini-Miura model for *f* (see S6 in the appendix for parameter values).

#### Pattern transformations in the combination of L^+^ and L^-^ subnetworks

In this section we first consider combinations between single L^+^ and single L^-^ subnetworks. If an L^+^ subnetwork is upstream of an L^-^ subnetwork, and not downstream of it (i.e., they form no loop), the L^+^ subnetwork is unaffected by the L^-^ subnetwork and consequently, its genes will be homogeneously expressed regardless of the initial pattern (see Fig. 17-18). This reduces to what we have seen is possible for L^-^ gene networks acting on homogeneous-with-noise initial patterns, except that the L^+^ subnetwork can eliminate the noise and preclude any pattern transformation. If the L^-^ subnetwork is upstream of the L^+^ subnetwork, then the resulting pattern of the latter subnetwork can only be homogeneous because, as we explained in previous subsections, L^+^ subnetworks produce these patterns whenever any of its signals is expressed in at least one cell (see Fig 17-18).

**Figure 17.**
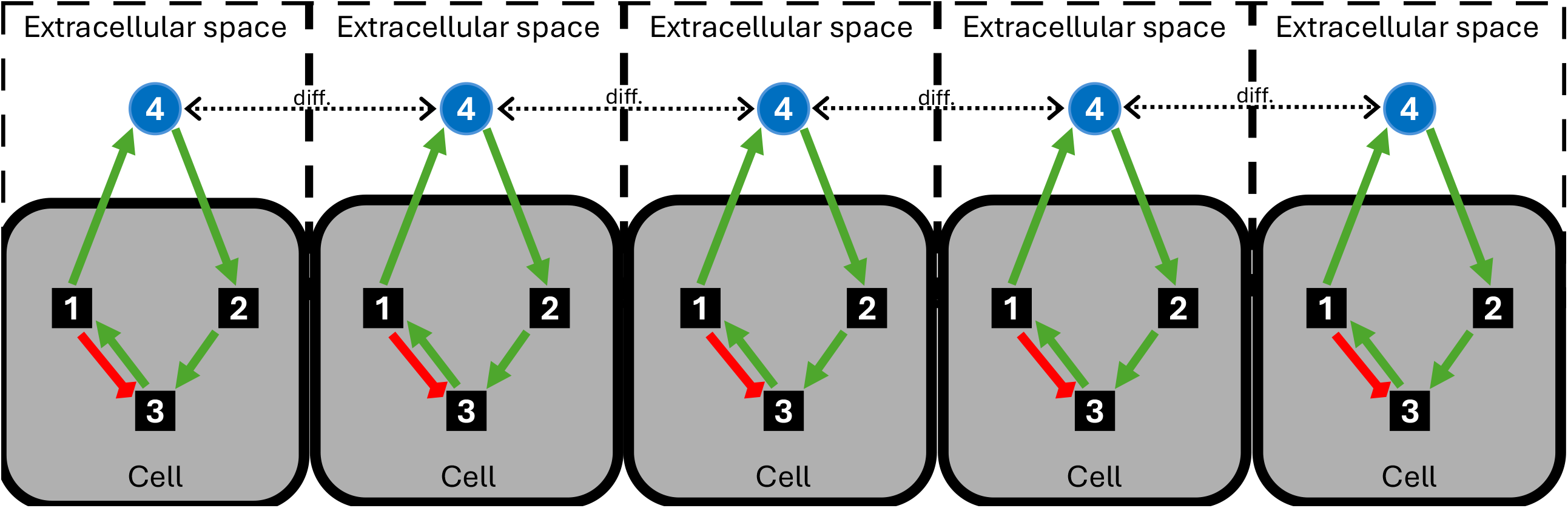
L^+^, l^+^L^-^ and L^+^L^-^ networks do not produce non-trivial pattern transformations. The first column depicts simple examples of each type of these gene network topologies. The upper row shows the three initial patterns. Intermediate panels show each type of possible resulting pattern arising from each combination of initial pattern and gene network topology. Note that all pattern transformations are trivial. Network colors and shapes as in Figure S1. The y-axis in all plots represents gene product concentrations. *P* stands for the gene product plotted as resulting pattern while *I* stands for the gene product in the initial pattern. Simulations as in Figure S3 (see section S6 for parameter values).

**Figure 18.**
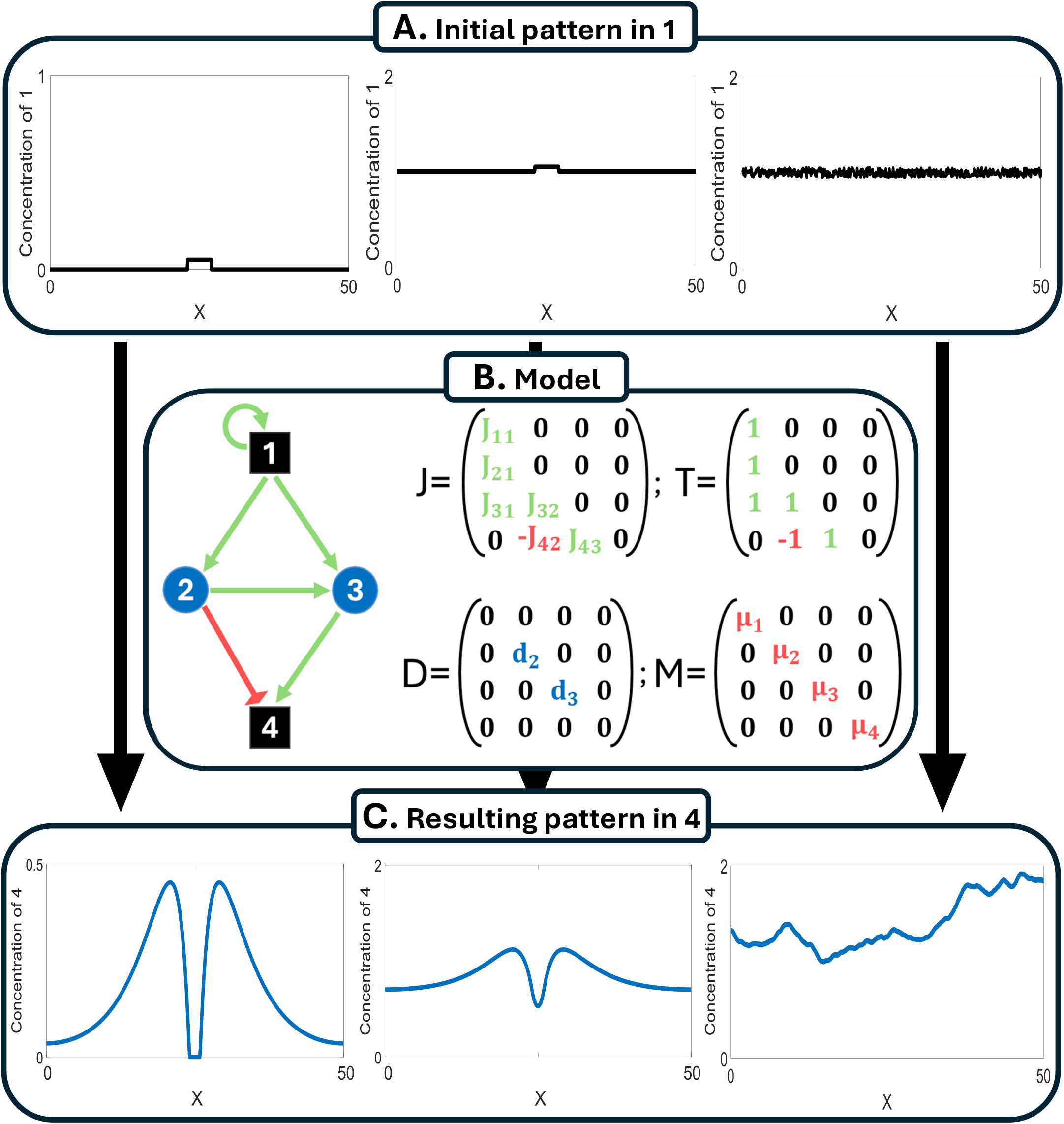
L^-^, L^-^l^+^ and L^-^L^+^ networks do not produce non-trivial pattern transformations. The first column depicts simple examples of each type of these gene network topologies. The upper row shows the three initial patterns. Intermediate panels show each type of possible resulting pattern arising from each combination of initial pattern and gene network topology. Note that all pattern transformations are trivial. Network colors and shapes as in Figure S1. The y-axis in all plots represents gene product concentrations. *P* stands for the gene product plotted as resulting pattern while*I* stands for the gene product in the initial pattern. Simulations as in Figure S3 (see section S6 for parameter values).

L^+^ and L^-^ subnetworks can be combined into loops. Previous research has extensively studied pattern transformations occurring from such networks (Cotterell & Sharpe, 2010), for the case of gene networks with up to three gene products. From these, the only resulting patterns and developmental dynamics that we have not discussed so far are those of Turing gene networks, also called Turing mechanisms (Turing, 1952; Murray, 2002; Maini *et al*., 2006; Meinhardt, 2008). These occur, for example, when a L^+^ subnetwork is positively upstream of an L^-^ subnetwork, and the latter is negatively upstream of the former (e.g., the signal of the latter inhibits, directly or indirectly, the signal of the former).

In Turing’s seminal work (Turing, 1952), each subnetwork is in fact a single extracellular signal, but it has been widely reported that these mechanisms also apply to gene networks where extracellular signals regulate each other indirectly, through an intracellular part of the gene network (Salazar-Ciudad, *et al*. 2000; Satnoianu, *et al*., 2000; Maini *et al*., 2006; Meinhardt, 2008; Cotterell & Sharpe, 2010).

Given that Turing gene networks contain an extracellular positive loop (i.e. an L^+^ subnetwork), they are, depending on the parameters, either RD-stable or RD-unstable of the first kind (see Fig. 6A-B). In the former case the resulting patterns are periodic. For homogeneous-with-noise initial patterns, a good approximation for the wavenumber of the resulting pattern is given by the unstable wavenumber closest to the maximum of the real part of the principal branch of the dispersion relation (Murray, 2002) (see Fig. 6A-B). For combined spike-homogeneous initial pattern a more detailed analysis is required to understand the wavenumbers in the resulting pattern (Tarumi & Mueller, 1989; Klika *et al*., 2024) but the resulting pattern is also periodic.

In Turing gene networks, the larger wavenumbers are stable (i.e. do not grow) because of the L^+^ subnetwork. The signal in the L^+^ subnetwork diffuses between cells and activates its own production in nearby cells. This produces a homogenizing effect in which, contrarily to what happens in over-Turing networks, the initial differences in concentration between contiguous cells do not necessarily grow over time. In fact, the positive extracellular loop precludes large concentration differences between nearby cells and thus, precludes large unstable wavenumbers (i.e. having many concentrations peaks per unit of space and so, large relative differences in concentration between contiguous cells).

In 1D, the possible resulting patterns of Turing networks are similar to the periodic patterns arising from over-Turing gene networks (see Fig. 9). The difference is that, while in Turing networks these periodic patterns can arise from both homogeneous-with-noise and combined spike-homogeneous initial patterns, in over-Turing networks they can only arise from combined spike-homogeneous initial patterns. In 2D and 3D, Turing networks acting on homogeneous-with-noise initial patterns can lead to resulting patterns consisting of several concentration peaks (e.g., dots) or ridges (e.g., stripes) with a constant spacing and regularly distributed over space, or forming labyrinths (Meinhardt, 1982; Murray, 2002), while over-Turing networks acting on those initial patterns can only produce noisy resulting patterns (see Fig.10). From combined spike-homogeneous initial patterns in 2D and 3D domains, over-Turing networks can lead to resulting patterns consisting of concentrically arranged concentration ridges and basins due to the radial symmetry of the problem (see Fig.10). This is also possible from Turing gene networks (Meinhardt, 1982).

As in the case of over-Turing gene networks, changes in the parameters of a Turing gene network (e.g., in diffusion coefficients) can change the global number, height or width of concentration peaks and valleys in the resulting pattern (Turing, 1952; Murray, 2002; Maini *et al*., 2006; Meinhardt, 2008). However, in contrast to what we see in H networks (Salazar-Ciudad et a., 2000), these parameter changes lead to all concentration peaks and valleys to change in the same way. The reasons for this difference are the same we discussed for over-Turing gene networks (see Fig 17-18) (i.e. a relatively small number of gene products is responsible for the resulting pattern and no part of the network is responsible for any specific subset of concentration peaks or valleys) For this same reason the ensemble of distinct resulting patterns attainable by Turing gene networks is smaller and less diverse than the ensemble of distinct resulting patterns attainable by H gene networks. Simply, the ensemble of possible Turing networks can only produce resulting patterns where all concentration peaks (and valleys) are identical while the ensemble of possible H networks can produce resulting patterns where different concentrations peaks and valleys can have different heights, weights and spacings (see Salazar-Ciudad, *et al*. 2000, 2001 for a more detailed exploration).

Turing subnetworks can be combined into composite gene networks (see Fig.16). By combining Turing subnetworks in series, the resulting pattern of one subnetwork can serve as the initial pattern of the other (Fujita & Kawaguchi, 2013; Moustakas-Verho *et al*., 2014). As long as these combinations the only positive loops are extracellular, the system can only be, depending on the parameters, RD-stable or RD-unstable of the first kind (just like single Turing networks). Consequently, regardless of how complex the composite network is, the possible resulting patterns are still periodic patterns, that may combine several wavelengths (Fujita & Kawaguchi, 2013; Moustakas-Verho *et al*., 2014). Combining Turing gene networks with over-Turing gene networks in series leads to composite gene networks that behave either as RD-unstable networks of the first or second kind depending of model parameters (see section S2 in the appendix).

In general, gene networks that combine Turing gene networks and H gene networks in series have variational properties with features of both the Turing and H gene networks (see Fig.16). There are, in fact, many articles studying this (Salazar-Ciudad *et al*., 2001; Miura, 2013; Green & Sharpe, 2015; Glim *et al*., 2021; Tzika *et al*., 2023). If the Turing networks are upstream of the H networks, then each concentration peak resulting from the Turing network acts as a different spike in the input pattern received by the H network. Thus, the peaks and valleys resulting from the H network can be repeated around each of the concentration peaks produced by the Turing network. If the peaks produced by the H network are wider than those produced by the Turing, however, these latter peaks may fuse and the resulting pattern simply resembles that of the Turing network (see Fig.16).

In gene networks in which H networks are upstream of the Turing networks, the resulting pattern can be periodic, but the H networks can make the periodic pattern slightly different in different spatial regions (by specific gene product in the H networks regulating specific gene products in the Turing networks). Thus, the height or spacing of concentration peaks in the resulting pattern of the Turing network can be larger, or smaller, where patterned genes of the H network are expressed (see Fig.15).

Besides being combined into Turing or over-Turing subnetworks, or their combinations, L^+^ and L^-^ can be combined into any arrangement (e.g. into multiple nested loops). As we have shown in previous sections, however, a gene network with such complex combinations can only be, nonetheless, RD-unstable of the first or second kind. In the former case the resulting pattern will be periodic, as when combining Turing gene networks. In the second case, we do not know for sure which are the possible resulting patterns but since the number of unstable wavenumbers is infinite, we can expect that, from homogeneous patterns with noise, the resulting patterns are amplified noise patterns (just as combining several over-Turing networks). Similarly, from combined spike-homogeneous patterns we expect froze-wave patterns, possibly combining different periods.

## Discussion

The main conclusions of this article are that all gene networks capable of non-trivial pattern transformations can be classified into three distinct topological classes (and their combinations) and that each of these classes allow for qualitatively different types of resulting patterns. For gene networks within each class we found some additional topological requirements.

All gene networks can be decomposed into subnetworks. Because we study pattern transformation by extracellular signaling, we decompose gene networks based on the subnetworks downstream of each of its extracellular signal. These subnetworks are classified based on whether their extracellular signals are positively downstream of themselves (L^+^ subnetworks), negatively downstream of themselves (L^-^ subnetworks) or not downstream of themselves (H subnetworks). This classification exhaust all possibilities, except for subnetworks that arise as a combination of several of these simpler subnetworks.

We have shown that pattern transformation requires gene networks with positive regulatory loops and that, if these loops are extracellular, then the resulting patterns, when heterogeneous, are periodic. We have also shown that, if the positive regulatory loops are intracellular, then the resulting patterns, when heterogeneous, are characterized by an infinite amount of unstable wavenumbers. However, negative interactions are also required for pattern transformation. The combination of the former findings with this latter requirement implies that there are only three types of gene networks leading to pattern transformations: 1) Hierarchical gene networks with positive intracellular loops and negative regulations 2) Over-Turing gene networks (i.e. gene networks with negative extracellular loops and positive intracellular loops forming a loop between them) 3) and Turing gene networks (i.e. gene networks with positive and negative extracellular loops forming a loop between themselves). These three fundamental classes, and their combinations, exhaust all possibilities for gene networks capable of non-trivial pattern transformations through extracellular signaling.

In this article we also show that these three classes of gene networks, and their combinations, each lead to specific types of resulting patterns (see Fig.9-10). Hierarchical networks lead to resulting patterns that are radially symmetric and made of any combination of concentration peaks and valleys (ridges and basins in higher dimensions). Depending on the parameters of the network, these resulting patterns can contain more or less peaks and valleys with different positions, heights and different shapes. In over-Turing gene networks infinitely-many wavenumbers are linearly unstable and thus, many get amplified in the resulting pattern. From homogeneous-with-noise initial patterns, this leads to amplified-noise patterns, while from combined spike-homogeneous patterns, this leads to radially symmetric periodic patterns extending all over the domain. Similarly, we also prove that Turing networks only amplify a finite number of wavenumbers, if any, and then, the only non-trivial pattern transformations they can lead to result in periodic patterns. In the last part of the article, we discuss how the further combination of hierarchical with over-Turing, hierarchical and Turing or Turing and over-Turing networks leads to resulting patterns that can be seen, to a large extent, as the combination of the types of patterns possible from each of the three classes of networks.

It is worth acknowledging that we have used a deterministic, mean-field reaction-diffusion framework, which does not explicitly account for stochastic fluctuations, ubiquitous in any real biological system. These microscopic fluctuations can act as small perturbations everywhere in the domain, including in cases where we have considered a spike-homogeneous initial pattern without noise. The resulting patterns from a spike-homogeneous initial pattern with noise can be understood to a large extent as a combination of those arising from spike-homogeneous initial conditions near the spike (where radial symmetry may be preserved) and homogeneous-with-noise initial conditions (where radial symmetry may be lost) far from it, at least in those cases where the amplitude of the noise is small enough compared to the spike (Wang et al., 2022). Indeed, given that extracellular signals are typically received by membrane receptors that activate intracellular signal transduction pathways leading to changes in transcription, that only happen in the cell nucleus, spatial fluctuations at the scale of the cell membrane get effectively averaged out before influencing extracellular signal production (Berg and Purcell, 1977; Gregor et al., 2007). We therefore expect that the inclusion of stochasticity would not fundamentally alter the main conclusions of our framework, although it may affect the fine details of the resulting patterns (e.g., resulting patterns of hierarchical and over-Turing networks need not preserve the radial symmetry of the spike-homogeneous initial pattern).

Throughout our article, we have considered that cells are arranged in simple regular ways over space. In our arguments, the only element that depends on the geometry of the cellular arrangement is the explicit form of the eigenmodes in (6), whereas the construction of the dispersion relation itself is geometry-independent. The existence of a limited number of topological classes capable of generating patterns does not depend on the geometry of space (i.e., on how cells are distributed within the embryo). The type of pattern transformations for each class does not depend on the geometry of space either (e.g., Turing networks lead to periodic patterns, H networks do not, etc). The geometry of space, however, can change specific aspects of the specific resulting patterns possible from each specific gene network. Thus, for example, the height, spacing and shape of concentration peaks and valleys can change close to the boundaries of the system (Diambra & Costa, 2006; Glimm *et al*., 2014; Castelino *et al*., 2020). In Turing networks it is even possible that the resulting patterns changes substantially depending on the shape of the system (e.g., from producing stripes to producing spots), but in all cases the type of pattern, in the coarse way we have defined, remains periodic (Murray, 2002).

All the gene network topologies capable of pattern transformations that we report have been individually reported before in one way or another (Turing, 1952; Salazar-Ciudad *et al*., 2001; Cotterell *et al*., 2015; Wang, 2022). What has not been shown before is that these are the only possible gene network topologies in non-trivial pattern transformation, given the biologically reasonable restrictions on *f* that we describe. We think this is important both for theory and empirical studies. For theory, it is important because we provide a general description of what is possible at the level of non-trivial pattern transformations and underlying gene network topologies (as long as there is no cell movement). Experimentally, this is also important because it provides a simple guideline for experimental developmental biologists trying to understand pattern transformations in specific organs over a set of developmental stages. Thus, if no cell movements occur during these stages, the underlying gene network should have a network topology and variational properties among the ones described in this article (see Fig.19). This significantly narrows down the range of possible underlying gene networks and patterning mechanisms to consider when trying to understand the development of a multicellular system. Similarly, the results of this article allow to improve the study of the evolution of gene networks since they show that the gene networks capable of non-trivial pattern transformations can only change within a finite set of classes (Salazar-Ciudad *et al*., 2001).

**Figure 19.** Variational properties of the gene network topologies capable of pattern transformation. Network colors and shapes as in Figures 2 and 4.

Of the three classes of gene network topologies we identify, two have been widely studied before: hierarchical and Turing networks. Most of the research in Turing gene networks has been theoretical (Turing, 1952; Maini *et al*., 2006; Meinhardt, 2008), but there is also direct experimental research on the involvement of these gene networks in the development of many organs (e.g., Glover *et al*., 2017; Johnson *et al*., 2023; Tzika *et al*., 2023; Tseng *et al*., 2024, just to name some recent ones), although the underlying networks are usually more complex than the ones studied theoretically. Although hierarchical networks are usually not called this way, they constitute the bulk of the gene networks experimentally studied (Salazar-Ciudad *et al*., 2001; Gilbert and Barresi, 2023) and they are, by far, the easier to understand. This, and the fact that many networks seem hierarchical when only some of their interactions are known (Salazar-Ciudad, 2009), may have biased research to focus on them. Although over-Turing gene networks seem simpler and easier to understand than Turing networks, they have only been discovered very recently and only based on theoretical work (Cotterell *et al*., 2015; Marcon *et al*., 2016; Wang *et al*., 2022). It is thus, an open and suggestive question whether this class of gene networks is widely used in pattern transformations in multicellular systems. In fact, it is perfectly possible that organs that are though to use Turing networks may actually be using over-Turing networks instead, since the latter produce resulting patterns that are similar to the former, especially when the organ is effectively 1D.

## Acknowledgements

We thank H. Cano-Fernández, S. Greendal, C. Hedges and A. Loreto-Velázquez for their comments. We are also grateful to the Spanish Ministry of Science and Innovation for funding PID2021-122930NB-100 and CNS2022-135397 to I.S.-C. (MCIN/AEI/10.13039/501100011033/FEDER, UE) ; and to the Catalan Office of Universities and Research for 2022FI_B 00610 to K.M.-A. This work was likewise supported by the Spanish State Research Agency, through the Severo Ochoa and María de Maeztu Program for Centers and Units of Excellence in R&D (CEX2020-001084-M). We thank CERCA Programme/Generalitat de Catalunya for institutional support.

## Appendix

### S1: Formal definition of pattern transformation

Let us consider an arbitrary gene regulatory network, and let *g*_*i*_ (*t, x*) denote the concentration of gene product *i* at time *t* ≥ 0 and position *x* in an *n*-dimensional spatial domain, with *n*=1, 2, 3. At the beginning of the patterning process, gene product *i* exhibits an initial pattern given by 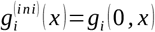. At the end of the patterning process, given requirement R2 in the main text, we know that the concentration of gene product *i* converges to a stationary resulting pattern that we denote by 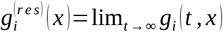. For the purpose of this article, we ignore non-stationary solutions to the model equations (e.g., time-periodic molecular clocks within cells).

In this framework, we say the gene product *i* undergoes a pattern transformation if its concentration changes during the patterning process at at least one position, that is,

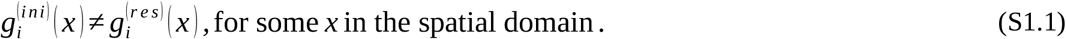

Moreover, we say that gene product *i* undergoes a heterogeneous pattern transformation if the resulting pattern at the end of the patterning process is heterogeneous, that is,

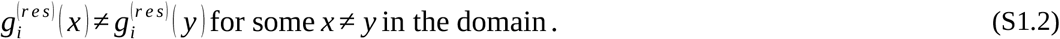

We say that gene product *i* undergoes a rearranging pattern transformation if the position of some critical point of its concentration changes along the patterning process (e.g., the emergence of a new maximum of concentration). If 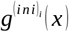 and 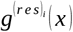 are both sufficiently smooth, this means that either,

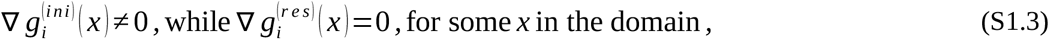

indicating the emergence of a new critical point in the resulting pattern 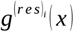; or,

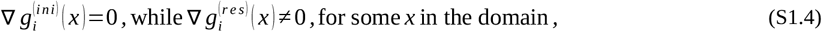

indicating the disappearance of a critical point from the initial pattern 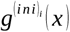. However, none of the three basic initial patterns that we take into consideration in this article is even continuous in space. In this sense, we consider that 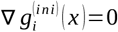 for all *x* in the domain, if 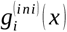 is a noise-homogeneous initial pattern; and we consider 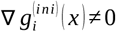 for all *x* ≠ *x*_*c*_ and 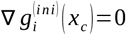0, if 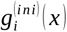 is a spike or spike-homogeneous initial pattern centered at some position *x*_*c*_. Formally, this two *ad hoc* assumptions correspond to the gradient of the convolution of 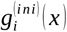 with the Gaussian kernel of diffusion for a brief time [0, *τ*), with *τ* >0 arbitrarily small.

Finally, we say that gene product *i* undergoes a non-replicating pattern transformation if the spatial arrangement of critical points in its resulting pattern does not coincide with the spatial arrangement of critical points in the initial pattern of any other gene product (i.e., any new piece of spatial information in the resulting pattern of *i* must be created during the patterning process and not be replicated from any initial pattern). This means that, for any gene product *j* ≠*i* in the gene network, either,

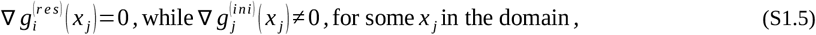

indicating that 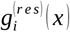 exhibits a critical point at a position where 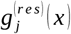 does not; or,

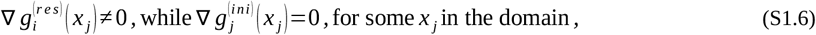

indicating that 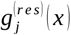 exhibits a critical point at a position where 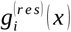 does not. Of course, the same considerations about the non-smoothness of initial conditions in conditions (S1.3) and (S1.4) must also be taken into consideration for (S1.5) and (S1.6).

All in all, for the purposes of this article, we say that gene product *i* undergoes a non-trivial pattern transformation if it undergoes a heterogeneous, rearranging and non-replicating pattern transformation, that is, if 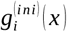 and 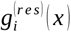 satisfy conditions (S1.1) to (S1.6). Condition (S1.2), together with the fact that the resulting pattern must be stationary, correspond to requirement P1 in the main text. Conditions (S1.3) to (S1.6) correspond to requirement P2 in the main text.

### S2: Positive regulatory loops and RD-instability

For a gene network to be RD-unstable of any kind it needs to have at least one positive regulatory loop: if all positive loops are all extracellular, then the gene network is RD-unstable of the first kind; if positive loops are all intracellular, then the gene network is RD-unstable of the second kind; and if the network has positive extracellular and intracellular loops, the type of RD-instability will depend on the particular choice of model parameters. In the main text, we provided proves for these statements in the case where the gene network displays only one regulatory loop, and this loop is positive. Here, we complete such results with a general proof in the case any number of positive and negative loops (i.e., for any general gene regulatory network).

Let 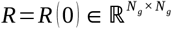 denote the reaction-diffusion matrix of a given gene network in the absence of diffusion. Given a set 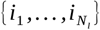 of *N*_*l*_ ≤ *N*_*g*_ gene products, the principal submatrix of *R* associated to 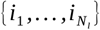 corresponds to the *N*_*l*_ *× N*_*l*_ -matrix 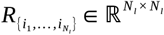 resulting from considering only those entries in *R* whose row and column indices are given by 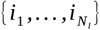 (i.e., considering only regulations between the gene products in 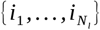. In this sense, the principal minor 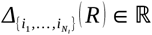 associated to 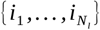 is the determinant of the corresponding principal submatrix of *R*, that is,

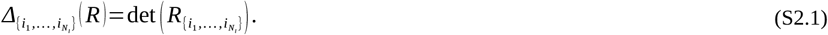

If no gene products in 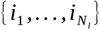 form a regulatory loop, the corresponding principal minor in (S2.1) reduces to the product of the diagonal entries in 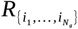, namely,

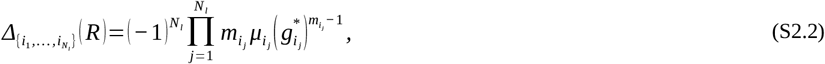

where 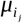 is the degradation coefficient of gene product 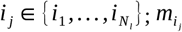 is the order of such degradation; and 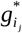 the corresponding steady-state concentration around which *R* was constructed (see Equation (14) in the main text). If 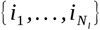 form one, and only one, regulatory loop, the corresponding principal minor is given by,

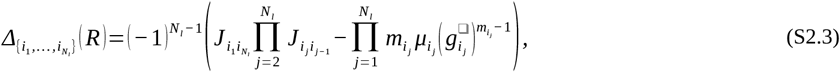

where 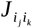 represents the regulation that gene product 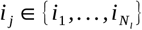 exerts of gene product 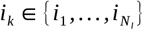 (see Equation (19) in the main text). If 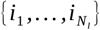 form more than one regulatory loop, each with a different length, more convoluted expressions than (S2.3) can be found using Leibniz formula (Strang, 2016).

Thus, in the case of regulatory loops, principal minors measure the net effect that the corresponding regulatory loop has on reaction-diffusion. For example, if we consider a direct self-regulatory loop (i.e., *N*_*l*_=1), we get,

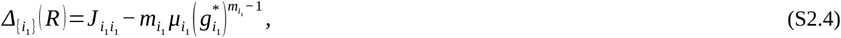

with 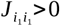 for a positive loop, and 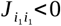 for a negative one. Then, for any choice of model parameters for which 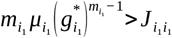, we have that the degradation overpowers any positive effect of the self-regulatory loop and then, any concentration perturbation in *i*_1_ would decay back to the steady-state concentration 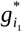 (i.e., perturbations are stable). However, if for some choice of model parameters 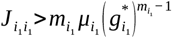, then concentration perturbations in *i*_1_ wouls grow in time (i.e., perturbations are unstable).

With this in mind, we say that a principal minor 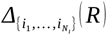 of the reaction-diffusion matrix *R* is a destabilizing minor if there exists some choice of model parameters such that,

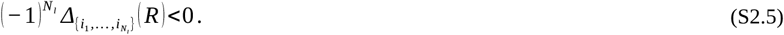

The following result shows that the principal minor associated to any set of gene products 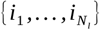 that forms a positive regulatory loop is indeed a destabilizing minor:

#### Lemma S2.1.

*Let us consider a gene regulatory network with N*_*g*_ ≥ 1 *gene products, and let* 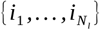, *with N*_*l*_ ≤ *N*_*g*_, *form a positive regulatory loop; then, the principal minor of R associated to* 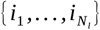 *is a destabilizing minor of R*.

**Proof (of Lemma S2.1).-** Without loss of generality, we assume that the *N*_*l*_ gene products in the positive loop are labeled {1, …, *N* } (otherwise, we can simply reorder the labels in the gene network without altering its network topology). Moreover, for the sake of simplicity, we denote by 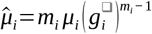 the whole degradation term corresponding to gene product *i*∈ {1, …, *N*_*l*_ }.

The principal submatrix of *R* associated to the positive loop {1, …, *N*_*l*_ } is given by,

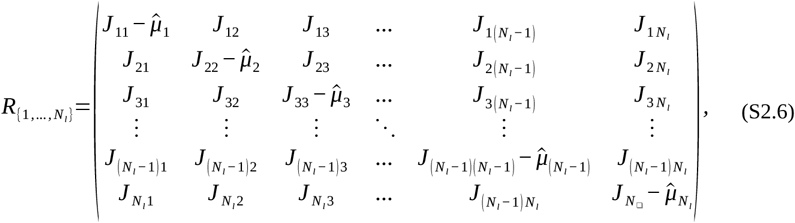

where the bold entries denote the regulations of the positive regulatory loop formed by {1, …, *N*_*l*_ } (i.e., the regulation of 1 on 2, of 2 on 3, and so forth). By definition of positive loop, this entries represent a set of non-zero entries of 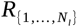 whose product is strictly positive (see Figure 4 in the main text). The remaining entries in (S2.6) correspond to other possible regulations between gene products {1, …, *N*_*l*_ } that do not belong to the main loop and hence, they can all be equal to zero (e.g., Equation (16) in the main text), or not.

Using Leibniz formula (Strang, 2016) for the determinant of (S2.6), we can write the principal minor of *R* associated {1, …, *N*_*l*_ } as a sum of the form,

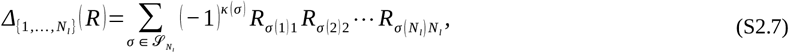

where *R*_*ij*_ ∈ℝ are the corresponding entries of 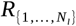 (i.e., 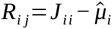; and *R*_*ij*_=*J*_*ij*_, if *j* ≠*i*); 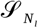 is the symmetric group of all possible permutations *σ* of the *N*_*l*_-element set {1, …, *N*_*l*_ } (Hungerford, 1974); and, *κ* (*σ*) is the number of inversions made by made by *σ*, that is, the number of ordered pairs (*i, j*), with *i*< *j*, for which *σ* (*i*)>*σ* (*j*).

In this sense, the product of all bold entries in (S2.6) corresponds to the term in (S2.7) defined by the cyclic permutation,

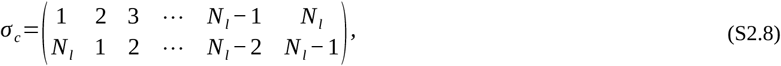

where each element in {1, …, *N*_*l*_ } (i.e., first row of (S2.8)) permutes with the element strictly to its left (i.e., second row of (S2.8)), except for 1, that permutes with *N*_*l*_. As we can see, *σ*_*c*_ has *N*_*l*_ − 1 inversions given by pairs (1, *i*) for all *i*∈ {2, …, *N*_*l*_ } (indeed, 1<*i* while *σ*_*c*_ (1)=*N*_*l*_>*σ*_*c*_ (*i*)). Consequently, *κ* (*σ*_*c*_)=*N*_*l*_ − 1, and then, the principal minor in (S2.7) can be written as,

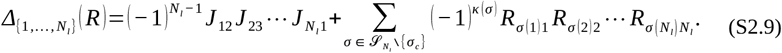

Then, multiplying both sides of (S2.9) by a factor of 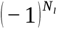, we get,

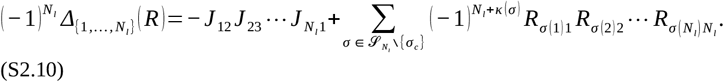

As we mentioned before, the product 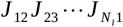 is strictly positive by definition and hence, the first term in (S2.10) is strictly negative. Consequently, regardless of the sign of the second term, there always exists some choice of model parameters for which (S2.10) becomes negative, namely,

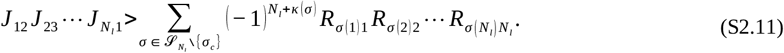

This means, according to (S2.5), that 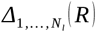 is a destabilizing minor and thus, the proof of Lemma S2.1 is finished.

#### Positive intracellular loops induce RD-instabilities of the second kind

Let us consider a gene regulatory network with *N*_*ic*_ ≥ 1 intrcellular gene products and *N*_*ec*_=*N*_*g*_ − *N*_*ic*_ extracellular signals, and let 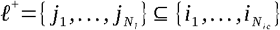 form a positive intracellular loop, with *N*_*l*_ ≤ *N*_*ic*_. In accordance with Lemma S2.1, we know that the principal minor of *R*=*R* (0) associated to *ℓ*^+^ is a destabilizing minor, as it corresponds to a positive regulatory loop. The following results extends this property to the principal minor of *R* associated to the whole set of intracellular gene products in the network (i.e., from 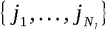 to 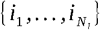:

##### Lemma S2.2.

*Let us consider a gene regulatory network with N*_*g*_>1 *gene products and let* 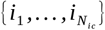 *denote its N*_*ic*_< *N*_*g*_ *intracellular gene products. If such gene network exhibits at least one positive intracellular loop* 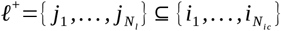, *with N*_*l*_ ≤ *N*_*g*_ ; *then, the principal minor of R*=*R* (0) *associated to* 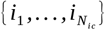 *is a destabilizing minor of R*.

**Proof (of Lemma S2.2).-** Without loss of generality, we can assume that the *N*_*ic*_ intracellular gene products in the network are labeled {1, …, *N*_*ic*_ }, with 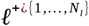 (otherwise, we can reorder the labels in the gene network without changing its network topology). Moreover, for the sake of simplicity, we denote 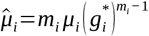 the degradation term corresponding to gene product *i*∈ {1, …, *N*_*ic*_ }.

From Lemma S2.1, we know that there exists some choice of model parameters such that 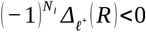0. Hence, if 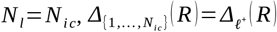 is trivially a destabilizing minor of *R*. If *N*_*l*_ < *N*_*i c*_, let *i*∈ {*N*_*l*_ +1, …, *N*_*i c*_ } be any intracellular gene product outside of *ℓ*^+^. Then, using the minor decomposition formula (Strang, 2016), we can write 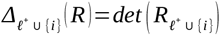 in the form,

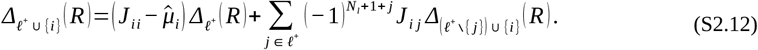

If we now multiply both sides of (S2.12) by a factor of 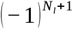, we get,

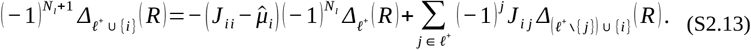

However, we know that there exists some choice of model parameters for which 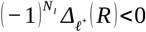 and then, complementing such choice with a set of model parameters for gene product *i* (namely, *J*_*ii*_, 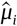 and *J*_*ij*_ for all *j*∈*ℓ*^+^) such that,

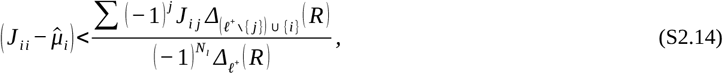

we get 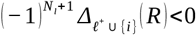 (note that the sign of the terms in the top sum of the right-hand side of (S2.14) alternates depending on the parity of *j*). This means, according to (S2.5), that the principal minor 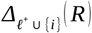 is a destabilizing minor of *R*. Then, the same argument can be iteratively applied to the remaining gene products outside of *ℓ*^+^ ∪{*i* } (i.e., gene products *j* ∈{*N*_*l*_+1, …, *N*_*ic*_ }\ {*i* }) until we end up showing that 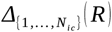 is indeed a destabilizing minor. This would finish the proof of Lemma S2.2.

So far, parameter choices involved in Lemmas S2.1 and S2.2 correspond exclusively to the reaction part of the reaction-diffusion equations (namely, Jacobian entries *J*_*ij*_, degradation coefficients *μ*_*i*_, degradation orders *m*_*i*_ and steady-state concentrations 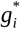). Nevertheless, to understand the true role of positive loops in pattern transformation we must examine the effect that destabilizing minors of *R*=*R* (0) have on the full reaction-diffusion matrix *R* (*k*^2^)=*R*−*k*^2^ *D*, for all wavenumbers *k* ∈ *σ* (∇^2^). To do so, given that *D* is by definition a positive semidefinite diagonal matrix, we can write the determinant of *R* (*k*^2^) in terms of the principal minors of *R* as follows (Wang & Li, 2001),

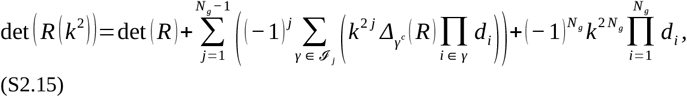

where *J* _*j*_={*γ* ={*i*_1_, …, *i* _*j*_ }/1 ≤*i*_1_ ≤ … ≤*i* _*j*_ ≤ *N*_*g*_ } denotes the set of all *j*-element index sequences extracted from {1, …, *N*_*q*_ }; and *γ*^*c*^={1, …, *N* }\ *γ* is the complementary sequence of *γ* ∈ *J* _*j*_.

The following result shows that any gene network containing at least one positive intracellular loop admits at least one choice of model parameters (including diffusion coefficients in *D*) for which it becomes an RD-unstable network of the second kind:

##### Proposition S2.3.

*Let us consider a gene regulatory network with N*_*ic*_ ≥ 1 *intracellular gene products and N*_*ec*_=*N*_*g*_ − *N*_*ic*_ ≥ 1 *extracellular signals. If the network contains at least one positive intracellular loop* 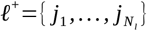, *with* 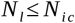; *then, there exists a choice of model parameters for which the gene network becomes an RD-unstable network of the second kind*.

**Proof (of Proposition S2.3).-** Without loss of generality, we can assume that the *N*_*ic*_ intracellular gene products in the network are labeled {1, …, *N*_*ic*_ }, with *ℓ*^+^={1, …, *N*_*l*_ } (otherwise, we can reorder the labels in the gene network without changing its network topology). Moreover, for the sake of simplicity, we denote by 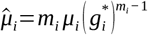 the full degradation term corresponding to gene product *i*∈ {1, …, *N*_*g*_ }.

From Lemmas S2.1 and S2.2, we know that the principal minor of *R*=*R* (0) associated to the full set of intracellular gene products {1, …, *N*_*ic*_ } is a destabilizing minor and consequently, that there exists a choice of model parameters for which,

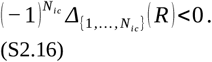

First, we will focus on proving that there exists a choice of model parameters for which the gene network is RD-unstable, that is, a choice of model parameters for which there exists at least one unstable wavenumber *k* ∈*σ* (∇^2^) for which its associated dispersion relation det (*λ I* − *R* (*k*^2^))=0 admits one solution with positive real part. In this sense, according to the Routh-Hurwitz conditions (Kurosh, 1984; Elsgolts, 2003), det (*λ I* − *R* (*k*^2^))=0 will admit a solution with positive real part if 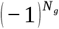 det (*R* (*k*^2^))<0. Hence, it suffices to show that there exists some choice of model parameters for which this latter inequality holds to prove that the corresponding gene network is RD-unstable for such choice of model parameters.

Let 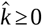 be a dummy continuous wavenumber variable. The determinant of 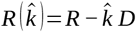 can be written in terms of the minors of *R* using (S2.15). However, since we have *d*_*i*_=0 for all *i*∈ {1, …, *N*_*ic*_ }, such formula reduces to,

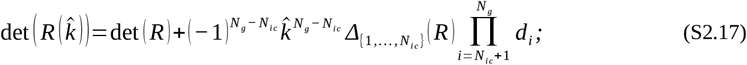

and then, if we multiply both sides of (S2.17) by a factor of 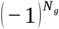, we get,

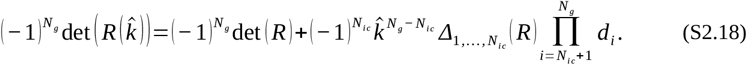

According to (S2.16), there exists some choice of model parameters for which 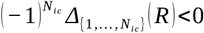. Thus, given that 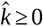 and *d*_*i*_ >0, for all *i*∈ {*N*_*ic*_ +1, …, *N*_*g*_ }, there exists some threshold wavenumber 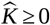 such that 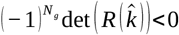 for all 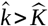, namely,

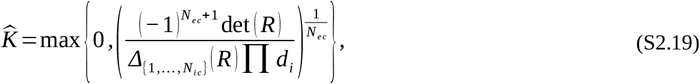

where *N*_*ec*_=*N*_*g*_ − *N*_*ic*_ ≥ 1 is the number of extracellular signals in the network. The argument is then finished by noting that *σ* (∇^2^) is a countably, infinite and unbounded set (Baker *et al*., 2008), so that we can always find some wavenumber *k* ∈*σ* (∇^2^) such that 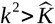, which ensures 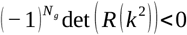. In other words, for any choice of model parameters enabling (S2.16) (whose existence is given by Lemma S2.2), the gene regulatory network admits at least one unstable wavenumber *k* ∈*σ* (∇^2^) and is, therefore, RD-unstable.

Finally, given that *σ* (∇^2^) is an unbounded set, for any unstable wavenumber *k*_1_ ∈*σ* (∇^2^) with 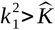, there always exists some other wavenumber *k*_2_ ∈*σ* (∇^2^) with 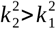. This means that 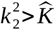 and ultimately, that 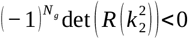, implying that *k*_2_ is also an unstable wavenumber. As a result, the gene regulatory network does not only admit one, but infinitely many unstable wavenumbers, that is, it is an RD-unstable network of the second kind. This finishes the proof of Proposition S2.3.

#### Positive extracellular loops induce RD-instabilities of the first kind

Let us consider a gene regulatory network with *N*_*ec*_ ≥ 1 extracellular signals and *N*_*ic*_=*N*_*g*_ − *N*_*ec*_ ≥ 0 intracellular gene products, and let 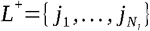 form the largest positive extracellular loop in the network (i.e., at least one gene product *j*_*i*_ ∈ *L*^+^ is an extracellular signal).

To isolate, and better understand, the role of positive extracellular loops in pattern transformation, we will assume for the remainder of this section that the gene networks under consideration do not contain any positive intracellular loop (the combination of positive extracellular and intracellular loops is considered in the following section). In particular, given this lack of positive intracellular loops in the network, we will have that the principal minor associated to the set of intracellular gene products 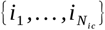 is never a destabilizing minor, that is, for any choice of model parameters, we have,

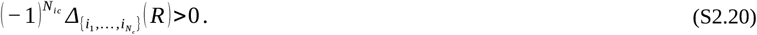

The following results shows that, if a gene regulatory network whose positive loops are all extracellular becomes RD-unstable for some choice of model parameters, then it can only be an RD-unstable network of the first kind (i.e., its associated dispersion relation would yield only a finite number of unstable wavenumbers *k* ∈*σ* (∇^2^)):

##### Proposition S2.4.

*Let us consider a gene regulatory network with N*_*ec*_ ≥ 1 *extracellular signals and N*_*ic*_=*N*_*g*_ − *N*_*ec*_ ≥ 0 *intracellular gene products, and assume all positive loops in the network are extracellular, with* 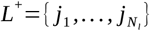 *denoting the largest one; then, if there exists some choiceof model parameters for which the network becomes RD-unstable, it can only be an RD-unstable network of the first kind*.

**Proof (of Proposition S2.4).-** Let 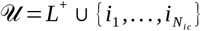 be the union set of loop *L*^+^ with the set of all intracellular gene products in the network 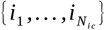, and let *M* =|*U*| be the number of elements in *U* . Without loss of generality, we can assume that the *M* gene products in *U* are labeled {1, …, *M* }, with intracellular gene products being {1, …, *N*_*ic*_ } (otherwise, we can always reorder the labels in the gene network without changing its network topology). Moreover, for the sake of simplicity, we denote by 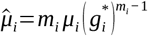 the full degradation term corresponding to gene product *i*∈ {1, …, *N*_*g*_ }.

From Lemma S2.1, we know that the principal minor of *R*=*R* (0) associated to *L*^+^ is a destabilizing minor (i.e., there exists some choice of model parameters for which 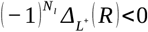. Moreover, proceeding as in Lemma S2.2, we can extend such property to the principal minor associated to *U*, namely, we can find a choice of model parameters for which,

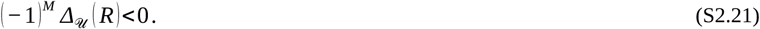

First, we will focus on showing that the positive extracellular loop *L*^+^ enables the existence of some choice of model parameters for which the network is RD-unstable (i.e., for which its associated dispersion relation yields at least one unstable wavenumber *k* ∈*σ* (∇^2^)). As in the proof of Proposition S2.3, we will do it using the Ruth-Hurwitz conditions (Kurosh, 1984; Elsgots, 2003) to ensure that det (*λ I* − *R* (*k*^2^))=0 has at least one solution with positive real part by ensuring that 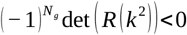.

Let 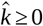 be a dummy continuous wavenumber variable, and let us take *d*_*i*_=*δ* >0 for all extracellular signal *i*∈ {*N*_*ic*_ +1, …, *N*_*g*_ }. The determinant of 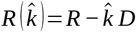 can be written in terms of the minors of *R* using formula (S2.15). Indeed, if *δ* is small enough, we have,

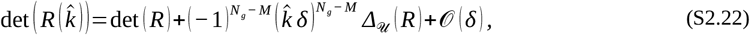

where *O* (*δ*) represents Landau’s big-O notation. Then, if we multiply both sides of (S2.22) by a factor of 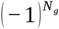, we get,

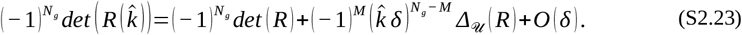

According to (S2.21), there exists some choice of model parameters for which (− 1)^*M*^ *Δ* (*R*)<0. Hence, if *δ* is small enough, it is possible to find some lower bound 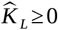 such that 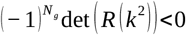 for some wavenumber *k* ∈ *σ* (∇^2^) with 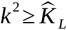, namely,

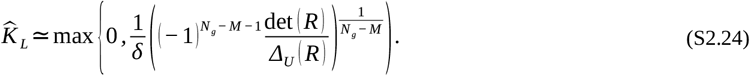

Nonetheless, in contrast to Proposition S2.3, the squared wavenumber *k*^2^ cannot be taken arbitrarily bigger than 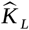 as higher order terms within *O* (*δ*) may become relevant and alter the sign of 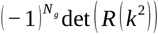. To illustrate this, let us extract the term of order 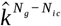 from *O* (*δ*) in (S2.23) and get,

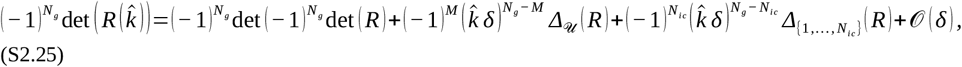

where, in virtue of (S2.20), 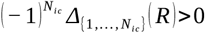 for any choice of model parameters. Thus, given that *M* > *N*_*ic*_, we have that *δ* can always be taken small enough so that the third term in (S2.25) is negligible with respect to the second one. However, for any fixed value of *δ*, this third term (with order 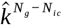 and positive sign) will dominate over the second term (with order 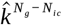 and negative sign) if we let 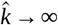. In particular, this means that one can always find some upper limit 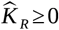 such that,

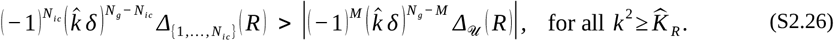

Notice that it can be the case that the interval 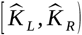 is so narrow that no wavenumber *k* ∈*σ* (∇^2^) satisfies 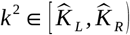. In this case, the gene network under consideration would not be RD-unstable. However, if the interval is wide enough to admit at least one wavenumber *k* ∈*σ* (∇^2^) such that 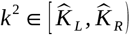 (i.e., if the gene network is RD-unstable), then there can only be a finite number of such wavenumbers: *σ* (∇^2^) is an unbounded set (Baker *et al*., 2008) and hence, there will always be some big wavenumber *k* ∈*σ* (∇^2^) for which 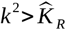. This means that, if the gene network is RD-unstable for some choice of model parameters, it can only be RD-unstable of the first kind and thus, the proof of Proposition S2.4 is finished.

Finally, two observations are worth being made. First, note that tuning model parameters to get an RD-unstable network of the first kind generally requires a much finer degree of adjustment than that needed to get an RD-unstable network of the second kind. In the former case, one must find some wavenumber *k* ∈ *σ* (∇^2^) such that 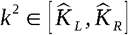, where 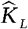 and 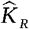 are given in the proof of Proposition S2.4. Conversely, in the latter case it suffices to select a sufficiently large *k* ∈ *σ* (∇^2^) such that 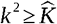, with 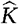 given by (S2.19).

On the other hand, it is worth recalling that although the parameter selection employed for the proof of Proposition S2.4 may appear rather artificial, instances of RD-unstable networks of the first kind arising from less convoluted parameter choices have been reported both in this article (e.g., Figures 8 and 9 in the main text), and in the literature (e.g., Gierer & Meinhardt, 1972; Murray, 2003; Levine & Rappel, 2005).

#### Combination of positive extracellular and intracellular loops

Let us consider a gene regulatory network with *N*_*ic*_ ≥ 1 intracellular gene products and *N*_*ec*_ =*N*_*g*_ − *N*_*ic*_ ≥ 1 extracellular signals, and let 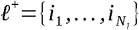 and 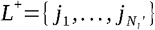 represent two positive intracellular and extracellular loops, respectively. Given that the network contains at least one positive intracellular loop, Proposition S2.3 guarantees the existence of some choice of model parameters for which the network becomes RD-unstable of the second kind. In such parameter set, the intracellular loop *ℓ*^+^ is effectively overweighted relative to the extracellular loop *L*^+^, thereby exerting a stronger influence on the corresponding pattern transformation.

However, model parameters may alternatively be tuned so as to satisfy 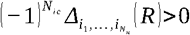. In this case, Proposition S2.4 ensures that this parameter set can be completed to a choice of model parameters for which the network becomes RD-unstable of the first kind. Under this conditions, the extracellular loop *L*^+^ becomes overweighted relative to the intracellular loop *ℓ*^+^, and consequently dominates the reaction-diffusion dynamics, leading to a bigger influence on the corresponding pattern transformation.

Thus, gene regulatory networks that combine positive extracellular and intracellular loops can exhibit RD-instabilities of either types, depending on the specific choice of model parameters. As a result, such networks have access to the full spectrum of pattern transformations associated with both classes of RD-unstable networks.

### S3: Gradients of extracellular signals from spikes

In this section, we analyze the pattern transformations produced by a simple two-node gene network consisting of a self-activating intracellular gene product, denoted 1, that activates an extracellular signal, denoted *A*, when acting on a 1D spike initial pattern. As described in the main text, extracellular signals in this situation are secreted from the spike and form exponential gradients whose slopes depend on model parameters such as their diffusion and degradation coefficient. This subsection therefore provides a detailed derivation of Equation (23) in the main text.

We begin by solving for a stationary concentration of intracellular gene product 1. The concentration dynamics of this intracellular gene product for each *x* is given by the following initial value problem (recall Equation (1) in the main text),

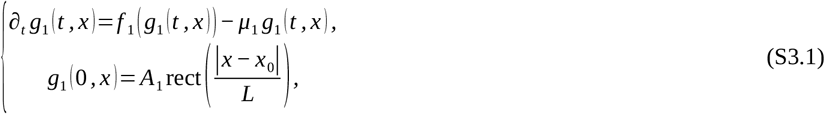

where *A*_1_ >0 is the small amplitude of the spike, *L*>0 is its width, and rect (*y*) denotes the rectangular functions, that is, rect (*y*)=1, if − 1 ≤ *y* ≤ 1, and rect (*y*)=0, otherwise (Abramowitz & Stegun, 1964). Without loss of generality, we can assume that the spike is centered at the origin *x*_0_=0 (otherwise, we can simply consider a spatial translation of the problem).

For any position outside the spike (i.e., *x* ∉ [− *L, L*]), we have *g*_1_ (*t, x*)=0 for all *t* >0, since no production of gene product 1 can ever occur without any non-zero initial concentration and 1 is regulated by no extracellular signal. On the contrary, for any position inside the spike *x* ∈ [− *L, L*], we have, up to a linear level,

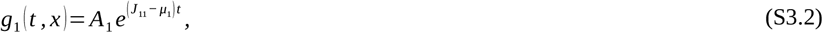

where 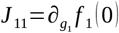. Then, if *μ*_1_ < *J* _11_, the concentration of gene product 1 inside the spike will grow away from 0 until it eventually saturates at some *G*_1_ >0 due to the effect of the non-linear terms of *f* _1_ (*g*_1_) (recall requirements R2 and R4 in the main text). As a result, the resulting pattern of gene product 1 is a spike centered at the same position and a different height,

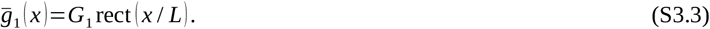

Now, we turn to the analysis of pattern transformations in extracellular signal *A*. If we assume that gene product 1 reaches its stationary concentration fast enough, the concentration dynamics of signal *A* is given by (recall Equation (1) in the main text),

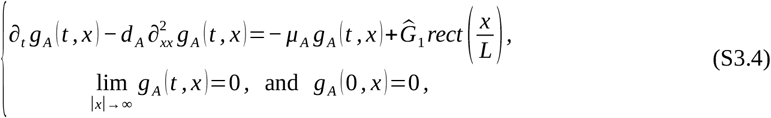

where 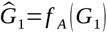. Solutions to a partial differential equation like (S3.4) can be written in the form 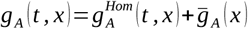 (Sneddon, 1957), where 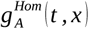 is a solution to the transient homogeneous equation,

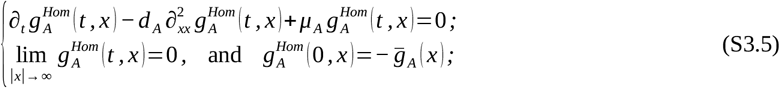

and 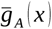 is a solution to the stationary equation,

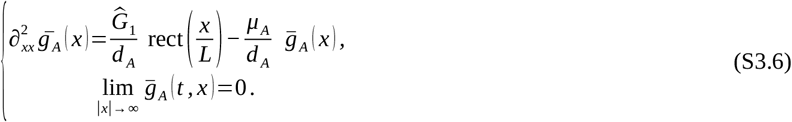

However, for the purposes of this article we are just interested in the formation of stationary solutions of system (S3.4) and thus, we can assume that 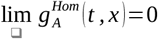 and ignore (S3.5). Regarding the stationary equation, note that (S3.6) is invariant to sign reflection and so, 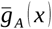 must be an even function (i.e., 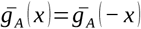). Consequently, we must have 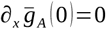, and we can restrict our search for a closed form of 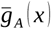 to only the right half-line (i.e., *x* ≥ 0) by adding the extra zero flux boundary condition 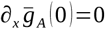 to equation (S3.6).

In this sense, for positions within the spike (i.e., 0 ≤ *x* ≤ *L*), *rect* (*x* / *L*)=1 and the stationary equation (S3.6) reads,

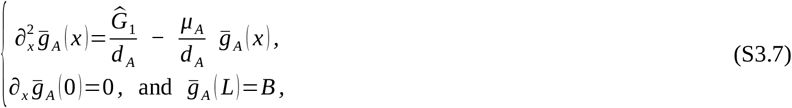

where *B*∈ℝ is a splicing constant that we must later find. Solutions to (S3.7) are given by downward-facing hyperbolic cosines of the form,

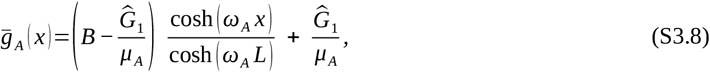

where 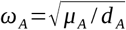. On the other hand, for positions outside the spike (i.e., *x* > *L*), *rect* (*x* / *L*)=0 and then, equation (S3.6) reduces to,

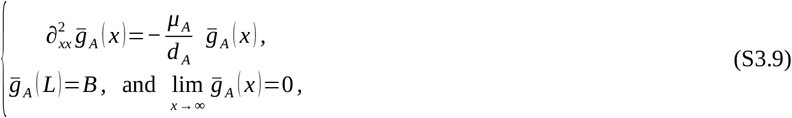

where *B* is the same splicing constant as in (S3.7). Then, solutions to (S3.9) are given by a negative exponential of the form,

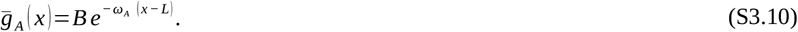

By definition, solutions (S3.8) and (S3.10) match continuously at *x*=*L*. However, we can push this matching to be not only continuous but also continuously differentiable (i.e., class *C* ^1^) by equating the first derivative of both functions at *x*=*L*, and solving for a closed form of the splicing constant *B*. Hence, after some algebra, we get,

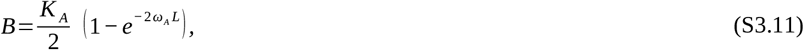

where 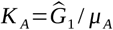. Finally, if we substitute (S3.11) back into (S3.8) and (S3.10), we get that the stationary concentration of extracellular signal *A*=2 or 3 is given by,

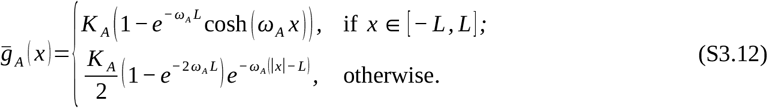

As we can see, the stationary concentration of extracellular signal *A* exhibits a unique, symmetric concentration peak centered at the initial spike (*x*_0_=0 in our case), and two gradients of exponential decay on both sides of the spike. The steepness of such gradients is controlled by parameter *ω*_*A*_ (i.e., the square root of the ratio between the degradation rate and the diffusion rate) in the sense that the smaller *ω*_*A*_ is, the steeper the corresponding gradient. Similarly, constant *C* in Equation (23) of the main text corresponds to,

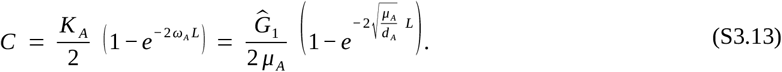

Here, we focus exclusively on the 1D pattern transformations in signal *A*. However, equation (S3.4) corresponds to the classical Helmholtz equation, whose solutions in higher dimensions are well known and extensively documented in the literature (Sommerfeld, 1949; Stakgold & Holst, 2011). Hence, we know that in 2D the signal concentration gradient outside the spike in (S3.12) decays as 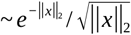, where ||*x*||_2_ denotes the Euclidean norm; whereas in 3D, the corresponding gradient decays as 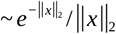.

### S4. Pattern transformations in diamond gene networks for spike initial patterns

In this section, we discuss some details of the 1D pattern transformations that a diamond gene network is able to produce when acting on spike initial pattern. We focus on the diamond network because its topology is simple enough to allow for a precise analysis of the pattern-formation process, while remaining versatile enough to serve as a representative example of many of the features exhibited by more general hierarchical networks.

By definition, a diamond network consists of five distinct gene products (see Figure 11 in the main text). Specifically, it comprises an upstream intracellular gene product (denoted 1) that self-activates and activates two different extracellular signals (denoted 2 and 3). These signals regulate a common downstream intracellular gene product (denoted 4) in such a way that 2 activates 4 whereas 3 inhibits it. Finally, a fifth intracellular gene product (denoted 5) is directly activated by gene product 4. Moreover, we will assume that all gene products suffer from linear degradation (i.e., *m*_*i*_=1 for all *i*∈ {1, 2, 3, 4, 5 }).

As we saw in the previous section, extracellular signals 2 and 3 develop a resulting pattern consisting of a unique concentration peak centered at the spike and with exponential decay. Although this does not constitute a non-trivial pattern transformation (it fails to meet requirements (S1.5) and (S1.6)), the fact that the resulting pattern is heterogeneous, and spreads along the domain due to signal diffusion, allows the emergence of non-trivial pattern transformations in downstream gene products, namely, 4 and 5.

In this sense, the concentration dynamics of intracellular gene product 4 for each *x* is given by the initial value problem (recall equation (1) in the main text),

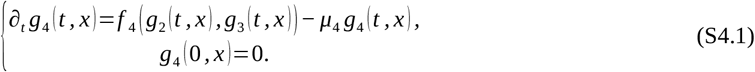

Taking the time derivative in (S4.1) equal to zero, we can easily see that the stationary concentration of gene product 4 is given by,

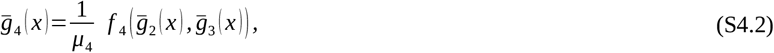

where we have assumed that both extracellular signals reach their stationary concentrations 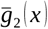 and 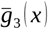 in (S3.12) fast enough. From requirement R5 in the main text, we know that *f* _4_ (*g*_2_, *g*_3_) is monotonously increasing with respect to *g*_2_ and monotonously decreasing with respect to *g*_3_. Then, if we focus on positions to the right of the spike (i.e., *x* > *L*) where both 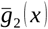 and 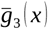 display an exponential gradient, model parameters can be tuned so that the inhibition of 4 by 3 is larger than its activation by 2 close to the spike, and smaller further away. As a result, a concentration peak in the stationary concentration of 4 will emerge at some position 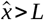 (and, by symmetry of (S3.12), also at 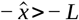).

To illustrate this, let us consider the simple case in which *f* _4_ (*g* _*A*_, *g*_*B*_)=*W* _24_ *g*_2_ −*W* _34_ *g*_3_ is a linear function where *W* _24_, *W* _34_ >0 represent the weights of the regulations that 2 and 3 exert on 4, respectively. In this scenario, (S4.2) reduces to,

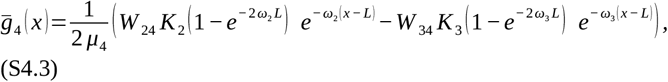

for all positions to the right of the spike (i.e., *x* > *L*). Hence, if we take the first spatial derivative of (S4.3), we get,

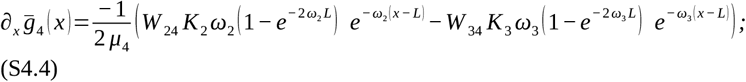

and thus, taking 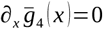 in (S4.4), we can derive a closed form for a unique critical point of 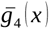 on the right-hand side of the initial spike given by,

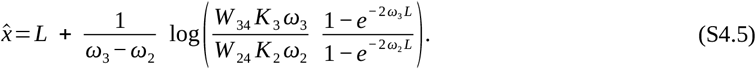

However, this critical point is only formal, as its existence relies on a choice of model parameters for which *x*^ > *L*. In this sense, after some computations on (S4.5), we can see that this is the case for any parameter set that satisfies,

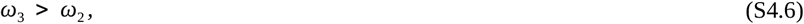

so that the fraction in the second term of (S4.5) is positive; and,

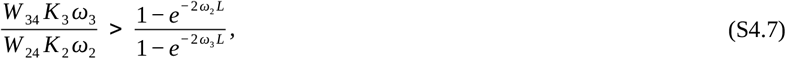

so that the logarithm in the second term of (S3.18) is also positive. Then, if conditions (S4.6) and (S4.7) hold, we have 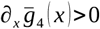 for all positions between *x*=*L* and 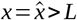; while 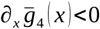 for all positions larger than 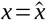. Consequently, 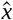 is a concentration peak of 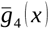 and thus, we can conclude that gene product undergoes a non-trivial pattern transformation given by the emergence of such concentration peak to the right of the initial spike. In fact, given the parity of equation (S3.12),we know that a symmetric concentration peak of 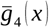 also emerges at 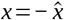, to the left of the spike.

Finally, the concentration dynamics of intracellular gene product 5 for each position *x* is given by the initial value problem (recall equation (1) in the main text),

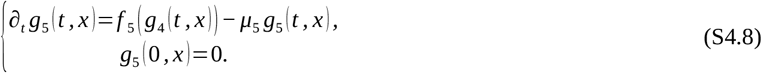

Then, if we set the time derivative in (S4.8) equal to zero, we can easily see that the stationary concentration of gene product 5 is given by,

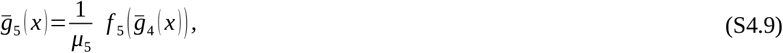

where we have assumed that gene product 4 reaches its stationary concentration 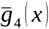 fast enough. We know from R5 in the main text, that *f* _5_ (*g*_4_) must preserve the monotony of 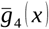 (i.e., it must increase where 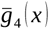 increases, and decrease where 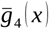 decreases). As a result, the resulting pattern of 5 replicates the resulting pattern of 4 in the sense that concentration peaks and valleys in 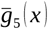 are located at the same positions than those of 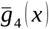. Nonetheless, by tuning the functional form of *f* _5_ (*g*_4_), as well as the value of model parameters like *μ*_5_, we can make the concentration peaks are valleys of 5 higher, deeper or steeper than those of 4.

All in all, we have proven that a simple diamond H network is enough to transform a spike initial pattern into a resulting pattern with two symmetric peaks of concentrations on both sides of the initial pattern. Moreover, as suggested by equations (S4.5) and (S4.9), model parameters and regulation function *f* (*g*) can be tuned to make such concentration peaks appear at any position out of the spike, and have any desired height and steepness.

### S5: Peak addition and subtraction in H gene regulatory networks

In this section, we are going to show how two patterned gene products in a hierarchic network can lead to more complex patterns in downstream gene products without requiring to additional extracellular signals. This is can be achieved by two simple mechanisms that include positive (for ‘addition’ of patterns) and negative (for ‘subtraction’ of patterns) intracellular regulations between the patterned gene products.

Let us consider two gene products *i* and *j* exhibiting heterogeneous resulting patterns 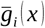 and 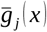 that directly regulate intracellular gene product *k*. If no other gene product in the network regulates *k*, its concentration dynamics for each *x* is given by,

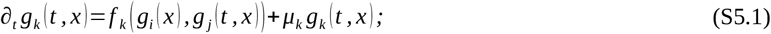

and then, assuming that *i* and *j* reach their respective stationary concentrations fast enough, we can take the time derivative in (S5.1) equal to zero and get that the stationary concentration of gene product *k* is given by,

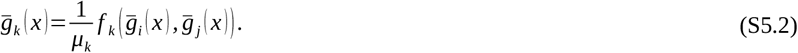

First, we consider the case of the peak addition mechanism, that is, we assume that both *i* and *j* positively regulate gene product *k* (see Figure 10 in the main text). In this case, according to requirement R5 in the main text, *f* _*k*_ (*g*_*i*_, *g* _*j*_) is component-wise monotonously increasing with respect to both *g*_*i*_ and *g* _*j*_. Hence, we know that the stationary concentration of gene product *k* will increase wherever the stationary concentrations of both *i* and *j* simultaneously increase; and it will decrease wherever those of *i* and *j*simultaneously decrease. As a result, if both 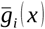 and 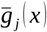 have a concentration peak at the same position 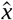, then 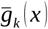 will also exhibit a peak at 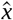.

Ambiguity appears in those cases for which the peaks of concentration of *i* and *j* do not coincide at the same spatial position. In this sense, let us assume that the stationary concentrations of gene products *i* and *j* exhibit a total amount of *N* and *M* concentration peaks along the domain, respectively; and let us focus in two consecutive such peaks, that is, let us consider a concentration peak in 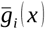 located at a given position 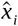 and a concentration peak in 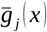 at a different position 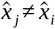 such that neither 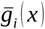 nor 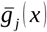 exhibit an intermediate concentration peak along the line going from 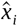 to 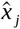. In this setting, if 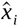 and 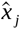 are far enough from each other and both concentration peaks are steep enough so that the influence of 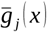 (resp., 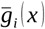) is negligible near 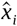 (resp., 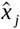); then, *f* _*k*_ (*g*_*i*_, *g* _*j*_) will behave as a function of only *g*_*i*_(resp., *g* _*j*_) in a vicinity of 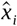 (resp., 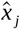) and thus, 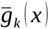 will exhibit one concentration peak at 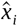 and another concentration peak at 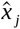. Moreover, if this condition holds for any pair of consecutive peaks of 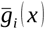 and 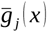, 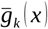 will exhibit a total amount of *N* + *M* concentration peaks along the domain (see Figure 10 in the main text). On the contrary, if two such consecutive peaks of *i* and *j* are too close to each other, the expected concentration peaks of *k* may merge into just one located along the line from 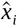 to 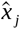 and then, the total number of concentration peaks that *k* will exhibit along the domain is strictly smaller than*N* + *M* .

On the other hand, let us consider the case of the peak subtraction mechanism, that is, let us assume that *i* activates *k* while *j* inhibits it (see Figure 10 in the main text). In this setting, according to requirement R5 in the main text, *f* _*k*_ (*g*_*i*_, *g* _*j*_) is monotonously increasing with respect to *g*_*i*_ and monotonously decreasing with respect to *g* _*j*_. As in the case of peak addition, let us assume that the stationary concentrations of gene products *i* and *j* display a total amount of *N* concentration peaks each. Also, let us assume that two such peaks of concentration coincide at the same position 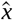. Then, if the activation of *i* on *k* is much stronger than the inhibition of *j* on *k*, 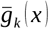 will portray a concentration peak at 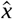. Conversely, if the inhibition of *j* on *k* is much stronger than the activation of *i* on *k*, no concentration peak of *k* will emerge at 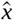. Thus, if the former situation takes place at all possible pairs of concentration peaks of *i* and *j*, 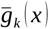 will display a total amount of *N* concentration peaks along the domain. However, if the latter situation takes place for all possible pairs of peaks, 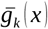 will retract back to a homogeneous concentration as all possible peaks of concentration will be eventually swept away by the inhibition of *j*.

However, as we have seen in section S3, under some specific conditions on function *f* and specific parameter values, the inhibition that *j* exerts on *k* can split the emerging concentration peak in two in such a way that, on a 1D domain, 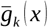 displays two symmetric concentration peaks on both sides of 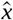 (this pair of symmetric peaks becomes a ring around 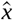 in 2D, and a hollow sphere in 3D). Moreover, this split would also be the case even if the concentration peaks in 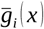 and 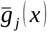 did not coincide at the same position, but stand in close positions (although in this case the height of the emerging peaks in 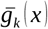 would be uneven). Anyhow, if this splitting of peaks occurs for all possible pairs of concentration peaks of *i* and *j*, 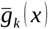 will exhibit a total amount of 2 *N* peaks along the 1D domain (in higher dimensions, each pair of peaks reduces to one single ring or sphere and hence, the total amount of structures in 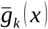 reduces to *N*).

### S6: Numerical simulations

In this supplementary section, we provide the necessary information to reproduce the numerical simulations leading to all resulting patterns presented in the various figures along the main text and supplementary information of this article. Hence, we first offer a detailed description of the Maini-Miura model (Miura and Maini, 2004; Marcon *et al*., 2016), which serves as the basis for the reaction-diffusion equations in all our simulations (i.e., equations (1) of the main text); and we list the specific parameter values applied in each case. All parameters presented in this supplementary section are expressed in arbitrary units (a.u.).

#### The Maini-Miura model

In all our simulations, the underlying reaction-diffusion equations correspond to the Maini-Miura model (Miura and Maini, 2004; Marcon *et al*., 2016), or slight modifications of it. The governing equations of this simple model of pattern formation are given by,

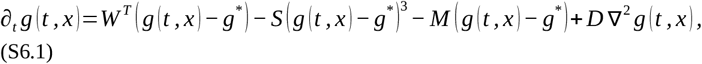

where 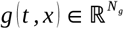 is the vector of gene product concentrations at time *t* ≥ 0 and position 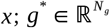 is a homogeneous steady state concentration; 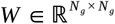 is the weight matrix of the corresponding gene regulatory network; 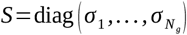 is the cubic saturation matrix; 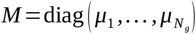 is the linear degradation matrix; and 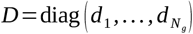 is the diffusion matrix. As in equation (1) of the main text, *g*^3^ denotes component wise exponentiation.

Function *f* (*g*) in equation (1) of the main text corresponds to the combination of terms *W*^*T*^ (*g* (*t, x*) − *g*^*^) −*S* (*g* (*t, x*) − *g*^*^)^3^ + *M g*^*^ in equation (S6.1). Given its polynomial structure, it is straightforward to check that it satisfies requirements R1 and R3 in the main text and, if *σ*_*i*_>0 for at least one gene product, also requirement R2. The Jacobian matrix of *f* (*g*) at *g*^*^ coincides with the transpose weight matrix (i.e., *J* _*f*_ (*g*^*^)=*W*^*T*^) and then, *f* (*g*) satisfies requirement R5 in the main text by continuity (at least in a neighborhood of *g*^*^). Regarding requirement R4 in the main text, concentration *g* (*t, x*) is the Maini-Miura model is always bounded from above because for large values of *g* (*t, x*) both degradation terms in (S6.1) become negative. Also, the entries in *g*^*^ can be tuned to avoid *g* (*t, x*)<0 for all time *t* ≥ 0 and all positions *x*.

Despite its many simplifications, the Maini-Miura model offers a practical framework for the study of pattern formation governed gene regulatory networks, as its formulation enables a direct manipulation of gene network topology by simply changing the entries of the weight matrix *W* . This adaptability makes the Maini-Miura model particularly suitable for the study of gene regulatory networks comprising a large number of gene products, unlike classical models such as the Schnackenberg (Schnackenberg, 1979; Murray, 2003) or Gierer-Meinhardt models (Gierer & Meinhardt, 1972; Meinhardt, 1982), which are constrained to a very specific two-node Turing gene network.

In our simulations, we build spike, spike-homogeneous and noise-homogeneous initial patterns using *g*^*^ as their basal concentrations. For spike- and noise-homogeneous initial patterns we consider 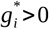 big enough so that *g*_*i*_ (*t, x*) never becomes negative. For spike initial patterns, we take *g*^*^ ≡ 0 and clamp possible negative concentration by introducing a smooth step-function in the first term of (S6.1), namely,

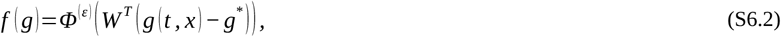

where each entry 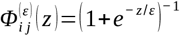 is a narrow sigmoid function, with *ε* >0 arbitrarily small. Note that the inclusion of the smooth step function *Φ*^(*ε*)^ in *f* (*g*) is consistent with requirements R1-R5 in the main text. Moreover, if we choose *ε* small enough, it acts as an effective clamping term in model equations that prevents the occurrence of negative concentrations during the simulations.

In a similar spirit, several figures in the main text and the supplementary information explicitly require different components of *f* (*g*) to adopt different functional forms (e.g., Figure 7 in the main text, evaluating conditions RH1 and RH2). In such simulations, only some specific components of *f* (*g*) are modified, leaving all remaining terms of equation (S6.1) unchanged. Each such modification is listed bellow, together with the corresponding set of model parameters that lead to the pattern transformation under consideration.

#### Parameter values for patterning simulations

All simulations in this article have been performed using a Forward-Euler algorithm (Grossmann and Ross, 2007) on a 1-dimensional interval [0, 50] split into a line array of 500 cells with a constant spatial step (i.e., distance between cell centers) of *h*=0.1, or a 2-dimensional square [0, 50] *×* [0, 50] split into a square ensemble of 250 *×* 250 cells with a constant spatial step of *h*=0.1. The time step *τ* of the algorithm is adjusted for every simulation in such away that the convergence condition of the Forward-Euler algorithm is satisfied for all gene products in the gene network (i.e., 2^*n*^ *τ d*_*i*_ <*h*^2^ for all *i* and *n*=1, 2). This results in a *D*-dependent time step of the form,

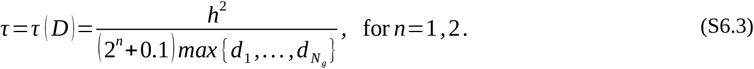

All simulations start with an initial pattern built around the equilibrium point of equation (S6.1), that is, 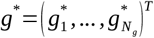. Spike (*g*^*^=0) and spike-homogeneous (*g*^*^ ≠ 0) initial conditions centered at *x*_0_ ∈ ℝ^*n*^ are given by rectangular functions of the form,

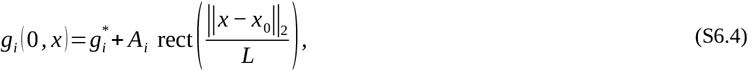

where *A*_*i*_>0 is the amplitude of the spike; *L*>0 is the width of the spike (equal for all gene products); || · ||_2_ denotes the euclidean norm; and rect (*y*)=1, if − 1 ≤ *y* ≤ 1, while rect (*y*)=0, elsewhere (Abramowitz and Stegun, 1964). On the other hand, noise-homogeneous initial patterns are given by 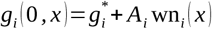, where *A*_*i*_>0 is the amplitude of the noise and wn (*x*) is a random value uniformly chosen in the interval [− 1, 1] for each different position *x* and each different gene product *i*∈ {1, …, *N*_*g*_ }.

*Figure 2 in the main text*

Figure 2 in the main text shows the pattern transformations that a 4-node hierarchical gene regulatory network produces on a spike (*g*^*^=0, *x*_0_=25, *A* =(0.05, 0, 0, 0)^*T*^, *L*=2), a combined spike-homogenenous (*g*^*^=(1, 0.5, 1, 0.6)^*T*^, *x*_0_=25, *A*=(0.05, 0, 0, 0)^*T*^, *L*=2) and a noise-homogeneous (*g*^*^=(1, 0.5, 1, 0.6)^*T*^, *A*_*i*_=0.05 for all *i*) initial pattern. Simulations for the spike initial conditions include the clamping introduced in (S6.2). The common choice of model parameters for all simulations corresponds to,

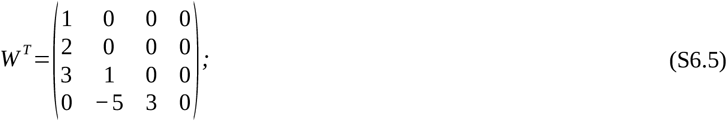

and

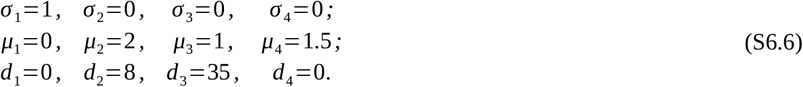

*Figure 7 in the main text*

Figure 7 in the main text shows the pattern transformations that two 5-node H ^0^ gene networks and one 4-node H gene network produce on a spike initial pattern (*g*^*^=0, *x*_0_=25, *A* =(0.05, 0, 0, 0)^*T*^, *L*=2). The two H^0^ networks include non-linear regulations in the first term of (S6.1), that is, one of the entries of function *f* . Also, the clamping introduced in (S6.2) is considered to avoid negative concentrations. The parameter choice for panel A is given by,

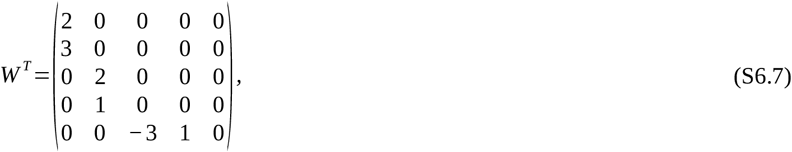

with 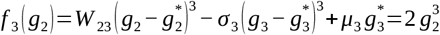; and

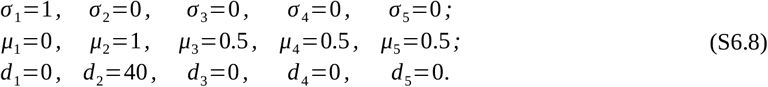

The parameter choice for panel B is given by,

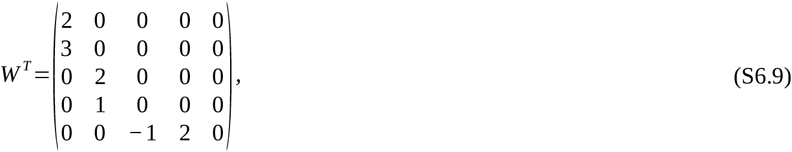

with 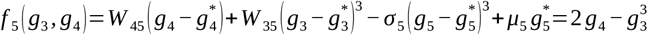;and

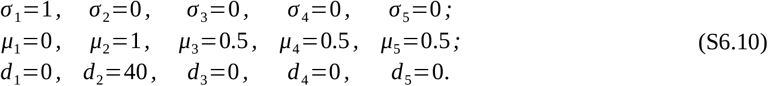

The parameter choice for panel C is given by,

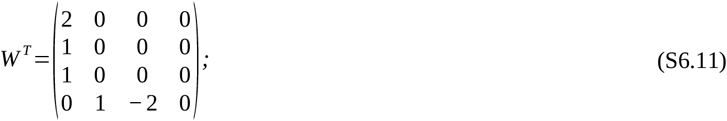

and

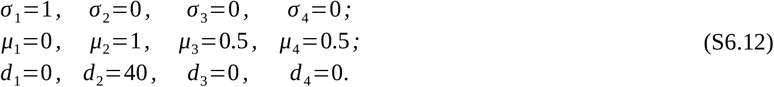

*Figures 8 and 9 in the main text*

Figures 8 and 9 in the main text show the 1D and 2D pattern transformations that a 4-node hierarchic gene network, a 3-node over-Turing gene network and a 3-node Turing gene network produce on a spike (*x*_0_=25 and *x*_0_=(25, 25)^*T*^, *A*=(0.05, 0, 0, 0)^*T*^, *L*_1D_=2 and *L*_2D_=3), a spike-homogeneous (*x*_0_=25 and *x*_0_=(25, 25)^*T*^, *A* =(0.05, 0, 0, 0)^*T*^, *L*_1D_=2 and *L*_2D_=3), and a noise-homogeneous (*A*_*i*_=0.05 for all *i*) initial pattern. Also, the clamping introduced in (S6.2) is considered in the case of spike initial patterns.

The choice of model parameters is different for each of the three types of gene regulatory networks, but identical regardless the initial pattern being considered. Networks of the same kind use the same set of model parameters in both figures. For the hierarchical network the parameter choice is given by,

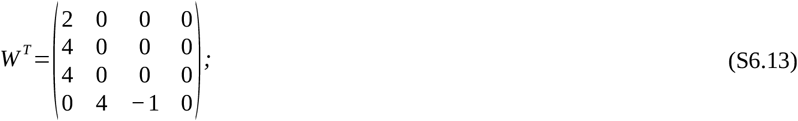

and

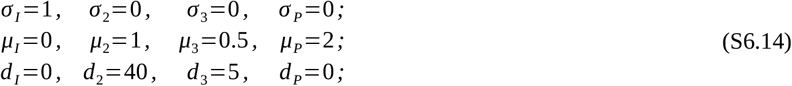

with *g*^*^=0 for the spike initial pattern, and *g*^*^=(1, 1, 2, 0.5)^*T*^ for the spike-homogeneous and noise-homogeneous initial patterns.

For the over-Turing network, the parameter choice is given by,

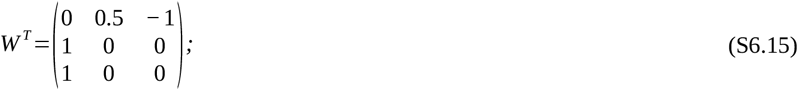

and

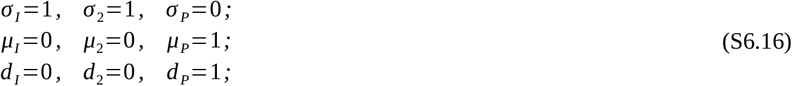

with *g*^*^=0 for the spike initial pattern, and *g*^*^=(1, 1, 1)^*T*^ for the spike-homogeneous and noise-homogeneous initial patterns.

Finally, for the Turing network, the choice of model parameters is given by,

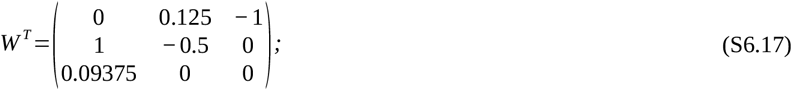

and

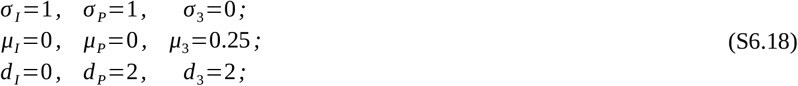

with *g*^*^=0 for the spike initial pattern, and *g*^*^=(1, 1, 4)^*T*^ for the spike-homogeneous and noise-homogeneous initial patterns.

Figure 10 in the main text shows the pattern transformations that two 6-node hierarchic gene regulatory networks produce on a spike initial pattern (*g*^*^=0, *x*_0_=25, *A* =(0.05, 0, 0, 0)^*T*^, *L*=2). The clamping introduced in (S6.2) is considered to prevent negative concentrations. The parameter choice for panel A is given by,

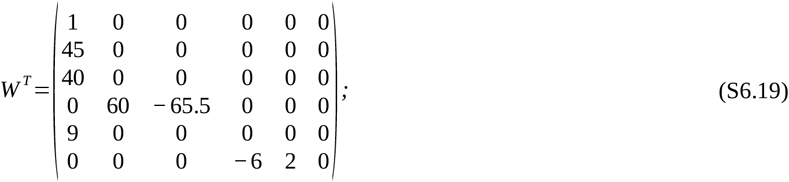

and

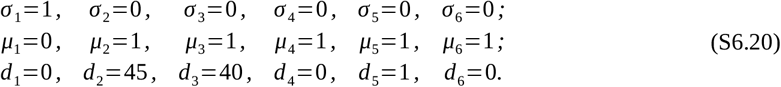

Similarly, the parameter choice for panel B is given by,

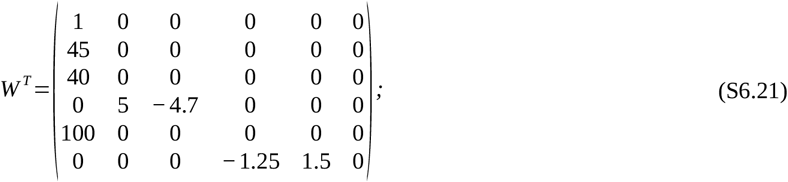

and

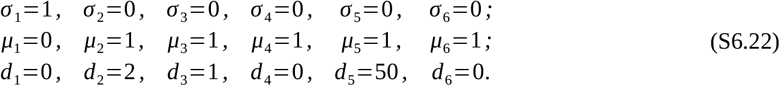

*Figure 11 in the main text*

Figure 11 in the main text shows the pattern transformations that a 5-node hierarchic gene network produces on a spike initial pattern (*g*^*^=0, *x*_0_ =25, *A* =(0.05, 0, 0, 0)^*T*^, *L*=2) for several sets of model parameters. The clamping introduced in (S6.2) is considered to prevent negative concentrations. The parameter choices for panel C correspond to,

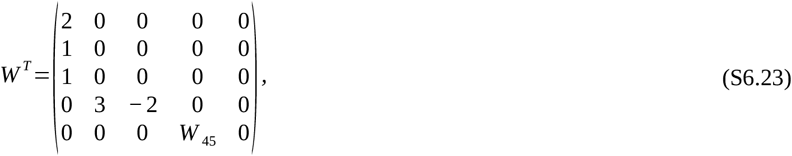

with *W* _45_ ∈ {0.5 \\(blue\\), 1 \\(green\\), 2 \\(red\\) }; and,

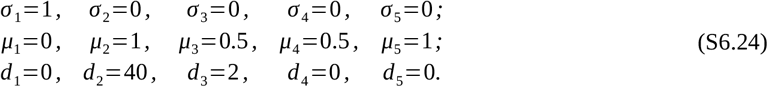

Similarly, the parameter choices for panel D are given by,

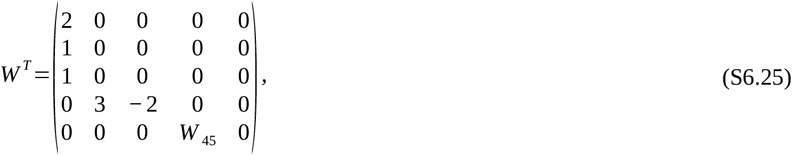

with *W* _45_ ∈ {0.94984 \\(blue\\), 1.79324 \\(green\\), 3.03495 \\(red\\) }; and,

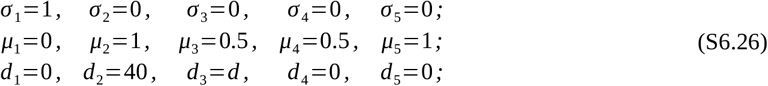

with *d* ∈ {0.5 \\(blue\\), 2 \\(green\\), 4 \\(red\\) }.

*Figure 12 in the main text*

Figure 12 in the main text shows the pattern transformations that the combinations of hierarchic, over-Turing and Turing gene regulatory subnetworks produce on a spike-homogeneous initial pattern (*x*_0_=25, *A*_*I*_ =0.05 and *A*_*i*_=0 for any other *i, L*=2). The choice of model parameters for (S6.1) is different for each of the combinations. Also, some of the simulations include non-linear regulations in the first term of system (S6.1), that is, one of the entries of function *f* .

The top-left panel shows the pattern transformation produced by an 8-node hierarchic + hierarchic gene network. The parameter choice in this case is given by,

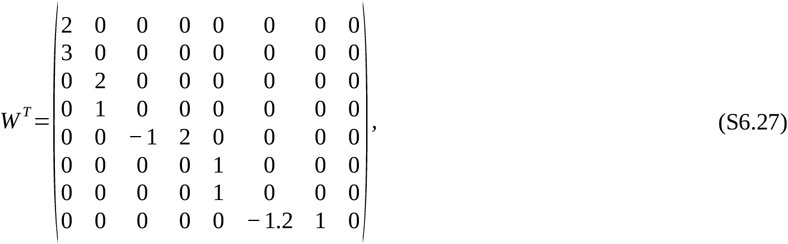

with 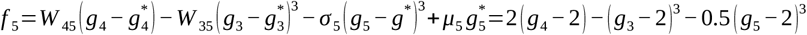 and 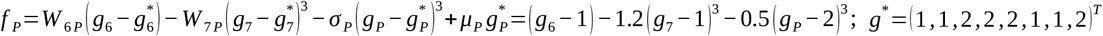 and,

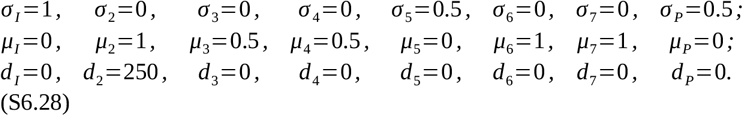

The top-middle panel shows the pattern transformation produced by an 8-node hierarchic + over-Turing gene network. The parameter choice in this case is given by,

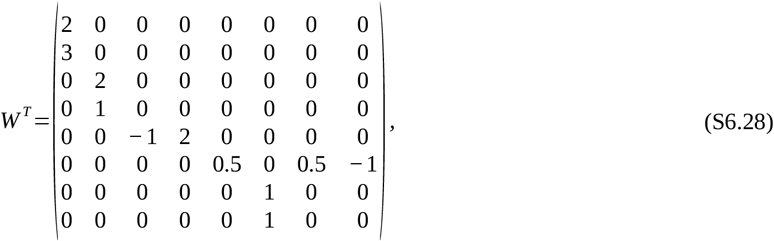

with 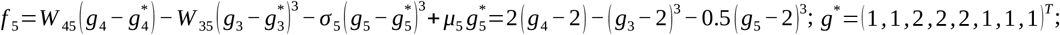; and,

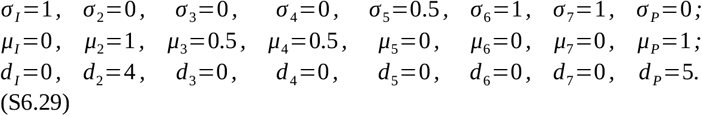

The top-right panel shows the pattern transformation produced by an 8-node hierarchic + Turing gene network. The parameter choice in this case is given by,

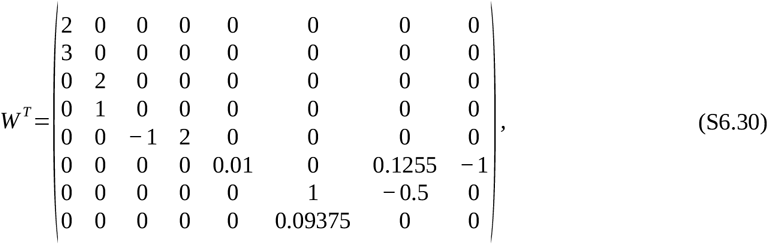

with 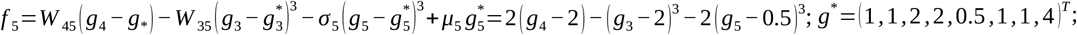; and,

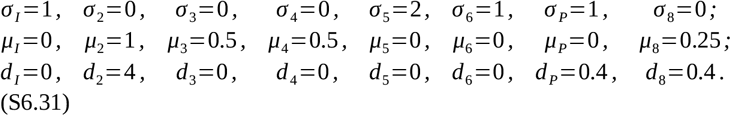

The middle-left panel shows the pattern transformation produced by a 6-node over-Turing + hierarchic gene network. The parameter choice in this case is given by,

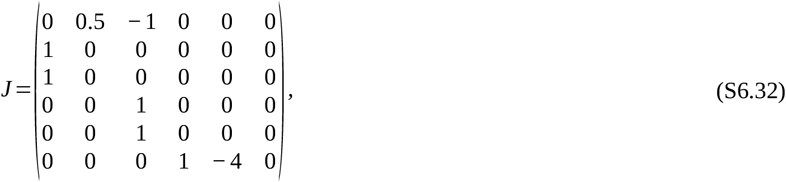

with 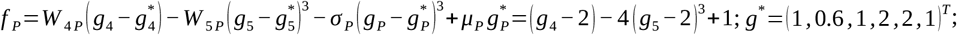; and,

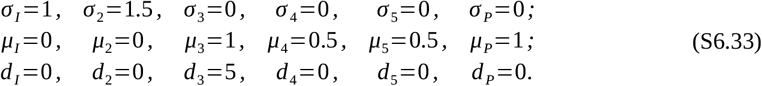

The central panel shows the pattern transformation produced by a 6-node over-Turing + over-Turing gene network. The parameter choice in this case is given by,

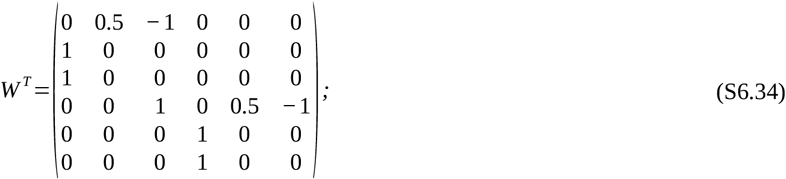

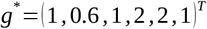; and,

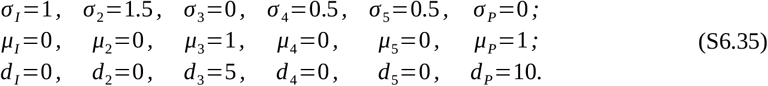

Panel B3 shows the pattern transformation produced by a 6-node over-Turing + Turing gene network. The parameter choice in this case is given by,

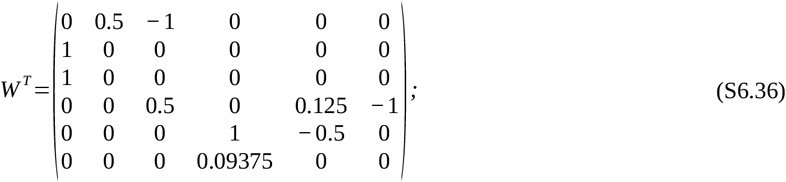

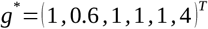; and,

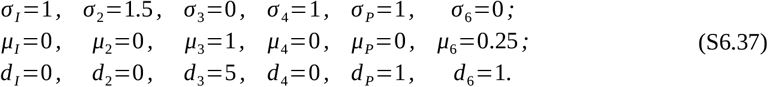

The bottom-left panel shows the pattern transformation produced by a 6-node Turing + hierarchic gene network. The parameter choice in this case is given by,

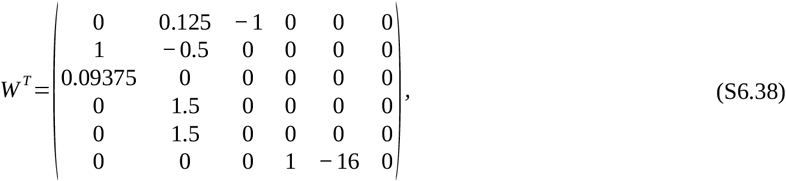

with 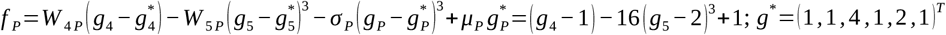; and,

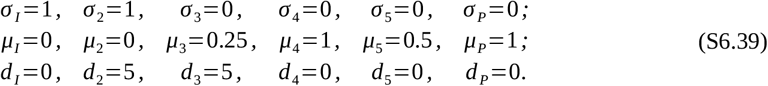

The bottom-middle panel shows the pattern transformation produced by a 6-node Turing + over-Turing gene network. The parameter choice in this case is given by,

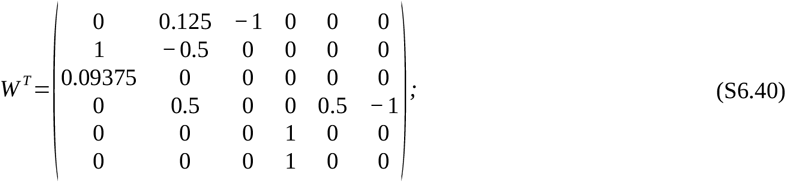

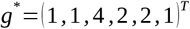; and,

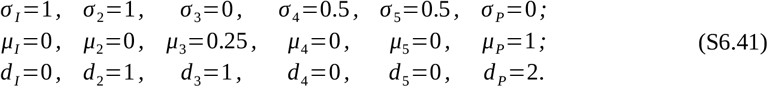

Finally, the bottom-right panel shows the pattern transformation produced by a 6-node Turing + Turing gene network. The parameter choice in this case is given by,

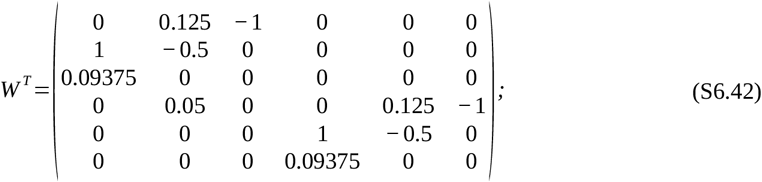

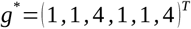; and,

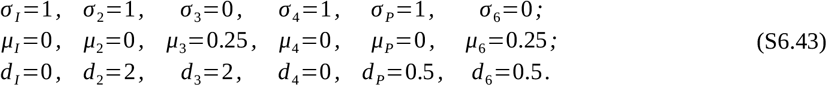

*Figure S2 in the supporting information*

Figure S2 in the supporting information shows the pattern transformations that an 8-node hierarchic gene network produces on a spike initial pattern (*g*^*^=0, *x*_0_=25, *A* =(0.05, 0, 0, 0)^*T*^, *L*=2) for several sets of model parameters. The clamping introduced in (S6.2) is considered to prevent negative concentrations. The parameter choices for panel C correspond to,

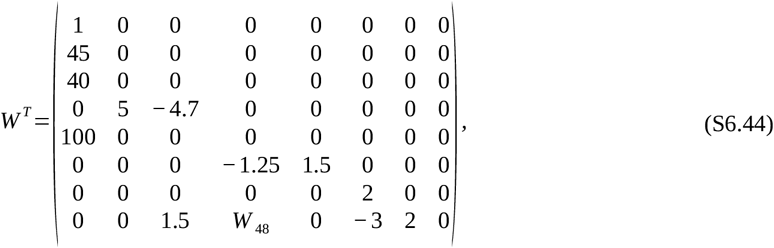

with *W* _48_ ∈ {− 1.8 \\(blue\\), − 0.3 \\(red\\) }; and,

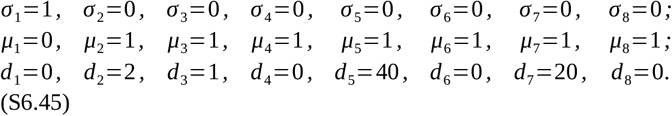

Similarly, the parameter choices for panel D are given by,

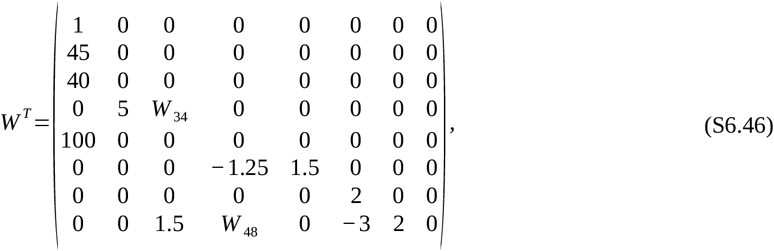

with *W* _34_ ∈ {− 4.7 \\(blue\\), − 4.9 \\(green\\) } and *W* _48_ ∈ {− 1.8 \\(blue\\), − 2.7 \\(green\\) }; and,

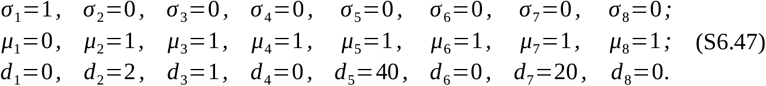

*Figure S3 in the supporting information*

Figure S3 in the supporting information shows the pattern transformations that a 2-node over-Truing network produces on a noise-homogeneous initial pattern (*g*^*^=(1.25, 0.6)^*T*^, *A*_*i*_=0.05 for all *i*) and two spike-homogeneous initial patterns (*g*^*^=(1.25, 0.6)^*T*^, *x*_0_=25 (left), *x*_0_=0 (right), *A*=(0.05, 0, 0, 0)^*T*^, *L*=2). The parameter choices for all panels in the same and it is given by,

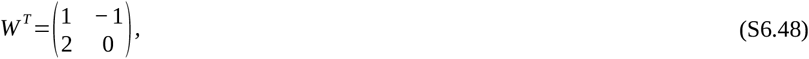

and,

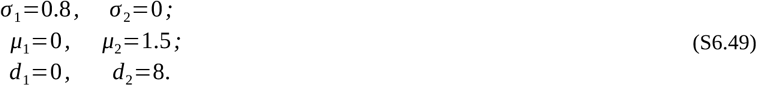

*Figure S4 in the supporting information*

Figure S4 in the supporting information show the pattern transformations that a 3-node over-Turing gene network and a 3-node Turing gene network produce on a spike-homogeneous initial pattern (*x*_0_=25 (left), *x*_0_=0 (right), *A* =(0.05, 0, 0)^*T*^, *L*=2). In panel D, the basal concentration is *g*^*^=(1, 0.6, 1)^*T*^, and the choice of model parameters is given by,

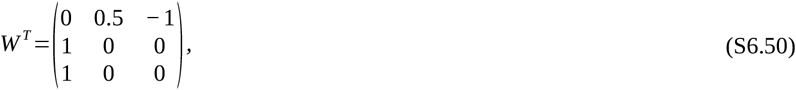

and,

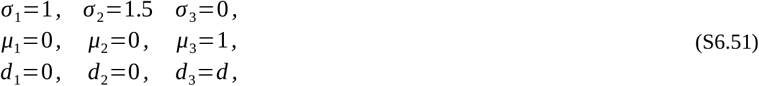

with *d* ∈ {0.5 \\(blue\\), 1 \\(green\\), 5 \\(red\\) }.

In panel E, the basal concentration is *g*^*^=(1, 1, 4)^*T*^, and the choice of model parameters is given by,

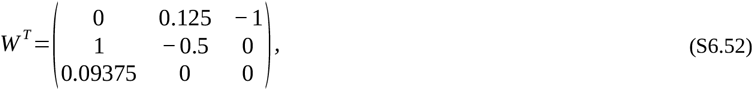

and,

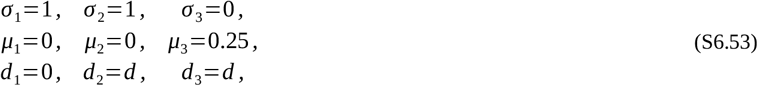

with *d* ∈ {0.5 \\(blue\\), 1 \\(green\\), 2 \\(red\\) }.

*Figure S5 in the supporting information*

Figure S5 in the supporting information shows the pattern transformations that a 2-node L gene network, a 4-node l^+^L^-^ gene network and a 4-node L^+^L^-^ gene network produce on a spike (*x*_0_=25, *A*=(0.05, 0, 0, 0)^*T*^, *L*=2), a spike-homogeneous (*x*_0_=25, *A* =(0.05, 0, 0, 0)^*T*^, *L*=2), and a noise-homogeneous (*A*_*i*_=0.05 for all *i*) initial pattern. The clamping introduced in (S6.2) is considered in the case of spike initial patterns.

The choice of model parameters is different for each of the three types of gene regulatory networks, but identical regardless the initial pattern being considered. For the L^+^ network the parameter choice is given by,

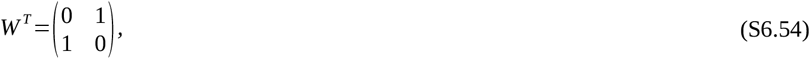

and,

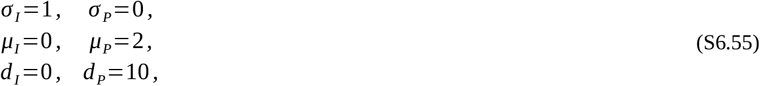

with *g*^*^=0 for the spike initial pattern, and *g*^*^=(1, 0.5)^*T*^ for the spike-homogeneous and noise-homogeneous initial patterns.

For the l^+^L^-^ network, the parameter choice is given by,

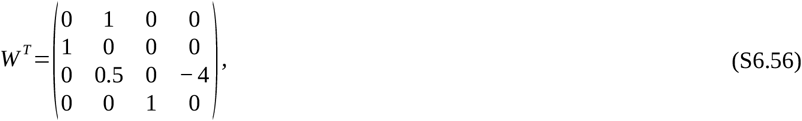

and,

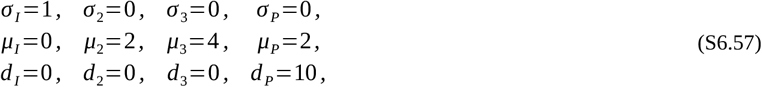

with *g*^*^=0 for the spike initial pattern, and *g*^*^=(1, 0.5, 0.25, 0.5)^*T*^ for the spike-homogeneous and noise-homogeneous initial patterns.

For the L^+^L^-^ network, the parameter choice is give by,

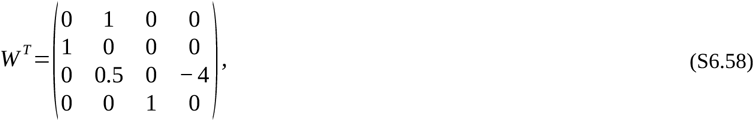

and,

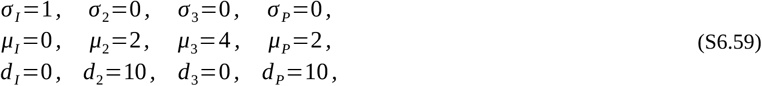

with *g*^*^=0 for the spike initial pattern, and *g*^*^=(1, 0.5, 0.25, 0.5)^*T*^ for the spike-homogeneous and noise-homogeneous initial patterns.

*Figure S6 in the supporting information*

Figure S6 in the supporting information shows the pattern transformations that a 3-node L gene network, a 5-node L^-^l^+^ gene network and a 5-node L^-^L+ gene network produce on a spike (*x*_0_=25, *A*=(0.05, 0, 0, 0)^*T*^, *L*=2), a spike-homogeneous (*x*_0_=25, *A* =(0.05, 0, 0, 0)^*T*^, *L*=2), and a noise-homogeneous (*A*_*i*_=0.05 for all *i*) initial pattern. The clamping in (S6.2) is considered in the case of spike initial patterns.

The choice of model parameters is different for each of the three types of gene regulatory networks, but identical regardless the initial pattern being considered. For the L^-^ network the parameter choice is given by,

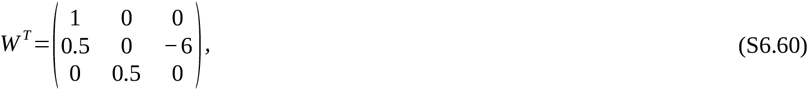

and,

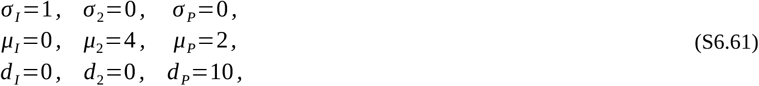

with *g*^*^=0 for the spike initial pattern, and *g*^*^=(1, 0.25, 0.5)^*T*^ for the spike-homogeneous and noise-homogeneous initial patterns.

For the l^+^L^-^ network, the parameter choice is given by,

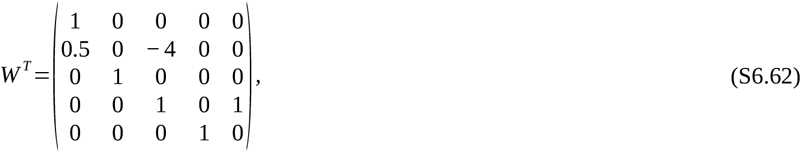

and,

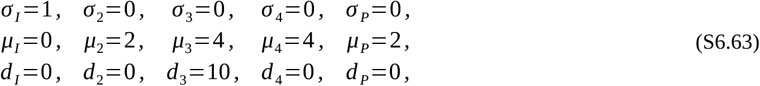

with *g*^*^=0 for the spike initial pattern, and *g*^*^=(1, 0.5, 0.25, 0.25, 0.5)^*T*^ for the spike-homogeneous and noise-homogeneous initial patterns.

For the L^+^L^-^ network, the parameter choice is give by,

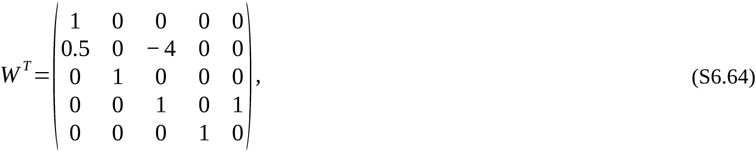

and,

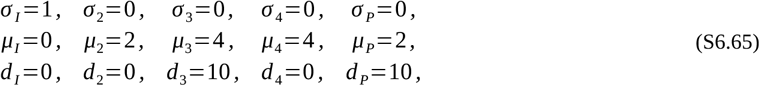

with *g*^*^=0 for the spike initial pattern, and *g*^*^=(1, 0.5, 0.25, 0.25, 0.5)^*T*^ for the spike-homogeneous and noise-homogeneous initial patterns.

---

**Figure S1. If function *f* did not satisfy R5, then a two gene network could lead to any resulting pattern in which concentrations are finite. (A)** Gene network topology. An extracellular signal activating an intracellular gene product. 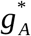 stands for the concentration of the extracelllular signal in the steady state and 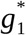 for that of the intracellular gene product. **(B)** The extracellular signal is secreted from the boundary of an array of cells (x-axis), leading to a gradient in the concentration of the concentration of the signal. Labels c1 to c6 are shown to facilitate the interpretation of the figures but we assume *f* can be understood as continuous. **(C)** Different example functions *f* are plotted. In the x-axis we represent the different concentrations of the signal and in the y-axis the concentration of the intracellular gene product arising from each concentration of the signal. Notice that none of the example functions *f* satisfy R5, i.e. the function is not monotonously growing with the concentration of the signal **(D)** The resulting patterns arising from each *f* . By choosing an adequate *f*, any resulting pattern can be achieved.

**Figure S2. Example H gene network where by changing the parameters it is possible to vary the height of the different concentration peaks. (A)** Gene network topology. **(B)** Spike initial pattern. **(C-D)** The blue, red and green plots show the concentration of gene product 8 for different combinations of parameter values. As we can see, the height of the concentration peaks of 8 around the initial spike can be independently tuned. Simulations were run using a Forward-Euler algorithm on the Maini-Miura model (see section S6 for parameter values).

**Figure S3. Pattern formation in over-Turing gene networks. (A)** Noise-homogeneous initial patterns are transformed into amplified noise patterns. Even with the same choices of model parameters, the amplified noise resulting patterns are different every time. **(B)** Spike-homogeneous initial patterns lead to periodical patterns. If the choice of model parameters is the same, the final pattern is the same for each transformation. **(C)** The peaks of concentration formed from a spike-homogeneous initial pattern are produced one by one. The whole process resembles a traveling wave that moves away from the spike and, at the same time, freezes whenever a new peak is formed. Network colors and shapes as in Figure 2. *P*stands for the gene product plotted as resulting pattern while *I* stands for the gene product in the initial pattern. Simulations as in Figure S2 (see section S6 for parameter values).

**Figure S4. Variational properties of emergent networks. (A-B)** An over-Turing (left) and Turing (right) gene networks produce a very similar periodical pattern from a spike-homogeneous initial pattern. A change in any model parameter (such as the diffusivity of an extracellular signal) alters the height and position of all peaks at the same time. **(C)** Spike initial pattern. **(D-E)**. Network colors and shapes as in Figure S1. *P* stands for the gene product plotted as resulting pattern while *I* stands for the gene product in the initial pattern. Simulations as in Figure S3 (see section S6 for parameter values).

**Figure S5. L^+^, l^+^L^-^ and L^+^L^-^ networks do not produce non-trivial pattern transformations.** The first column depicts simple examples of each type of these gene network topologies. The upper row shows the three initial patterns. Intermediate panels show each type of possible resulting pattern arising from each combination of initial pattern and gene network topology. Note that all pattern transformations are trivial. Network colors and shapes as in Figure S1. The y-axis in all plots represents gene product concentrations. *P* stands for the gene product plotted as resulting pattern while *I* stands for the gene product in the initial pattern. Simulations as in Figure S3 (see section S6 for parameter values).

**Figure S6. L^-^, L^-^l^+^ and L^-^L^+^ networks do not produce non-trivial pattern transformations.** The first column depicts simple examples of each type of these gene network topologies. The upper row shows the three initial patterns. Intermediate panels show each type of possible resulting pattern arising from each combination of initial pattern and gene network topology. Note that all pattern transformations are trivial. Network colors and shapes as in Figure S1. The y-axis in all plots represents gene product concentrations. *P* stands for the gene product plotted as resulting pattern while*I* stands for the gene product in the initial pattern. Simulations as in Figure S3 (see section S6 for parameter values).

