## Supplementary material for "The zoo of the gene networks capable of pattern formation by extracellular signaling": Supplemenary material

Isaac Salazar-Ciudad

**This PDF file includes:**

Supporting text
Figures S to S
SI References

### Suporting text

#### S1: Formal definition of pattern transformation

Let us consider an arbitrary gene regulatory network, and let $g_i(t, \boldsymbol{x})$ denote the concentration of gene product $i$ at time $t \geq 0$ and position $\boldsymbol{x}$ in an $n$-dimensional spatial domain, with $n = 1, 2, 3$. At the beginning of the patterning process, gene product $i$ exhibits an initial pattern given by $g_i^{(ini)}(\boldsymbol{x}) = g_i(0, \boldsymbol{x})$. At the end of the patterning process, given requirement R2 in the main text, we know that the concentration of gene product $i$ converges to a stationary resulting pattern that we denote by $g_i^{(res)}(\boldsymbol{x}) = \lim_{t \to \infty} g_i(t, \boldsymbol{x})$. For the purpose of this article, we ignore non-stationary solutions to the model equations (e.g., time-periodic molecular clocks within cells).

In this framework, we say the gene product $i$ undergoes a pattern transformation if its concentration changes during the patterning process at at least one position, that is,

$$g_i^{(ini)}(\boldsymbol{x}) \neq g_i^{(res)}(\boldsymbol{x}), \text{ for some } \boldsymbol{x} \text{ in the spatial domain}. \quad \text{(S1.1)}$$

Moreover, we say that gene product $i$ undergoes a heterogeneous pattern transformation if the resulting pattern at the end of the patterning process is heterogeneous, that is,

$$g_i^{(res)}(\boldsymbol{x}) \neq g_i^{(res)}(\boldsymbol{y}) \text{ for some } \boldsymbol{x} \neq \boldsymbol{y} \text{ in the domain}. \quad \text{(S1.2)}$$

We say that gene product $i$ undergoes a rearranging pattern transformation if the position of some critical point of its concentration changes along the patterning process (e.g., the emergence of a new maximum of concentration). If $g^{(ini)_i}(\boldsymbol{x})$ and $g^{(res)_i}(\boldsymbol{x})$ are both sufficiently smooth, this means that either,

$$\nabla g_i^{(ini)}(\boldsymbol{x}) \neq 0 \text{, while } \nabla g_i^{(res)}(\boldsymbol{x}) = 0 \text{, for some } \boldsymbol{x} \text{ in the domain,} \tag{S1.3}$$

indicating the emergence of a new critical point in the resulting pattern $g^{(res)_i}(\boldsymbol{x})$; or,

$$\nabla g_i^{(ini)}(\boldsymbol{x}) = 0 \text{, while } \nabla g_i^{(res)}(\boldsymbol{x}) \neq 0 \text{, for some } \boldsymbol{x} \text{ in the domain,} \tag{S1.4}$$

indicating the disappearance of a critical point from the initial pattern $g^{(ini)_i}(\boldsymbol{x})$. However, none of the three basic initial patterns that we take into consideration in this article is even continuous in space. In this sense, we consider that $\nabla g_i^{(ini)}(\boldsymbol{x}) = 0$ for all $\boldsymbol{x}$ in the domain, if $g_i^{(ini)}(\boldsymbol{x})$ is a noise-homogeneous initial pattern; and we consider $\nabla g_i^{(ini)}(\boldsymbol{x}) \neq 0$ for all $\boldsymbol{x} \neq \boldsymbol{x}_c$ and $\nabla g_i^{(ini)}(\boldsymbol{x}_c) = 0$, if $g_i^{(ini)}(\boldsymbol{x})$ is a spike or spike-homogeneous initial pattern centered at some position $\boldsymbol{x}_c$. Formally, this two *ad hoc* assumptions correspond to the gradient of the convolution of $g_i^{(ini)}(\boldsymbol{x})$ with the Gaussian kernel of diffusion for a brief time $[0, \tau)$, with $\tau > 0$ arbitrarily small.

Finally, we say that gene product $i$ undergoes a non-replicating pattern transformation if the spatial arrangement of critical points in its resulting pattern does not coincide with the spatial arrangement of critical points in the initial pattern of any other gene product (i.e., any new piece of spatial information in the resulting pattern of $i$ must be created during the patterning process and not be replicated from any initial pattern). This means that, for any gene product $j \neq i$ in the gene network, either,

$$\nabla g_i^{(res)}(\boldsymbol{x}_j) = 0 \text{, while } \nabla g_j^{(ini)}(\boldsymbol{x}_j) \neq 0 \text{, for some } \boldsymbol{x}_j \text{ in the domain,} \tag{S1.5}$$

indicating that $g_i^{(res)}(\boldsymbol{x})$ exhibits a critical point at a position where $g_j^{(res)}(\boldsymbol{x})$ does not; or,

$$\nabla g_i^{(res)}(\boldsymbol{x}_j) \neq 0 \text{, while } \nabla g_j^{(ini)}(\boldsymbol{x}_j) = 0 \text{, for some } \boldsymbol{x}_j \text{ in the domain,} \tag{S1.6}$$

indicating that $g_j^{(res)}(\boldsymbol{x})$ exhibits a critical point at a position where $g_i^{(res)}(\boldsymbol{x})$ does not. Of course, the same considerations about the non-smoothness of initial conditions in conditions (S1.3) and (S1.4) must also be taken into consideration for (S1.5) and (S1.6).

All in all, for the purposes of this article, we say that gene product $i$ undergoes a non-trivial pattern transformation if it undergoes a heterogeneous, rearranging and non-replicating pattern transformation, that is, if $g_i^{(ini)}(\boldsymbol{x})$ and $g_i^{(res)}(\boldsymbol{x})$ satisfy conditions (S1.1) to (S1.6). Condition (S1.2), together with the fact that the resulting pattern must be stationary, correspond to requirement P1 in the main text. Conditions (S1.3) to (S1.6) correspond to requirement P2 in the main text.

#### S2: Positive regulatory loops and RD-instability

For a gene network to be RD-unstable of any kind it needs to have at least one positive regulatory loop: if all positive loops are all extracellular, then the gene network is RD-unstable of the first kind; if positive loops are all intracellular, then the gene network is RD-unstable of the second kind; and if the network has positive extracellular and intracellular loops, the type of RD-instability will depend on the particular choice of model parameters. In the main text, we provided proves for these statements in the case where the gene network displays only one regulatory loop, and this loop is positive. Here, we complete such results with a general proof in the case any number of positive and negative loops (i.e., for any general gene regulatory network).

Let $\boldsymbol{R}=\boldsymbol{R}(0)\in \mathbb{R}^{N_g\times N_g}$ denote the reaction-diffusion matrix of a given gene network in the absence of diffusion. Given a set $\{i_1,\ldots,i_{N_l}\}$ of $N_l\leq N_g$ gene products, the principal submatrix of $\boldsymbol{R}$ associated to $\{i_1,\ldots,i_{N_l}\}$ corresponds to the $N_l\times N_l$-matrix $\boldsymbol{R}_{\{i_1,\ldots,i_{N_l}\}}\in \mathbb{R}^{N_l\times N_l}$ resulting from considering only those entries in $\boldsymbol{R}$ whose row and column indices are given by $\{i_1,\ldots,i_{N_l}\}$ (i.e., considering only regulations between the gene products in $\{i_1,\ldots,i_{N_l}\}$). In this sense, the principal minor $\Delta_{\{i_1,\ldots,i_{N_l}\}}(\boldsymbol{R})\in \mathbb{R}$ associated to $\{i_1,\ldots,i_{N_l}\}$ is the determinant of the corresponding principal submatrix of $\boldsymbol{R}$, that is,

$$\Delta_{\{i_1,\ldots,i_{N_l}\}}(\boldsymbol{R})=\det\left(\boldsymbol{R}_{\{i_1,\ldots,i_{N_l}\}}\right). \qquad \text{(S2.1)}$$

If no gene products in $\{i_1,\ldots,i_{N_l}\}$ form a regulatory loop, the corresponding principal minor in (S2.1) reduces to the product of the diagonal entries in $\boldsymbol{R}_{\{i_1,\ldots,i_{N_g}\}}$, namely,

$$\Delta_{\{i_1,\ldots,i_{N_l}\}}(\boldsymbol{R})=(-1)^{N_l}\prod_{j=1}^{N_l} m_{i_j}\mu_{i_j}\left(g_{i_j}^{*}\right)^{m_{i_j}-1}, \qquad \text{(S2.2)}$$

where $\mu_{i_j}$ is the degradation coefficient of gene product $i_j\in\{i_1,\ldots,i_{N_l}\}$; $m_{i_j}$ is the order of such degradation; and $g_{i_j}^{*}$ the corresponding steady-state concentration around which $R$ was constructed (see equation (14) in the main text). If $\{i_1,\ldots,i_{N_l}\}$ form one, and only one, regulatory loop, the corresponding principal minor is given by,

$$\Delta_{\{i_1,\ldots,i_{N_l}\}}(\boldsymbol{R})=(-1)^{N_l-1}\left(J_{i_1 i_{N_l}}\prod_{j=2}^{N_l} J_{i_j i_{j-1}}-\prod_{j=1}^{N_l} m_{i_j}\mu_{i_j}\left(g_{i_j}^{¿}\right)^{m_{i_j}-1}\right), \qquad \text{(S2.3)}$$

where $J_{i_j i_k}$ represents the regulation that gene product $i_j\in\{i_1,\ldots,i_{N_l}\}$ exerts of gene product $i_k\in\{i_1,\ldots,i_{N_l}\}$ (see equation (19) in the main text). If $\{i_1,\ldots,i_{N_l}\}$ form more than one regulatory loop, each with a different length, more convoluted expressions than (S2.3) can be found using Leibniz formula (Strang, 2016).

Thus, in the case of regulatory loops, principal minors measure the net effect that the corresponding regulatory loop has on reaction-diffusion. For example, if we consider a direct self-regulatory loop (i.e., $N_l=1$), we get,

$$\Delta_{\{i_1\}}(\boldsymbol{R}) = J_{i_1 i_1} - m_{i_1} \mu_{i_1} \left(g_{i_1}^{*}\right)^{m_{i_1}-1}, \qquad \text{(S2.4)}$$

with $J_{i_1 i_1} > 0$ for a positive loop, and $J_{i_1 i_1} < 0$ for a negative one. Then, for any choice of model parameters for which $m_{i_1} \mu_{i_1} \left(g_{i_1}^{*}\right)^{m_{i_1}-1} > J_{i_1 i_1}$, we have that the degradation overpowers any positive effect of the self-regulatory loop and then, any concentration perturbation in $i_1$ would decay back to the steady-state concentration $g_{i_1}^{*}$ (i.e., perturbations are stable). However, if for some choice of model parameters $J_{i_1 i_1} > m_{i_1} \mu_{i_1} \left(g_{i_1}^{*}\right)^{m_{i_1}-1}$, then concentration perturbations in $i_1$ wouls grow in time (i.e., perturbations are unstable).

With this in mind, we say that a principal minor $\Delta_{\{i_1,\ldots,i_{N_l}\}}(\boldsymbol{R})$ of the reaction-diffusion matrix $\boldsymbol{R}$ is a destabilizing minor if there exists some choice of model parameters such that,

$$(-1)^{N_l} \Delta_{\{i_1,\ldots,i_{N_l}\}}(\boldsymbol{R}) < 0. \qquad \text{(S2.5)}$$

The following result shows that the principal minor associated to any set of gene products $\{i_1,\ldots,i_{N_l}\}$ that forms a positive regulatory loop is indeed a destabilizing minor:

*Lemma S2.1. Let us consider a gene regulatory network with $N_g \geq 1$ gene products, and let $\{i_1,\ldots,i_{N_l}\}$, with $N_l \leq N_g$, form a positive regulatory loop; then, the principal minor of $\boldsymbol{R}$ associated to $\{i_1,\ldots,i_{N_l}\}$ is a destabilizing minor of $\boldsymbol{R}$.*

**Proof (of Lemma S2.1).-** Without loss of generality, we assume that the $N_l$ gene products in the positive loop are labeled $\{1,\ldots,N\}$ (otherwise, we can simply reorder the labels in the gene network without altering its network topology). Moreover, for the sake of simplicity, we denote by $\hat{\mu}_i = m_i \mu_i \left(g_i^{¿}\right)^{m_i-1}$ the whole degradation term corresponding to gene product $i \in \{1,\ldots,N_l\}$.

The principal submatrix of $\boldsymbol{R}$ associated to the positive loop $\{1,\ldots,N_l\}$ is given by,

$$\boldsymbol{R}_{\{1,\ldots,N_l\}} = \begin{pmatrix} J_{11}-\hat{\mu}_1 & J_{12} & J_{13} & \cdots & J_{1(N_l-1)} & \boldsymbol{J_{1N_l}} \\ \boldsymbol{J_{21}} & J_{22}-\hat{\mu}_2 & J_{23} & \cdots & J_{2(N_l-1)} & J_{2N_l} \\ J_{31} & \boldsymbol{J_{32}} & J_{33}-\hat{\mu}_3 & \cdots & J_{3(N_l-1)} & J_{3N_l} \\ \vdots & \vdots & \vdots & \ddots & \vdots & \vdots \\ J_{(N_l-1)1} & J_{(N_l-1)2} & J_{(N_l-1)3} & \cdots & J_{(N_l-1)(N_l-1)}-\hat{\mu}_{(N_l-1)} & J_{(N_l-1)N_l} \\ J_{N_l1} & J_{N_l2} & J_{N_l3} & \cdots & \boldsymbol{J_{(N_l-1)N_l}} & J_{N_¿}-\hat{\mu}_{N_l} \end{pmatrix}, \qquad \text{(S2.6)}$$

where the bold entries denote the regulations of the positive regulatory loop formed by $\{1,\ldots,N_l\}$ (i.e., the regulation of 1 on 2, of 2 on 3, and so forth). By definition of positive loop, this entries represent a set of non-zero entries of $\boldsymbol{R}_{\{1,\ldots,N_l\}}$ whose product is strictly positive (see Figure 4 in the main text). The remaining entries in (S2.6) correspond to other possible regulations between gene products $\{1,\ldots,N_l\}$ that do not belong to the main loop and hence, they can all be equal to zero (e.g., equation (16) in the main text), or not.

Using Leibniz formula (Strang, 2016) for the determinant of (S2.6), we can write the principal minor of $\boldsymbol{R}$ associated $\{1,\ldots,N_l\}$ as a sum of the form,

$$\Delta_{\{1,\ldots,N_l\}}(\boldsymbol{R})=\sum_{\sigma\in\mathcal{S}_{N_l}}(-1)^{\kappa(\sigma)}R_{\sigma(1)1}R_{\sigma(2)2}\cdots R_{\sigma(N_l)N_l}, \tag{S2.7}$$

where $R_{ij}\in\mathbb{R}$ are the corresponding entries of $\boldsymbol{R}_{\{1,\ldots,N_l\}}$ (i.e., $R_{ij}=J_{ii}-\hat{\mu}_i$, if $j=i$; and $R_{ij}=J_{ij}$, if $j\neq i$); $\mathcal{S}_{N_l}$ is the symmetric group of all possible permutations $\sigma$ of the $N_l$-element set $\{1,\ldots,N_l\}$ (Hungerford, 1974); and, $\kappa(\sigma)$ is the number of inversions made by made by $\sigma$, that is, the number of ordered pairs $(i,j)$, with $i<j$, for which $\sigma(i)>\sigma(j)$.

In this sense, the product of all bold entries in (S2.6) corresponds to the term in (S2.7) defined by the cyclic permutation,

$$\sigma_c=\begin{pmatrix}1 & 2 & 3 & \cdots & N_l-1 & N_l\\ N_l & 1 & 2 & \cdots & N_l-2 & N_l-1\end{pmatrix}, \tag{S2.8}$$

where each element in $\{1,\ldots,N_l\}$ (i.e., first row of (S2.8)) permutes with the element strictly to its left (i.e., second row of (S2.8)), except for $1$, that permutes with $N_l$. As we can see, $\sigma_c$ has $N_l-1$ inversions given by pairs $(1,i)$ for all $i\in\{2,\ldots,N_l\}$ (indeed, $1<i$ while $\sigma_c(1)=N_l>\sigma_c(i)$). Consequently, $\kappa(\sigma_c)=N_l-1$, and then, the principal minor in (S2.7) can be written as,

$$\Delta_{\{1,\ldots,N_l\}}(\boldsymbol{R})=(-1)^{N_l-1}J_{12}J_{23}\cdots J_{N_l1}+\sum_{\sigma\in\mathcal{S}_{N_l}\setminus\{\sigma_c\}}(-1)^{\kappa(\sigma)}R_{\sigma(1)1}R_{\sigma(2)2}\cdots R_{\sigma(N_l)N_l}. \tag{S2.9}$$

Then, multiplying both sides of (S2.9) by a factor of $(-1)^{N_l}$, we get,

$$(-1)^{N_l}\Delta_{\{1,\ldots,N_l\}}(\boldsymbol{R})=-J_{12}J_{23}\cdots J_{N_l1}+\sum_{\sigma\in\mathcal{S}_{N_l}\setminus\{\sigma_c\}}(-1)^{N_l+\kappa(\sigma)}R_{\sigma(1)1}R_{\sigma(2)2}\cdots R_{\sigma(N_l)N_l}. \tag{S2.10}$$

As we mentioned before, the product $J_{12}J_{23}\cdots J_{N_l1}$ is strictly positive by definition and hence, the first term in (S2.10) is strictly negative. Consequently, regardless of the sign of the second term, there always exists some choice of model parameters for which (S2.10) becomes negative, namely,

$$J_{12}J_{23}\cdots J_{N_l1}>\sum_{\sigma\in\mathcal{S}_{N_l}\setminus\{\sigma_c\}}(-1)^{N_l+\kappa(\sigma)}R_{\sigma(1)1}R_{\sigma(2)2}\cdots R_{\sigma(N_l)N_l}. \tag{S2.11}$$

This means, according to (S2.5), that $\Delta_{1,\ldots,N_l}(R)$ is a destabilizing minor and thus, the proof of Lemma S2.1 is finished. ∎

#### Positive intracellular loops induce RD-instabilities of the second kind

Let us consider a gene regulatory network with $N_{ic}\geq 1$ intrcellular gene products and $N_{ec}=N_g-N_{ic}$ extracellular signals, and let $\ell^{+}=\{j_1,\ldots,j_{N_l}\}\subseteq\{i_1,\ldots,i_{N_{ic}}\}$ form a positive intracellular loop, with $N_l\leq N_{ic}$. In accordance with Lemma S2.1, we know that the principal minor of $\boldsymbol{R}=\boldsymbol{R}(0)$ associated to $\ell^{+}$ is a destabilizing minor, as it corresponds to a positive regulatory loop. The following results extends this property to the principal minor of $\boldsymbol{R}$

associated to the whole set of intracellular gene products in the network (i.e., from $\{j_1,\ldots,j_{N_l}\}$ to $\{i_1,\ldots,i_{N_l}\}$):

*Lemma S2.2. Let us consider a gene regulatory network with $N_g>1$ gene products and let $\{i_1,\ldots,i_{N_{ic}}\}$ denote its $N_{ic}<N_g$ intracellular gene products. If such gene network exhibits at least one positive intracellular loop $\ell^+=\{j_1,\ldots,j_{N_l}\}\subseteq\{i_1,\ldots,i_{N_{ic}}\}$, with $N_l\leq N_g$; then, the principal minor of $\boldsymbol{R}=\boldsymbol{R}(0)$ associated to $\{i_1,\ldots,i_{N_{ic}}\}$ is a destabilizing minor of $\boldsymbol{R}$.*

**Proof (of Lemma S2.2).-** Without loss of generality, we can assume that the $N_{ic}$ intracellular gene products in the network are labeled $\{1,\ldots,N_{ic}\}$, with $\ell^+=\{1,\ldots,N_l\}$ (otherwise, we can reorder the labels in the gene network without changing its network topology). Moreover, for the sake of simplicity, we denote $\hat{\mu}_i=m_i\mu_i\left(g_i^*\right)^{m_i-1}$ the degradation term corresponding to gene product $i\in\{1,\ldots,N_{ic}\}$.

From Lemma S2.1, we know that there exists some choice of model parameters such that $(-1)^{N_l}\Delta_{\ell^+}(\boldsymbol{R})<0$. Hence, if $N_l=N_{ic}$, $\Delta_{\{1,\ldots,N_{ic}\}}(\boldsymbol{R})=\Delta_{\ell^+}(\boldsymbol{R})$ is trivially a destabilizing minor of $\boldsymbol{R}$. If $N_l<N_{ic}$, let $i\in\{N_l+1,\ldots,N_{ic}\}$ be any intracellular gene product outside of $\ell^+$. Then, using the minor decomposition formula (Strang, 2016), we can write $\Delta_{\ell^+\cup\{i\}}(\boldsymbol{R})=det\left(\boldsymbol{R}_{\ell^+\cup\{i\}}\right)$ in the form,

$$\Delta_{\ell^+\cup\{i\}}(\boldsymbol{R})=\left(J_{ii}-\hat{\mu}_i\right)\Delta_{\ell^+}(\boldsymbol{R})+\sum_{j\in\ell^+}(-1)^{N_l+1+j}J_{ij}\Delta_{(\ell^+\setminus\{j\})\cup\{i\}}(\boldsymbol{R}). \quad \text{(S2.12)}$$

If we now multiply both sides of (S2.12) by a factor of $(-1)^{N_l+1}$, we get,

$$(-1)^{N_l+1}\Delta_{\ell^+\cup\{i\}}(\boldsymbol{R})=-\left(J_{ii}-\hat{\mu}_i\right)(-1)^{N_l}\Delta_{\ell^+}(\boldsymbol{R})+\sum_{j\in\ell^+}(-1)^{j}J_{ij}\Delta_{(\ell^+\setminus\{j\})\cup\{i\}}(\boldsymbol{R}). \quad \text{(S2.13)}$$

However, we know that there exists some choice of model parameters for which $(-1)^{N_l}\Delta_{\ell^+}(\boldsymbol{R})<0$ and then, complementing such choice with a set of model parameters for gene product $i$ (namely, $J_{ii}$, $\hat{\mu}_i$ and $J_{ij}$ for all $j\in\ell^+$) such that,

$$\left(J_{ii}-\hat{\mu}_i\right)<\frac{\sum_{j\in\ell^+}(-1)^{j}J_{ij}\Delta_{(\ell^+\setminus\{j\})\cup\{i\}}(\boldsymbol{R})}{(-1)^{N_l}\Delta_{\ell^+}(\boldsymbol{R})}, \quad \text{(S2.14)}$$

we get $(-1)^{N_l+1}\Delta_{\ell^+\cup\{i\}}(\boldsymbol{R})<0$ (note that the sign of the terms in the top sum of the right-hand side of (S2.14) alternates depending on the parity of $j$). This means, according to (S2.5), that the principal minor $\Delta_{\ell^+\cup\{i\}}(\boldsymbol{R})$ is a destabilizing minor of $R$. Then, the same argument can be iteratively applied to the remaining gene products outside of $\ell^+\cup\{i\}$ (i.e., gene products $j\in\{N_l+1,\ldots,N_{ic}\}\setminus\{i\}$) until we end up showing that $\Delta_{\{1,\ldots,N_{ic}\}}(\boldsymbol{R})$ is indeed a destabilizing minor. This would finish the proof of Lemma S2.2. ■

So far, parameter choices involved in Lemmas S2.1 and S2.2 correspond exclusively to the reaction part of the reaction-diffusion equations (namely, Jacobian entries $J_{ij}$, degradation coefficients $\mu_i$, degradation orders $m_i$ and steady-state concentrations $g_i^*$). Nevertheless, to understand the true role of positive loops in pattern transformation we must examine the effect that destabilizing minors of $\boldsymbol{R}=\boldsymbol{R}(0)$ have on the full reaction-diffusion matrix $\boldsymbol{R}(\boldsymbol{k}^2)=\boldsymbol{R}-\boldsymbol{k}^2\boldsymbol{D}$, for all wavenumbers $\boldsymbol{k}\in\sigma(\nabla^2)$. To do so, given that $\boldsymbol{D}$ is by definition a positive semidefinite diagonal matrix, we can write the determinant of $\boldsymbol{R}(\boldsymbol{k}^2)$ in terms of the principal minors of $\boldsymbol{R}$ as follows (Wang & Li, 2001),

$$\det(\boldsymbol{R}(\boldsymbol{k}^2))=\det(\boldsymbol{R})+\sum_{j=1}^{N_g-1}\left((-1)^j\sum_{\gamma\in\mathcal{I}_j}\left(\boldsymbol{k}^{2j}\Delta_{\gamma^c}(\boldsymbol{R})\prod_{i\in\gamma}d_i\right)\right)+(-1)^{N_g}\boldsymbol{k}^{2N_g}\prod_{i=1}^{N_g}d_i, \tag{S2.15}$$

where $\mathcal{I}_j=\{\gamma=\{i_1,\dots,i_j\}/1\le i_1\le\dots\le i_j\le N_g\}$ denotes the set of all $j$-element index sequences extracted from $\{1,\dots,N_q\}$; and $\gamma^c=\{1,\dots,N_g\}\backslash\gamma$ is the complementary sequence of $\gamma\in\mathcal{I}_j$.

The following result shows that any gene network containing at least one positive intracellular loop admits at least one choice of model parameters (including diffusion coefficients in $\boldsymbol{D}$) for which it becomes an RD-unstable network of the second kind:

*Proposition S2.3. Let us consider a gene regulatory network with $N_{ic}\ge1$ intracellular gene products and $N_{ec}=N_g-N_{ic}\ge1$ extracellular signals. If the network contains at least one positive intracellular loop $\ell^+=\{j_1,\dots,j_{N_l}\}$, with $N_l\le N_{ic}$; then, there exists a choice of model parameters for which the gene network becomes an RD-unstable network of the second kind.*

**Proof (of Proposition S2.3).-** Without loss of generality, we can assume that the $N_{ic}$ intracellular gene products in the network are labeled $\{1,\dots,N_{ic}\}$, with $\ell^+=\{1,\dots,N_l\}$ (otherwise, we can reorder the labels in the gene network without changing its network topology). Moreover, for the sake of simplicity, we denote by $\hat{\mu}_i=m_i\mu_i(g_i^*)^{m_i-1}$ the full degradation term corresponding to gene product $i\in\{1,\dots,N_g\}$.

From Lemmas S2.1 and S2.2, we know that the principal minor of $\boldsymbol{R}=\boldsymbol{R}(0)$ associated to the full set of intracellular gene products $\{1,\dots,N_{ic}\}$ is a destabilizing minor and consequently, that there exists a choice of model parameters for which,

$$(-1)^{N_{ic}}\Delta_{\{1,\dots,N_{ic}\}}(\boldsymbol{R})<0. \tag{S2.16}$$

First, we will focus on proving that there exists a choice of model parameters for which the gene network is RD-unstable, that is, a choice of model parameters for which there exists at least one unstable wavenumber $\boldsymbol{k}\in\sigma(\nabla^2)$ for which its associated dispersion relation $\det(\lambda\boldsymbol{I}-\boldsymbol{R}(\boldsymbol{k}^2))=0$ admits one solution with positive real part. In this sense, according to the Routh-Hurwitz conditions (Kurosh, 1984; Elsgolts, 2003), $\det(\lambda\boldsymbol{I}-\boldsymbol{R}(\boldsymbol{k}^2))=0$ will admit a solution with positive real part if $(-1)^{N_g}\det(\boldsymbol{R}(\boldsymbol{k}^2))<0$. Hence, it suffices to show that there exists some choice of model parameters for which this latter inequality holds to prove that the corresponding gene network is RD-unstable for such choice of model parameters.

Let $\hat{k} \geq 0$ be a dummy continuous wavenumber variable. The determinant of $\boldsymbol{R}(\hat{k}) = \boldsymbol{R} - \hat{k}\boldsymbol{D}$ can be written in terms of the minors of $\boldsymbol{R}$ using (S2.15). However, since we have $d_i = 0$ for all $i \in \{1, \ldots, N_{ic}\}$, such formula reduces to,

$$\det\left(\boldsymbol{R}\left(\hat{k}\right)\right) = \det\left(\boldsymbol{R}\right) + (-1)^{N_g - N_{ic}} \hat{k}^{N_g - N_{ic}} \Delta_{\{1,\ldots,N_{ic}\}}\left(\boldsymbol{R}\right) \prod_{i=N_{ic}+1}^{N_g} d_i; \tag{S2.17}$$

and then, if we multiply both sides of (S2.17) by a factor of $(-1)^{N_g}$, we get,

$$(-1)^{N_g} \det\left(\boldsymbol{R}\left(\hat{k}\right)\right) = (-1)^{N_g} \det\left(\boldsymbol{R}\right) + (-1)^{N_{ic}} \hat{k}^{N_g - N_{ic}} \Delta_{1,\ldots,N_{ic}}\left(\boldsymbol{R}\right) \prod_{i=N_{ic}+1}^{N_g} d_i. \tag{S2.18}$$

According to (S2.16), there exists some choice of model parameters for which $(-1)^{N_{ic}} \Delta_{\{1,\ldots,N_{ic}\}}(\boldsymbol{R}) < 0$. Thus, given that $\hat{k} \geq 0$ and $d_i > 0$, for all $i \in \{N_{ic}+1, \ldots, N_g\}$, there exists some threshold wavenumber $\widehat{K} \geq 0$ such that $(-1)^{N_g} \det\left(\boldsymbol{R}\left(\hat{k}\right)\right) < 0$ for all $\hat{k} > \widehat{K}$, namely,

$$\widehat{K} = \max\left\{0, \left(\frac{(-1)^{N_{ec}+1} \det(\boldsymbol{R})}{\Delta_{\{1,\ldots,N_{ic}\}}(\boldsymbol{R}) \prod_{i=N_{ic}+1}^{N_g} d_i}\right)^{\frac{1}{N_{ec}}}\right\}, \tag{S2.19}$$

where $N_{ec} = N_g - N_{ic} \geq 1$ is the number of extracellular signals in the network. The argument is then finished by noting that $\sigma\left(\nabla^2\right)$ is a countably, infinite and unbounded set (Baker *et al.*, 2008), so that we can always find some wavenumber $\boldsymbol{k} \in \sigma\left(\nabla^2\right)$ such that $\boldsymbol{k}^2 > \widehat{K}$, which ensures $(-1)^{N_g} \det\left(\boldsymbol{R}\left(\boldsymbol{k}^2\right)\right) < 0$. In other words, for any choice of model parameters enabling (S2.16) (whose existence is given by Lemma S2.2), the gene regulatory network admits at least one unstable wavenumber $\boldsymbol{k} \in \sigma\left(\nabla^2\right)$ and is, therefore, RD-unstable.

Finally, given that $\sigma\left(\nabla^2\right)$ is an unbounded set, for any unstable wavenumber $\boldsymbol{k}_1 \in \sigma\left(\nabla^2\right)$ with $\boldsymbol{k}_1^2 > \widehat{K}$, there always exists some other wavenumber $\boldsymbol{k}_2 \in \sigma\left(\nabla^2\right)$ with $\boldsymbol{k}_2^2 > \boldsymbol{k}_1^2$. This means that $\boldsymbol{k}_2^2 > \widehat{K}$ and ultimately, that $(-1)^{N_g} \det\left(\boldsymbol{R}\left(\boldsymbol{k}_2^2\right)\right) < 0$, implying that $\boldsymbol{k}_2$ is also an unstable wavenumber. As a result, the gene regulatory network does not only admit one, but infinitely many unstable wavenumbers, that is, it is an RD-unstable network of the second kind. This finishes the proof of Proposition S2.3. ■

#### Positive extracellular loops induce RD-instabilities of the first kind

Let us consider a gene regulatory network with $N_{ec} \geq 1$ extracellular signals and $N_{ic} = N_g - N_{ec} \geq 0$ intracellular gene products, and let $L^+ = \{j_1, \ldots, j_{N_l}\}$ form the largest positive extracellular loop in the network (i.e., at least one gene product $j_i \in L^+$ is an extracellular signal). To isolate, and better understand, the role of positive extracellular loops in pattern transformation, we will assume for the remainder of this section that the gene networks under consideration do not contain any positive intracellular loop (the combination of positive extracellular and intracellular loops is considered in the following section). In particular, given this lack of positive intracellular loops in the network, we will have that the principal minor associated to the set of intracellular gene products $\{i_1, \ldots, i_{N_{ic}}\}$ is never a destabilizing minor, that is, for any choice of model parameters, we have,

$$(-1)^{N_{ic}} \Delta_{\{i_1,\ldots,i_{N_{ic}}\}}(\boldsymbol{R}) > 0. \tag{S2.20}$$

The following results shows that, if a gene regulatory network whose positive loops are all extracellular becomes RD-unstable for some choice of model parameters, then it can only be an RD-unstable network of the first kind (i.e., its associated dispersion relation would yield only a finite number of unstable wavenumbers $\boldsymbol{k} \in \sigma(\nabla^2)$):

*Proposition S2.4. Let us consider a gene regulatory network with $N_{ec} \geq 1$ extracellular signals and $N_{ic} = N_g - N_{ec} \geq 0$ intracellular gene products, and assume all positive loops in the network are extracellular, with $L^+ = \{j_1, \ldots, j_{N_l}\}$ denoting the largest one; then, if there exists some choiceof model parameters for which the network becomes RD-unstable, it can only be an RD-unstable network of the first kind.*

**Proof (of Proposition S2.4).-** Let $\mathcal{U} = L^+ \cup \{i_1, \ldots, i_{N_{ic}}\}$ be the union set of loop $L^+$ with the set of all intracellular gene products in the network $\{i_1, \ldots, i_{N_{ic}}\}$, and let $M = |\mathcal{U}|$ be the number of elements in $\mathcal{U}$. Without loss of generality, we can assume that the $M$ gene products in $\mathcal{U}$ are labeled $\{1, \ldots, M\}$, with intracellular gene products being $\{1, \ldots, N_{ic}\}$ (otherwise, we can always reorder the labels in the gene network without changing its network topology). Moreover, for the sake of simplicity, we denote by $\hat{\mu}_i = m_i \mu_i (g_i^*)^{m_i - 1}$ the full degradation term corresponding to gene product $i \in \{1, \ldots, N_g\}$.

From Lemma S2.1, we know that the principal minor of $R = R(0)$ associated to $L^+$ is a destabilizing minor (i.e., there exists some choice of model parameters for which $(-1)^{N_l} \Delta_{L^+}(\boldsymbol{R}) < 0$). Moreover, proceeding as in Lemma S2.2, we can extend such property to the principal minor associated to $\mathcal{U}$, namely, we can find a choice of model parameters for which,

$$(-1)^M \Delta_{\mathcal{U}}(\boldsymbol{R}) < 0. \tag{S2.21}$$

First, we will focus on showing that the positive extracellular loop $L^+$ enables the existence of some choice of model parameters for which the network is RD-unstable (i.e., for which its associated dispersion relation yields at least one unstable wavenumber $\boldsymbol{k} \in \sigma(\nabla^2)$). As in the proof of Proposition S2.3, we will do it using the Ruth-Hurwitz conditions (Kurosh, 1984; Elsgots, 2003) to ensure that $\det(\lambda \boldsymbol{I} - \boldsymbol{R}(\boldsymbol{k}^2)) = 0$ has at least one solution with positive real part by ensuring that $(-1)^{N_g} \det(\boldsymbol{R}(\boldsymbol{k}^2)) < 0$.

Let $\hat{k} \geq 0$ be a dummy continuous wavenumber variable, and let us take $d_i = \delta > 0$ for all extracellular signal $i \in \{N_{ic}+1, \ldots, N_g\}$. The determinant of $\boldsymbol{R}(\hat{k}) = \boldsymbol{R} - \hat{k} \boldsymbol{D}$ can be written in terms of the minors of $R$ using formula (S2.15). Indeed, if $\delta$ is small enough, we have,

$$\det(\boldsymbol{R}(\hat{k})) = \det(\boldsymbol{R}) + (-1)^{N_g - M} (\hat{k}\delta)^{N_g - M} \Delta_{\mathcal{U}}(\boldsymbol{R}) + \mathcal{O}(\delta), \tag{S2.22}$$

where $\mathcal{O}(\delta)$ represents Landau's big-O notation. Then, if we multiply both sides of (S2.22) by a factor of $(-1)^{N_g}$, we get,

$$(-1)^{N_g} det\left(\boldsymbol{R}(\hat{k})\right)=(-1)^{N_g} det(\boldsymbol{R})+(-1)^M (\hat{k}\,\delta)^{N_g-M} \Delta_U(\boldsymbol{R})+O(\delta). \tag{S2.23}$$

According to (S2.21), there exists some choice of model parameters for which $(-1)^M \Delta_U(\boldsymbol{R})<0$. Hence, if $\delta$ is small enough, it is possible to find some lower bound $\widehat{K}_L \geq 0$ such that $(-1)^{N_g} \det\left(\boldsymbol{R}(\boldsymbol{k}^2)\right)<0$ for some wavenumber $\boldsymbol{k} \in \sigma(\nabla^2)$ with $\boldsymbol{k}^2 \geq \widehat{K}_L$, namely,

$$\widehat{K}_L \simeq \max\left\{0, \frac{1}{\delta}\left((-1)^{N_g-M-1}\frac{\det(R)}{\Delta_U(R)}\right)^{\frac{1}{N_g-M}}\right\}. \tag{S2.24}$$

Nonetheless, in contrast to Proposition S2.3, the squared wavenumber $\boldsymbol{k}^2$ cannot be taken arbitrarily bigger than $\widehat{K}_L$ as higher order terms within $O(\delta)$ may become relevant and alter the sign of $(-1)^{N_g}\det\left(\boldsymbol{R}(\boldsymbol{k}^2)\right)$. To illustrate this, let us extract the term of order $\hat{k}^{N_g-N_{ic}}$ from $\mathcal{O}(\delta)$ in (S2.23) and get,

$$\begin{aligned}(-1)^{N_g}\det\left(\boldsymbol{R}(\hat{k})\right)=&\\ &(-1)^{N_g}\det(\boldsymbol{R})+(-1)^M(\hat{k}\,\delta)^{N_g-M}\Delta_U(\boldsymbol{R})+(-1)^{N_{ic}}(\hat{k}\,\delta)^{N_g-N_{ic}}\Delta_{\{1,\dots,N_{ic}\}}(\boldsymbol{R})+\mathcal{O}(\delta),\end{aligned} \tag{S2.25}$$

where, in virtue of (S2.20), $(-1)^{N_{ic}}\Delta_{\{1,\dots,N_{ic}\}}(\boldsymbol{R})>0$ for any choice of model parameters. Thus, given that $M>N_{ic}$, we have that $\delta$ can always be taken small enough so that the third term in (S2.25) is negligible with respect to the second one. However, for any fixed value of $\delta$, this third term (with order $\hat{k}^{N_g-N_{ic}}$ and positive sign) will dominate over the second term (with order $\hat{k}^{N_g-N_{ic}}$ and negative sign) if we let $\hat{k}\to\infty$. In particular, this means that one can always find some upper limit $\widehat{K}_R \geq 0$ such that,

$$(-1)^{N_{ic}}(\hat{k}\,\delta)^{N_g-N_{ic}}\Delta_{\{1,\dots,N_{ic}\}}(\boldsymbol{R}) > \left|(-1)^M(\hat{k}\,\delta)^{N_g-M}\Delta_U(\boldsymbol{R})\right|, \quad \text{for all } \boldsymbol{k}^2 \geq \widehat{K}_R. \tag{S2.26}$$

Notice that it can be the case that the interval $\left[\widehat{K}_L, \widehat{K}_R\right)$ is so narrow that no wavenumber $\boldsymbol{k} \in \sigma(\nabla^2)$ satisfies $\boldsymbol{k}^2 \in \left[\widehat{K}_L, \widehat{K}_R\right)$. In this case, the gene network under consideration would not be RD-unstable. However, if the interval is wide enough to admit at least one wavenumber $\boldsymbol{k} \in \sigma(\nabla^2)$ such that $\boldsymbol{k}^2 \in \left[\widehat{K}_L, \widehat{K}_R\right)$ (i.e., if the gene network is RD-unstable), then there can only be a finite number of such wavenumbers: $\sigma(\nabla^2)$ is an unbounded set (Baker *et al.*, 2008) and hence, there will always be some big wavenumber $\boldsymbol{k} \in \sigma(\nabla^2)$ for which $\boldsymbol{k}^2 > \widehat{K}_R$. This means that, if the gene network is RD-unstable for some choice of model parameters, it can only be RD-unstable of the first kind and thus, the proof of Proposition S2.4 is finished. ■

Finally, two observations are worth being made. First, note that tuning model parameters to get an RD-unstable network of the first kind generally requires a much finer degree of adjustment than that needed to get an RD-unstable network of the second kind. In the former case, one must find some wavenumber $\boldsymbol{k}\in\sigma\left(\nabla^2\right)$ such that $\boldsymbol{k}^2\in\left[\widehat{K}_L,\widehat{K}_R\right]$, where $\widehat{K}_L$ and $\widehat{K}_R$ are given in the proof of Proposition S2.4. Conversely, in the latter case it suffices to select a sufficiently large $\boldsymbol{k}\in\sigma\left(\nabla^2\right)$ such that $\boldsymbol{k}^2\geq\widehat{K}$, with $\widehat{K}$ given by (S2.19).

On the other hand, it is worth recalling that although the parameter selection employed for the proof of Proposition S2.4 may appear rather artificial, instances of RD-unstable networks of the first kind arising from less convoluted parameter choices have been reported both in this article (e.g., Figures 8 and 9 in the main text), and in the literature (e.g., Gierer & Meinhardt, 1972; Murray, 2003; Levine & Rappel, 2005).

##### Combination of positive extracellular and intracellular loops

Let us consider a gene regulatory network with $N_{ic}\geq1$ intracellular gene products and $N_{ec}=N_g-N_{ic}\geq1$ extracellular signals, and let $\ell^+=\{i_1,\ldots,i_{N_l}\}$ and $L^+=\{j_1,\ldots,j_{N_l'}\}$ represent two positive intracellular and extracellular loops, respectively. Given that the network contains at least one positive intracellular loop, Proposition S2.3 guarantees the existence of some choice of model parameters for which the network becomes RD-unstable of the second kind. In such parameter set, the intracellular loop $\ell^+$ is effectively overweighted relative to the extracellular loop $L^+$, thereby exerting a stronger influence on the corresponding pattern transformation.

However, model parameters may alternatively be tuned so as to satisfy $(-1)^{N_{ic}}\Delta_{i_1,\ldots,i_{N_{ic}}}(\boldsymbol{R})>0$. In this case, Proposition S2.4 ensures that this parameter set can be completed to a choice of model parameters for which the network becomes RD-unstable of the first kind. Under this conditions, the extracellular loop $L^+$ becomes overweighted relative to the intracellular loop $\ell^+$, and consequently dominates the reaction-diffusion dynamics, leading to a bigger influence on the corresponding pattern transformation.

Thus, gene regulatory networks that combine positive extracellular and intracellular loops can exhibit RD-instabilities of either types, depending on the specific choice of model parameters. As a result, such networks have access to the full spectrum of pattern transformations associated with both classes of RD-unstable networks.

#### S3: Gradients of extracellular signals from spikes

In this section, we analyze the pattern transformations produced by a simple two-node gene network consisting of a self-activating intracellular gene product, denoted $1$, that activates an extracellular signal, denoted $A$, when acting on a 1D spike initial pattern. As described in the main text, extracellular signals in this situation are secreted from the spike and form exponential gradients whose slopes depend on model parameters such as their diffusion and degradation coefficient. This subsection therefore provides a detailed derivation of equation (23) in the main text.

We begin by solving for a stationary concentration of intracellular gene product $1$. The concentration dynamics of this intracellular gene product for each $x$ is given by the following initial value problem (recall equation (1) in the main text),

$$\begin{cases} \partial_t g_1(t,x) = f_1(g_1(t,x)) - \mu_1 g_1(t,x), \\ g_1(0,x) = A_1 \operatorname{rect}\left(\frac{|x-x_0|}{L}\right), \end{cases} \tag{S3.1}$$

where $A_1 > 0$ is the small amplitude of the spike, $L > 0$ is its width, and $\operatorname{rect}(y)$ denotes the rectangular functions, that is, $\operatorname{rect}(y) = 1$, if $-1 \le y \le 1$, and $\operatorname{rect}(y) = 0$, otherwise (Abramowitz & Stegun, 1964). Without loss of generality, we can assume that the spike is centered at the origin $x_0 = 0$ (otherwise, we can simply consider a spatial translation of the problem).

For any position outside the spike (i.e., $x \notin [-L, L]$), we have $g_1(t,x) = 0$ for all $t > 0$, since no production of gene product $1$ can ever occur without any non-zero initial concentration and $1$ is regulated by no extracellular signal. On the contrary, for any position inside the spike $x \in [-L, L]$, we have, up to a linear level,

$$g_1(t,x) = A_1 e^{(J_{11} - \mu_1)t}, \tag{S3.2}$$

where $J_{11} = \partial_{g_1} f_1(0)$. Then, if $\mu_1 < J_{11}$, the concentration of gene product $1$ inside the spike will grow away from $0$ until it eventually saturates at some $G_1 > 0$ due to the effect of the non-linear terms of $f_1(g_1)$ (recall requirements R2 and R4 in the main text). As a result, the resulting pattern of gene product $1$ is a spike centered at the same position and a different height,

$$\bar{g}_1(x) = G_1 \operatorname{rect}(x/L). \tag{S3.3}$$

Now, we turn to the analysis of pattern transformations in extracellular signal $A$. If we assume that gene product $1$ reaches its stationary concentration fast enough, the concentration dynamics of signal $A$ is given by (recall equation (1) in the main text),

$$\begin{cases} \partial_t g_A(t,x) - d_A \partial_{xx}^2 g_A(t,x) = -\mu_A g_A(t,x) + \hat{G}_1 \operatorname{rect}\left(\frac{x}{L}\right), \\ \lim_{|x| \to \infty} g_A(t,x) = 0, \text{ and } g_A(0,x) = 0, \end{cases} \tag{S3.4}$$

where $\hat{G}_1 = f_A(G_1)$. Solutions to a partial differential equation like (S3.4) can be written in the form $g_A(t,x) = g_A^{\text{Hom}}(t,x) + \bar{g}_A(x)$ (Sneddon, 1957), where $g_A^{\text{Hom}}(t,x)$ is a solution to the transient homogeneous equation,

$$\begin{cases} \partial_t g_A^{\text{Hom}}(t,x) - d_A \partial_{xx}^2 g_A^{\text{Hom}}(t,x) + \mu_A g_A^{\text{Hom}}(t,x) = 0; \\ \lim_{|x| \to \infty} g_A^{\text{Hom}}(t,x) = 0, \quad \text{and} \quad g_A^{\text{Hom}}(0,x) = -\bar{g}_A(x); \end{cases} \tag{S3.5}$$

and $\bar{g}_A(x)$ is a solution to the stationary equation,

$$\begin{cases} \partial_{xx}^2 \bar{g}_A(x) = \frac{\widehat{G}_1}{d_A} \operatorname{rect}\left(\frac{x}{L}\right) - \frac{\mu_A}{d_A} \bar{g}_A(x), \\ \lim_{|x| \to \infty} \bar{g}_A(t, x) = 0. \end{cases} \tag{S3.6}$$

However, for the purposes of this article we are just interested in the formation of stationary solutions of system (S3.4) and thus, we can assume that $\lim_{t \to \infty} g_A^{\text{Hom}}(t, x) = 0$ and ignore (S3.5). Regarding the stationary equation, note that (S3.6) is invariant to sign reflection and so, $\bar{g}_A(x)$ must be an even function (i.e., $\bar{g}_A(x) = \bar{g}_A(-x)$). Consequently, we must have $\partial_x \bar{g}_A(0) = 0$, and we can restrict our search for a closed form of $\bar{g}_A(x)$ to only the right half-line (i.e., $x \geq 0$) by adding the extra zero flux boundary condition $\partial_x \bar{g}_A(0) = 0$ to equation (S3.6).

In this sense, for positions within the spike (i.e., $0 \leq x \leq L$), $\operatorname{rect}(x/L) = 1$ and the stationary equation (S3.6) reads,

$$\begin{cases} \partial_x^2 \bar{g}_A(x) = \frac{\widehat{G}_1}{d_A} - \frac{\mu_A}{d_A} \bar{g}_A(x), \\ \partial_x \bar{g}_A(0) = 0, \text{ and } \bar{g}_A(L) = B, \end{cases} \tag{S3.7}$$

where $B \in \mathbb{R}$ is a splicing constant that we must later find. Solutions to (S3.7) are given by downward-facing hyperbolic cosines of the form,

$$\bar{g}_A(x) = \left(B - \frac{\widehat{G}_1}{\mu_A}\right) \frac{\cosh(\omega_A x)}{\cosh(\omega_A L)} + \frac{\widehat{G}_1}{\mu_A}, \tag{S3.8}$$

where $\omega_A = \sqrt{\mu_A / d_A}$. On the other hand, for positions outside the spike (i.e., $x > L$), $\operatorname{rect}(x/L) = 0$ and then, equation (S3.6) reduces to,

$$\begin{cases} \partial_{xx}^2 \bar{g}_A(x) = -\frac{\mu_A}{d_A} \bar{g}_A(x), \\ \bar{g}_A(L) = B, \text{ and } \lim_{x \to \infty} \bar{g}_A(x) = 0, \end{cases} \tag{S3.9}$$

where $B$ is the same splicing constant as in (S3.7). Then, solutions to (S3.9) are given by a negative exponential of the form,

$$\bar{g}_A(x) = B e^{-\omega_A (x-L)}. \tag{S3.10}$$

By definition, solutions (S3.8) and (S3.10) match continuously at $x = L$. However, we can push this matching to be not only continuous but also continuously differentiable (i.e., class $\mathcal{C}^1$) by equating the first derivative of both functions at $x = L$, and solving for a closed form of the splicing constant $B$. Hence, after some algebra, we get,

$$B = \frac{K_A}{2} \left(1 - e^{-2\omega_A L}\right), \tag{S3.11}$$

where $K_A = \widehat{G}_1 / \mu_A$. Finally, if we substitute (S3.11) back into (S3.8) and (S3.10), we get that the stationary concentration of extracellular signal $A=2$ or $3$ is given by,

$$\bar{g}_A(x) = \begin{cases} K_A\left(1-e^{-\omega_A L}\cosh(\omega_A x)\right), & \text{if } x \in [-L, L]; \\ \dfrac{K_A}{2}\left(1-e^{-2\omega_A L}\right)e^{-\omega_A(|x|-L)}, & \text{otherwise.} \end{cases} \tag{S3.12}$$

As we can see, the stationary concentration of extracellular signal $A$ exhibits a unique, symmetric concentration peak centered at the initial spike ($x_0 = 0$ in our case), and two gradients of exponential decay on both sides of the spike. The steepness of such gradients is controlled by parameter $\omega_A$ (i.e., the square root of the ratio between the degradation rate and the diffusion rate) in the sense that the smaller $\omega_A$ is, the steeper the corresponding gradient. Similarly, constant $C$ in equation (23) of the main text corresponds to,

$$C = \frac{K_A}{2}\left(1-e^{-2\omega_A L}\right) = \frac{\widehat{G}_1}{2\mu_A}\left(1-e^{-2\sqrt{\frac{\mu_A}{d_A}}\,L}\right). \tag{S3.13}$$

Here, we focus exclusively on the 1D pattern transformations in signal $A$. However, equation (S3.4) corresponds to the classical Helmholtz equation, whose solutions in higher dimensions are well known and extensively documented in the literature (Sommerfeld, 1949; Stakgold & Holst, 2011). Hence, we know that in 2D the signal concentration gradient outside the spike in (S3.12) decays as $\sim e^{-\|\boldsymbol{x}\|_2} / \sqrt{\|\boldsymbol{x}\|_2}$, where $\|\boldsymbol{x}\|_2$ denotes the Euclidean norm; whereas in 3D, the corresponding gradient decays as $\sim e^{-\|\boldsymbol{x}\|_2} / \|\boldsymbol{x}\|_2$.

#### S4. Pattern transformations in diamond gene networks for spike initial patterns

In this section, we discuss some details of the 1D pattern transformations that a diamond gene network is able to produce when acting on spike initial pattern. We focus on the diamond network because its topology is simple enough to allow for a precise analysis of the pattern-formation process, while remaining versatile enough to serve as a representative example of many of the features exhibited by more general hierarchical networks.

By definition, a diamond network consists of five distinct gene products (see Figure 11 in the main text). Specifically, it comprises an upstream intracellular gene product (denoted $1$) that self-activates and activates two different extracellular signals (denoted $2$ and $3$). These signals regulate a common downstream intracellular gene product (denoted $4$) in such a way that $2$ activates $4$ whereas $3$ inhibits it. Finally, a fifth intracellular gene product (denoted $5$) is directly activated by gene product $4$. Moreover, we will assume that all gene products suffer from linear degradation (i.e., $m_i = 1$ for all $i \in \{1,2,3,4,5\}$).

As we saw in the previous section, extracellular signals $2$ and $3$ develop a resulting pattern consisting of a unique concentration peak centered at the spike and with exponential decay. Although this does not constitute a non-trivial pattern transformation (it fails to meet requirements (S1.5) and (S1.6)), the fact that the resulting pattern is heterogeneous, and spreads along the domain due to signal diffusion, allows the emergence of non-trivial pattern transformations in downstream gene products, namely, 4 and 5.

In this sense, the concentration dynamics of intracellular gene product 4 for each $x$ is given by the initial value problem (recall equation (1) in the main text),

$$\begin{cases} \partial_t g_4(t,x) = f_4(g_2(t,x), g_3(t,x)) - \mu_4 g_4(t,x), \\ g_4(0,x) = 0. \end{cases} \tag{S4.1}$$

Taking the time derivative in (S4.1) equal to zero, we can easily see that the stationary concentration of gene product 4 is given by,

$$\bar{g}_4(x) = \frac{1}{\mu_4} f_4(\bar{g}_2(x), \bar{g}_3(x)), \tag{S4.2}$$

where we have assumed that both extracellular signals reach their stationary concentrations $\bar{g}_2(x)$ and $\bar{g}_3(x)$ in (S3.12) fast enough. From requirement R5 in the main text, we know that $f_4(g_2, g_3)$ is monotonously increasing with respect to $g_2$ and monotonously decreasing with respect to $g_3$. Then, if we focus on positions to the right of the spike (i.e., $x > L$) where both $\bar{g}_2(x)$ and $\bar{g}_3(x)$ display an exponential gradient, model parameters can be tuned so that the inhibition of 4 by 3 is larger than its activation by 2 close to the spike, and smaller further away. As a result, a concentration peak in the stationary concentration of 4 will emerge at some position $\hat{x} > L$ (and, by symmetry of (S3.12), also at $-\hat{x} > -L$).

To illustrate this, let us consider the simple case in which $f_4(g_A, g_B) = W_{24} g_2 - W_{34} g_3$ is a linear function where $W_{24}, W_{34} > 0$ represent the weights of the regulations that 2 and 3 exert on 4, respectively. In this scenario, (S4.2) reduces to,

$$\bar{g}_4(x) = \frac{1}{2\mu_4}\left(W_{24} K_2 \left(1 - e^{-2\omega_2 L}\right) e^{-\omega_2(x-L)} - W_{34} K_3 \left(1 - e^{-2\omega_3 L}\right) e^{-\omega_3(x-L)}\right), \tag{S4.3}$$

for all positions to the right of the spike (i.e., $x > L$). Hence, if we take the first spatial derivative of (S4.3), we get,

$$\partial_x \bar{g}_4(x) = \frac{-1}{2\mu_4}\left(W_{24} K_2 \omega_2 \left(1 - e^{-2\omega_2 L}\right) e^{-\omega_2(x-L)} - W_{34} K_3 \omega_3 \left(1 - e^{-2\omega_3 L}\right) e^{-\omega_3(x-L)}\right); \tag{S4.4}$$

and thus, taking $\partial_x \bar{g}_4(x) = 0$ in (S4.4), we can derive a closed form for a unique critical point of $\bar{g}_4(x)$on the right-hand side of the initial spike given by,

$$\hat{x} = L + \frac{1}{\omega_3 - \omega_2} \log\left(\frac{W_{34} K_3 \omega_3}{W_{24} K_2 \omega_2} \frac{1 - e^{-2\omega_3 L}}{1 - e^{-2\omega_2 L}}\right). \tag{S4.5}$$

However, this critical point is only formal, as its existence relies on a choice of model parameters for which $\hat{x} > L$. In this sense, after some computations on (S4.5), we can see that this is the case for any parameter set that satisfies,

$$\omega_3 > \omega_2, \qquad \text{(S4.6)}$$

so that the fraction in the second term of (S4.5) is positive; and,

$$\frac{W_{34} K_3 \omega_3}{W_{24} K_2 \omega_2} > \frac{1-e^{-2\omega_2 L}}{1-e^{-2\omega_3 L}}, \qquad \text{(S4.7)}$$

so that the logarithm in the second term of (S3.18) is also positive. Then, if conditions (S4.6) and (S4.7) hold, we have $\partial_x \bar{g}_4(x)>0$ for all positions between $x=L$ and $x=\hat{x}>L$; while$\partial_x \bar{g}_4(x)<0$ for all positions larger than $x=\hat{x}$. Consequently, $\hat{x}$ is a concentration peak of $\bar{g}_4(x)$and thus, we can conclude that gene product undergoes a non-trivial pattern transformation given by the emergence of such concentration peak to the right of the initial spike. In fact, given the parity of equation (S3.12),we know that a symmetric concentration peak of $\bar{g}_4(x)$ also emerges at $x=-\hat{x}$, to the left of the spike.

Finally, the concentration dynamics of intracellular gene product $5$ for each position $x$ is given by the initial value problem (recall equation (1) in the main text),

$$\begin{cases} \partial_t g_5(t,x)=f_5(g_4(t,x))-\mu_5 g_5(t,x), \\ g_5(0,x)=0. \end{cases} \qquad \text{(S4.8)}$$

Then, if we set the time derivative in (S4.8) equal to zero, we can easily see that the stationary concentration of gene product $5$ is given by,

$$\bar{g}_5(x)=\frac{1}{\mu_5} f_5(\bar{g}_4(x)), \qquad \text{(S4.9)}$$

where we have assumed that gene product 4 reaches its stationary concentration $\bar{g}_4(x)$ fast enough. We know from R5 in the main text, that $f_5(g_4)$ must preserve the monotony of $\bar{g}_4(x)$ (i.e., it must increase where $\bar{g}_4(x)$ increases, and decrease where $\bar{g}_4(x)$ decreases). As a result, the resulting pattern of $5$ replicates the resulting pattern of 4 in the sense that concentration peaks and valleys in $\bar{g}_5(x)$ are located at the same positions than those of $\bar{g}_4(x)$. Nonetheless, by tuning the functional form of $f_5(g_4)$, as well as the value of model parameters like $\mu_5$, we can make the concentration peaks are valleys of $5$ higher, deeper or steeper than those of 4.

All in all, we have proven that a simple diamond H network is enough to transform a spike initial pattern into a resulting pattern with two symmetric peaks of concentrations on both sides of the initial pattern. Moreover, as suggested by equations (S4.5) and (S4.9), model parameters and regulation function $\boldsymbol{f}(\boldsymbol{g})$ can be tuned to make such concentration peaks appear at any position out of the spike, and have any desired height and steepness.

#### S5: Peak addition and subtraction in H gene regulatory networks

In this section, we are going to show how two patterned gene products in a hierarchic network can lead to more complex patterns in downstream gene products without requiring to additional extracellular signals. This is can be achieved by two simple mechanisms that include positive (for 'addition' of patterns) and negative (for 'subtraction' of patterns) intracellular regulations between the patterned gene products.

Let us consider two gene products $i$ and $j$ exhibiting heterogeneous resulting patterns $\bar{g}_i(\boldsymbol{x})$ and $\bar{g}_j(\boldsymbol{x})$ that directly regulate intracellular gene product $k$. If no other gene product in the network regulates $k$, its concentration dynamics for each $\boldsymbol{x}$ is given by,

$$\partial_t g_k(t,\boldsymbol{x})=f_k(g_i(x),g_j(t,\boldsymbol{x}))+\mu_k g_k(t,\boldsymbol{x}); \tag{S5.1}$$

and then, assuming that $i$ and $j$ reach their respective stationary concentrations fast enough, we can take the time derivative in (S5.1) equal to zero and get that the stationary concentration of gene product $k$ is given by,

$$\bar{g}_k(\boldsymbol{x})=\frac{1}{\mu_k} f_k(\bar{g}_i(\boldsymbol{x}),\bar{g}_j(\boldsymbol{x})). \tag{S5.2}$$

First, we consider the case of the peak addition mechanism, that is, we assume that both $i$ and $j$ positively regulate gene product $k$ (see Figure 10 in the main text). In this case, according to requirement R5 in the main text, $f_k(g_i,g_j)$ is component-wise monotonously increasing with respect to both $g_i$ and $g_j$. Hence, we know that the stationary concentration of gene product $k$ will increase wherever the stationary concentrations of both $i$ and $j$ simultaneously increase; and it will decrease wherever those of $i$ and $j$simultaneously decrease. As a result, if both $\bar{g}_i(\boldsymbol{x})$ and $\bar{g}_j(\boldsymbol{x})$ have a concentration peak at the same position $\hat{\boldsymbol{x}}$, then $\bar{g}_k(\boldsymbol{x})$will also exhibit a peak at $\hat{\boldsymbol{x}}$.

Ambiguity appears in those cases for which the peaks of concentration of $i$ and $j$ do not coincide at the same spatial position. In this sense, let us assume that the stationary concentrations of gene products $i$ and $j$ exhibit a total amount of $N$ and $M$ concentration peaks along the domain, respectively; and let us focus in two consecutive such peaks, that is, let us consider a concentration peak in $\bar{g}_i(\boldsymbol{x})$ located at a given position $\hat{\boldsymbol{x}}_i$ and a concentration peak in $\bar{g}_j(\boldsymbol{x})$ at a different position $\hat{\boldsymbol{x}}_j \neq \hat{\boldsymbol{x}}_i$ such that neither $\bar{g}_i(\boldsymbol{x})$ nor $\bar{g}_j(\boldsymbol{x})$ exhibit an intermediate concentration peak along the line going from $\hat{\boldsymbol{x}}_i$ to $\hat{\boldsymbol{x}}_j$. In this setting, if $\hat{\boldsymbol{x}}_i$ and $\hat{\boldsymbol{x}}_j$ are far enough from each other and both concentration peaks are steep enough so that the influence of $\bar{g}_j(\boldsymbol{x})$ (resp., $\bar{g}_i(\boldsymbol{x})$) is negligible near $\hat{\boldsymbol{x}}_i$ (resp., $\hat{\boldsymbol{x}}_j$); then, $f_k(g_i,g_j)$ will behave as a function of only $g_i$ (resp., $g_j$) in a vicinity of $\hat{\boldsymbol{x}}_i$ (resp., $\hat{\boldsymbol{x}}_j$) and thus, $\bar{g}_k(\boldsymbol{x})$ will exhibit one concentration peak at $\hat{\boldsymbol{x}}_i$ and another concentration peak at $\hat{\boldsymbol{x}}_j$. Moreover, if this condition holds for any pair of consecutive peaks of $\bar{g}_i(\boldsymbol{x})$ and $\bar{g}_j(\boldsymbol{x})$, $\bar{g}_k(\boldsymbol{x})$ will exhibit a total amount of $N+M$ concentration peaks along the domain (see Figure 10 in the main text). On the contrary, if two such consecutive peaks of $i$ and $j$ are too close to each other, the expected concentration peaks of $k$ may merge into just one located along the line from $\hat{\boldsymbol{x}}_i$ to $\hat{\boldsymbol{x}}_j$ and then, the total number of concentration peaks that $k$ will exhibit along the domain is strictly smaller than$N+M$.

On the other hand, let us consider the case of the peak subtraction mechanism, that is, let us assume that $i$ activates $k$ while $j$ inhibits it (see Figure 10 in the main text). In this setting, according to requirement R5 in the main text, $f_k(g_i, g_j)$ is monotonously increasing with respect to $g_i$ and monotonously decreasing with respect to $g_j$. As in the case of peak addition, let us assume that the stationary concentrations of gene products $i$ and $j$ display a total amount of $N$ concentration peaks each. Also, let us assume that two such peaks of concentration coincide at the same position $\hat{\boldsymbol{x}}$. Then, if the activation of $i$ on $k$ is much stronger than the inhibition of $j$ on $k$, $\bar{g}_k(\boldsymbol{x})$ will portray a concentration peak at $\hat{\boldsymbol{x}}$. Conversely, if the inhibition of $j$ on $k$ is much stronger than the activation of $i$ on $k$, no concentration peak of $k$ will emerge at $\hat{\boldsymbol{x}}$. Thus, if the former situation takes place at all possible pairs of concentration peaks of $i$ and $j$, $\bar{g}_k(\boldsymbol{x})$ will display a total amount of $N$ concentration peaks along the domain. However, if the latter situation takes place for all possible pairs of peaks, $\bar{g}_k(\boldsymbol{x})$ will retract back to a homogeneous concentration as all possible peaks of concentration will be eventually swept away by the inhibition of $j$.

However, as we have seen in section S3, under some specific conditions on function $\boldsymbol{f}$ and specific parameter values, the inhibition that $j$ exerts on $k$ can split the emerging concentration peak in two in such a way that, on a 1D domain, $\bar{g}_k(x)$ displays two symmetric concentration peaks on both sides of $\hat{x}$ (this pair of symmetric peaks becomes a ring around $\hat{\boldsymbol{x}}$ in 2D, and a hollow sphere in 3D). Moreover, this split would also be the case even if the concentration peaks in $\bar{g}_i(x)$ and $\bar{g}_j(x)$ did not coincide at the same position, but stand in close positions (although in this case the height of the emerging peaks in $\bar{g}_k(x)$ would be uneven). Anyhow, if this splitting of peaks occurs for all possible pairs of concentration peaks of $i$ and $j$, $\bar{g}_k(x)$ will exhibit a total amount of $2N$ peaks along the 1D domain (in higher dimensions, each pair of peaks reduces to one single ring or sphere and hence, the total amount of structures in $\bar{g}_k(x)$ reduces to $N$).

#### S6: Numerical simulations

In this supplementary section, we provide the necessary information to reproduce the numerical simulations leading to all resulting patterns presented in the various figures along the main text and supplementary information of this article. Hence, we first offer a detailed description of the Maini-Miura model (Miura and Maini, 2004; Marcon *et al.*, 2016), which serves as the basis for the reaction-diffusion equations in all our simulations (i.e., equations (1) of the main text); and we list the specific parameter values applied in each case. All parameters presented in this supplementary section are expressed in arbitrary units (a.u.).

##### The Maini-Miura model

In all our simulations, the underlying reaction-diffusion equations correspond to the Maini-Miura model (Miura and Maini, 2004; Marcon *et al.*, 2016), or slight modifications of it. The governing equations of this simple model of pattern formation are given by,

$$\partial_t \boldsymbol{g}(t,\boldsymbol{x}) = \boldsymbol{W}^T(\boldsymbol{g}(t,\boldsymbol{x}) - \boldsymbol{g}^*) - \boldsymbol{S}(\boldsymbol{g}(t,\boldsymbol{x}) - \boldsymbol{g}^*)^3 - \boldsymbol{M}(\boldsymbol{g}(t,\boldsymbol{x}) - \boldsymbol{g}^*) + \boldsymbol{D}\nabla^2 \boldsymbol{g}(t,\boldsymbol{x}), \qquad \text{(S6.1)}$$

where $\boldsymbol{g}(t,\boldsymbol{x})\in \mathbb{R}^{N_g}$ is the vector of gene product concentrations at time $t\geq 0$ and position $x$; $\boldsymbol{g}^{*}\in \mathbb{R}^{N_g}$ is a homogeneous steady state concentration; $\boldsymbol{W}\in \mathbb{R}^{N_g\times N_g}$ is the weight matrix of the corresponding gene regulatory network; $\boldsymbol{S}=\mathrm{diag}(\sigma_1,\ldots,\sigma_{N_g})$ is the cubic saturation matrix; $\boldsymbol{M}=\mathrm{diag}(\mu_1,\ldots,\mu_{N_g})$ is the linear degradation matrix; and $\boldsymbol{D}=\mathrm{diag}(d_1,\ldots,d_{N_g})$ is the diffusion matrix. As in equation (1) of the main text, $\boldsymbol{g}^3$ denotes component wise exponentiation.

Function $\boldsymbol{f}(\boldsymbol{g})$ in equation (1) of the main text corresponds to the combination of terms $\boldsymbol{W}^T(\boldsymbol{g}(t,\boldsymbol{x})-\boldsymbol{g}^{*})-\boldsymbol{S}(\boldsymbol{g}(t,\boldsymbol{x})-\boldsymbol{g}^{*})^3+\boldsymbol{M}\boldsymbol{g}^{*}$ in equation (S6.1). Given its polynomial structure, it is straightforward to check that it satisfies requirements R1 and R3 in the main text and, if $\sigma_i>0$ for at least one gene product, also requirement R2. The Jacobian matrix of $\boldsymbol{f}(\boldsymbol{g})$ at $\boldsymbol{g}^{*}$ coincides with the transpose weight matrix (i.e., $\boldsymbol{J}_f(\boldsymbol{g}^{*})=\boldsymbol{W}^T$) and then, $\boldsymbol{f}(\boldsymbol{g})$ satisfies requirement R5 in the main text by continuity (at least in a neighborhood of $\boldsymbol{g}^{*}$). Regarding requirement R4 in the main text, concentration $\boldsymbol{g}(t,\boldsymbol{x})$ is the Maini-Miura model is always bounded from above because for large values of $\boldsymbol{g}(t,\boldsymbol{x})$ both degradation terms in (S6.1) become negative. Also, the entries in $\boldsymbol{g}^{*}$ can be tuned to avoid $\boldsymbol{g}(t,\boldsymbol{x})<0$ for all time $t\geq 0$ and all positions $\boldsymbol{x}$.

Despite its many simplifications, the Maini-Miura model offers a practical framework for the study of pattern formation governed gene regulatory networks, as its formulation enables a direct manipulation of gene network topology by simply changing the entries of the weight matrix $\boldsymbol{W}$. This adaptability makes the Maini-Miura model particularly suitable for the study of gene regulatory networks comprising a large number of gene products, unlike classical models such as the Schnackenberg (Schnackenberg, 1979; Murray, 2003) or Gierer-Meinhardt models (Gierer & Meinhardt, 1972; Meinhardt, 1982), which are constrained to a very specific two-node Turing gene network.

In our simulations, we build spike, spike-homogeneous and noise-homogeneous initial patterns using $\boldsymbol{g}^{*}$ as their basal concentrations. For spike- and noise-homogeneous initial patterns we consider $g_i^{*}>0$ big enough so that $g_i(t,\boldsymbol{x})$ never becomes negative. For spike initial patterns, we take $\boldsymbol{g}^{*}\equiv 0$ and clamp possible negative concentration by introducing a smooth step-function in the first term of (S6.1), namely,

$$\boldsymbol{f}(\boldsymbol{g})=\boldsymbol{\Phi}^{(\varepsilon)}\left(\boldsymbol{W}^T(\boldsymbol{g}(t,\boldsymbol{x})-\boldsymbol{g}^{*})\right), \tag{S6.2}$$

where each entry $\Phi_{ij}^{(\varepsilon)}(z)=(1+e^{-z/\varepsilon})^{-1}$ is a narrow sigmoid function, with $\varepsilon>0$ arbitrarily small. Note that the inclusion of the smooth step function $\boldsymbol{\Phi}^{(\varepsilon)}$ in $\boldsymbol{f}(\boldsymbol{g})$ is consistent with requirements R1-R5 in the main text. Moreover, if we choose $\varepsilon$ small enough, it acts as an effective clamping term in model equations that prevents the occurrence of negative concentrations during the simulations.

In a similar spirit, several figures in the main text and the supplementary information explicitly require different components of $\boldsymbol{f}(\boldsymbol{g})$ to adopt different functional forms (e.g., Figure 7 in the main text, evaluating conditions RH1 and RH2). In such simulations, only some specific components of $\boldsymbol{f}(\boldsymbol{g})$ are modified, leaving all remaining terms of equation (S6.1) unchanged.

Each such modification is listed bellow, together with the corresponding set of model parameters that lead to the pattern transformation under consideration.

#### Parameter values for patterning simulations

All simulations in this article have been performed using a Forward-Euler algorithm (Grossmann and Ross, 2007) on a 1-dimensional interval $[0,50]$ split into a line array of 500 cells with a constant spatial step (i.e., distance between cell centers) of $h=0.1$, or a 2-dimensional square $[0,50]\times[0,50]$ split into a square ensemble of $250\times 250$ cells with a constant spatial step of $h=0.1$. The time step $\tau$ of the algorithm is adjusted for every simulation in such away that the convergence condition of the Forward-Euler algorithm is satisfied for all gene products in the gene network (i.e., $2^n \tau d_i < h^2$ for all $i$ and $n=1,2$). This results in a $\boldsymbol{D}$-dependent time step of the form,

$$\tau=\tau(\boldsymbol{D})=\frac{h^2}{(2^n+0.1)max\{d_1,\ldots,d_{N_g}\}}, \quad \text{for}\, n=1,2. \qquad \text{(S6.3)}$$

All simulations start with an initial pattern built around the equilibrium point of equation (S6.1), that is, $\boldsymbol{g}^*=\left(g_1^*,\ldots,g_{N_g}^*\right)^T$. Spike ($\boldsymbol{g}^*=\boldsymbol{0}$) and spike-homogeneous ($\boldsymbol{g}^*\neq\boldsymbol{0}$) initial conditions centered at $\boldsymbol{x}_0\in\mathbb{R}^n$ are given by rectangular functions of the form,

$$g_i(0,\boldsymbol{x})=g_i^*+A_i\ \mathrm{rect}\left(\frac{\|\boldsymbol{x}-\boldsymbol{x}_0\|_2}{L}\right), \qquad \text{(S6.4)}$$

where $A_i>0$ is the amplitude of the spike; $L>0$ is the width of the spike (equal for all gene products); $\|\cdot\|_2$ denotes the euclidean norm; and $\mathrm{rect}(y)=1$, if $-1\leq y\leq 1$, while $\mathrm{rect}(y)=0$, elsewhere (Abramowitz and Stegun, 1964). On the other hand, noise-homogeneous initial patterns are given by $g_i(0,\boldsymbol{x})=g_i^*+A_i \mathrm{wn}_i(\boldsymbol{x})$, where $A_i>0$ is the amplitude of the noise and $\mathrm{wn}_i(\boldsymbol{x})$ is a random value uniformly chosen in the interval $[-1,1]$ for each different position $\boldsymbol{x}$ and each different gene product $i\in\{1,\ldots,N_g\}$.

##### *Figure 2 in the main text*

Figure 2 in the main text shows the pattern transformations that a 4-node hierarchical gene regulatory network produces on a spike ($\boldsymbol{g}^*=\boldsymbol{0}$, $x_0=25$, $\boldsymbol{A}=(0.05,0,0,0)^T$, $L=2$), a combined spike-homogenenous ($\boldsymbol{g}^*=(1,0.5,1,0.6)^T$, $x_0=25$, $\boldsymbol{A}=(0.05,0,0,0)^T$, $L=2$) and a noise-homogeneous ($\boldsymbol{g}^*=(1,0.5,1,0.6)^T$, $A_i=0.05$ for all $i$) initial pattern. Simulations for the spike initial conditions include the clamping introduced in (S6.2). The common choice of model parameters for all simulations corresponds to,

$$\boldsymbol{W}^T=\begin{pmatrix}1 & 0 & 0 & 0\\ 2 & 0 & 0 & 0\\ 3 & 1 & 0 & 0\\ 0 & -5 & 3 & 0\end{pmatrix}; \qquad \text{(S6.5)}$$

and,

$$\begin{aligned} &\sigma_1=1, \quad \sigma_2=0, \quad \sigma_3=0, \quad \sigma_4=0; \\ &\mu_1=0, \quad \mu_2=2, \quad \mu_3=1, \quad \mu_4=1.5; \\ &d_1=0, \quad d_2=8, \quad d_3=35, \quad d_4=0. \end{aligned} \tag{S6.6}$$

##### *Figure 7 in the main text*

Figure 7 in the main text shows the pattern transformations that two 5-node $H^0$ gene networks and one 4-node H gene network produce on a spike initial pattern ($\boldsymbol{g}^*=\boldsymbol{0}$, $x_0=25$, $\boldsymbol{A}=(0.05,0,0,0)^T$, $L=2$). The two $H^0$ networks include non-linear regulations in the first term of (S6.1), that is, one of the entries of function $\boldsymbol{f}$. Also, the clamping introduced in (S6.2) is considered to avoid negative concentrations. The parameter choice for panel A is given by,

$$\boldsymbol{W}^T=\begin{pmatrix} 2 & 0 & 0 & 0 & 0 \\ 3 & 0 & 0 & 0 & 0 \\ 0 & 2 & 0 & 0 & 0 \\ 0 & 1 & 0 & 0 & 0 \\ 0 & 0 & -3 & 1 & 0 \end{pmatrix}, \tag{S6.7}$$

with $f_3(g_2)=W_{23}(g_2-g_2^*)^3-\sigma_3(g_3-g_3^*)^3+\mu_3 g_3^*=2g_2^3$; and,

$$\begin{aligned} &\sigma_1=1, \quad \sigma_2=0, \quad \sigma_3=0, \quad \sigma_4=0, \quad \sigma_5=0; \\ &\mu_1=0, \quad \mu_2=1, \quad \mu_3=0.5, \quad \mu_4=0.5, \quad \mu_5=0.5; \\ &d_1=0, \quad d_2=40, \quad d_3=0, \quad d_4=0, \quad d_5=0. \end{aligned} \tag{S6.8}$$

The parameter choice for panel B is given by,

$$\boldsymbol{W}^T=\begin{pmatrix} 2 & 0 & 0 & 0 & 0 \\ 3 & 0 & 0 & 0 & 0 \\ 0 & 2 & 0 & 0 & 0 \\ 0 & 1 & 0 & 0 & 0 \\ 0 & 0 & -1 & 2 & 0 \end{pmatrix}, \tag{S6.9}$$

with $f_5(g_3,g_4)=W_{45}(g_4-g_4^*)+W_{35}(g_3-g_3^*)^3-\sigma_5(g_5-g_5^*)^3+\mu_5 g_5^*=2g_4-g_3^3$; and,

$$\begin{aligned} &\sigma_1=1, \quad \sigma_2=0, \quad \sigma_3=0, \quad \sigma_4=0, \quad \sigma_5=0; \\ &\mu_1=0, \quad \mu_2=1, \quad \mu_3=0.5, \quad \mu_4=0.5, \quad \mu_5=0.5; \\ &d_1=0, \quad d_2=40, \quad d_3=0, \quad d_4=0, \quad d_5=0. \end{aligned} \tag{S6.10}$$

The parameter choice for panel C is given by,

$$\boldsymbol{W}^T=\begin{pmatrix} 2 & 0 & 0 & 0 \\ 1 & 0 & 0 & 0 \\ 1 & 0 & 0 & 0 \\ 0 & 1 & -2 & 0 \end{pmatrix}; \tag{S6.11}$$

and,

$$\begin{aligned}&\sigma_1=1, \quad \sigma_2=0, \quad \sigma_3=0, \quad \sigma_4=0;\\&\mu_1=0, \quad \mu_2=1, \quad \mu_3=0.5, \quad \mu_4=0.5;\\&d_1=0, \quad d_2=40, \quad d_3=0, \quad d_4=0.\end{aligned} \tag{S6.12}$$

*Figures 8 and 9 in the main text*

Figures 8 and 9 in the main text show the 1D and 2D pattern transformations that a 4-node hierarchic gene network, a 3-node over-Turing gene network and a 3-node Turing gene network produce on a spike ($x_0=25$ and $\boldsymbol{x_0}=(25,25)^T$, $\boldsymbol{A}=(0.05,0,0,0)^T$, $L_{1D}=2$ and $L_{2D}=3$), a spike-homogeneous ($x_0=25$ and $\boldsymbol{x_0}=(25,25)^T$, $\boldsymbol{A}=(0.05,0,0,0)^T$, $L_{1D}=2$ and $L_{2D}=3$), and a noise-homogeneous ($A_i=0.05$ for all $i$) initial pattern. Also, the clamping introduced in (S6.2) is considered in the case of spike initial patterns.

The choice of model parameters is different for each of the three types of gene regulatory networks, but identical regardless the initial pattern being considered. Networks of the same kind use the same set of model parameters in both figures. For the hierarchical network the parameter choice is given by,

$$\boldsymbol{W}^T=\begin{pmatrix}2 & 0 & 0 & 0\\4 & 0 & 0 & 0\\4 & 0 & 0 & 0\\0 & 4 & -1 & 0\end{pmatrix}; \tag{S6.13}$$

and,

$$\begin{aligned}&\sigma_I=1, \quad \sigma_2=0, \quad \sigma_3=0, \quad \sigma_P=0;\\&\mu_I=0, \quad \mu_2=1, \quad \mu_3=0.5, \quad \mu_P=2;\\&d_I=0, \quad d_2=40, \quad d_3=5, \quad d_P=0;\end{aligned} \tag{S6.14}$$

with $\boldsymbol{g}^*=\boldsymbol{0}$ for the spike initial pattern, and $\boldsymbol{g}^*=(1,1,2,0.5)^T$ for the spike-homogeneous and noise-homogeneous initial patterns.

For the over-Turing network, the parameter choice is given by,

$$\boldsymbol{W}^T=\begin{pmatrix}0 & 0.5 & -1\\1 & 0 & 0\\1 & 0 & 0\end{pmatrix}; \tag{S6.15}$$

and,

$$\begin{aligned}&\sigma_I=1, \quad \sigma_2=1, \quad \sigma_P=0;\\&\mu_I=0, \quad \mu_2=0, \quad \mu_P=1;\\&d_I=0, \quad d_2=0, \quad d_P=1;\end{aligned} \tag{S6.16}$$

with $\boldsymbol{g}^{*}=\mathbf{0}$ for the spike initial pattern, and $\boldsymbol{g}^{*}=(1,1,1)^{T}$ for the spike-homogeneous and noise-homogeneous initial patterns.

Finally, for the Turing network, the choice of model parameters is given by,

$$\boldsymbol{W}^{T}=\begin{pmatrix} 0 & 0.125 & -1 \\ 1 & -0.5 & 0 \\ 0.09375 & 0 & 0 \end{pmatrix}; \qquad \text{(S6.17)}$$

and,

$$\begin{array}{lll} \sigma_{I}=1, & \sigma_{P}=1, & \sigma_{3}=0; \\ \mu_{I}=0, & \mu_{P}=0, & \mu_{3}=0.25; \\ d_{I}=0, & d_{P}=2, & d_{3}=2; \end{array} \qquad \text{(S6.18)}$$

with $\boldsymbol{g}^{*}=\mathbf{0}$ for the spike initial pattern, and $\boldsymbol{g}^{*}=(1,1,4)^{T}$ for the spike-homogeneous and noise-homogeneous initial patterns.

##### *Figure 10 in the main text*

Figure 10 in the main text shows the pattern transformations that two 6-node hierarchic gene regulatory networks produce on a spike initial pattern ($\boldsymbol{g}^{*}=\mathbf{0}$, $x_0=25$, $\boldsymbol{A}=(0.05,0,0,0)^{T}$, $L=2$). The clamping introduced in (S6.2) is considered to prevent negative concentrations. The parameter choice for panel A is given by,

$$\boldsymbol{W}^{T}=\begin{pmatrix} 1 & 0 & 0 & 0 & 0 & 0 \\ 45 & 0 & 0 & 0 & 0 & 0 \\ 40 & 0 & 0 & 0 & 0 & 0 \\ 0 & 60 & -65.5 & 0 & 0 & 0 \\ 9 & 0 & 0 & 0 & 0 & 0 \\ 0 & 0 & 0 & -6 & 2 & 0 \end{pmatrix}; \qquad \text{(S6.19)}$$

and,

$$\begin{array}{llllll} \sigma_{1}=1, & \sigma_{2}=0, & \sigma_{3}=0, & \sigma_{4}=0, & \sigma_{5}=0, & \sigma_{6}=0; \\ \mu_{1}=0, & \mu_{2}=1, & \mu_{3}=1, & \mu_{4}=1, & \mu_{5}=1, & \mu_{6}=1; \\ d_{1}=0, & d_{2}=45, & d_{3}=40, & d_{4}=0, & d_{5}=1, & d_{6}=0. \end{array} \qquad \text{(S6.20)}$$

Similarly, the parameter choice for panel B is given by,

$$\boldsymbol{W}^{T}=\begin{pmatrix} 1 & 0 & 0 & 0 & 0 & 0 \\ 45 & 0 & 0 & 0 & 0 & 0 \\ 40 & 0 & 0 & 0 & 0 & 0 \\ 0 & 5 & -4.7 & 0 & 0 & 0 \\ 100 & 0 & 0 & 0 & 0 & 0 \\ 0 & 0 & 0 & -1.25 & 1.5 & 0 \end{pmatrix}; \qquad \text{(S6.21)}$$

and,

$$\begin{array}{llllll} \sigma_1=1, & \sigma_2=0, & \sigma_3=0, & \sigma_4=0, & \sigma_5=0, & \sigma_6=0; \\ \mu_1=0, & \mu_2=1, & \mu_3=1, & \mu_4=1, & \mu_5=1, & \mu_6=1; \\ d_1=0, & d_2=2, & d_3=1, & d_4=0, & d_5=50, & d_6=0. \end{array} \quad \text{(S6.22)}$$

*Figure 11 in the main text*

Figure 11 in the main text shows the pattern transformations that a 5-node hierarchic gene network produces on a spike initial pattern ($\boldsymbol{g}^*=\mathbf{0}$, $x_0=25$, $\boldsymbol{A}=(0.05,0,0,0)^T$, $L=2$) for several sets of model parameters. The clamping introduced in (S6.2) is considered to prevent negative concentrations. The parameter choices for panel C correspond to,

$$\boldsymbol{W}^T=\begin{pmatrix} 2 & 0 & 0 & 0 & 0 \\ 1 & 0 & 0 & 0 & 0 \\ 1 & 0 & 0 & 0 & 0 \\ 0 & 3 & -2 & 0 & 0 \\ 0 & 0 & 0 & W_{45} & 0 \end{pmatrix}, \quad \text{(S6.23)}$$

with $W_{45}\in\{0.5\ \text{(blue)}, 1\ \text{(green)}, 2\ \text{(red)}\}$; and,

$$\begin{array}{lllll} \sigma_1=1, & \sigma_2=0, & \sigma_3=0, & \sigma_4=0, & \sigma_5=0; \\ \mu_1=0, & \mu_2=1, & \mu_3=0.5, & \mu_4=0.5, & \mu_5=1; \\ d_1=0, & d_2=40, & d_3=2, & d_4=0, & d_5=0. \end{array} \quad \text{(S6.24)}$$

Similarly, the parameter choices for panel D are given by,

$$\boldsymbol{W}^T=\begin{pmatrix} 2 & 0 & 0 & 0 & 0 \\ 1 & 0 & 0 & 0 & 0 \\ 1 & 0 & 0 & 0 & 0 \\ 0 & 3 & -2 & 0 & 0 \\ 0 & 0 & 0 & W_{45} & 0 \end{pmatrix}, \quad \text{(S6.25)}$$

with $W_{45}\in\{0.94984\ \text{(blue)}, 1.79324\ \text{(green)}, 3.03495\ \text{(red)}\}$; and,

$$\begin{array}{lllll} \sigma_1=1, & \sigma_2=0, & \sigma_3=0, & \sigma_4=0, & \sigma_5=0; \\ \mu_1=0, & \mu_2=1, & \mu_3=0.5, & \mu_4=0.5, & \mu_5=1; \\ d_1=0, & d_2=40, & d_3=d, & d_4=0, & d_5=0; \end{array} \quad \text{(S6.26)}$$

with $d\in\{0.5\ \text{(blue)}, 2\ \text{(green)}, 4\ \text{(red)}\}$.

*Figure 12 in the main text*

Figure 12 in the main text shows the pattern transformations that the combinations of hierarchic, over-Turing and Turing gene regulatory subnetworks produce on a spike-homogeneous initial pattern ($x_0=25$, $A_I=0.05$ and $A_i=0$ for any other $i$, $L=2$). The choice of model parameters for (S6.1) is different for each of the combinations. Also, some of the simulations include non-linear regulations in the first term of system (S6.1), that is, one of the entries of function $\boldsymbol{f}$.

The top-left panel shows the pattern transformation produced by an 8-node hierarchic + hierarchic gene network. The parameter choice in this case is given by,

$$\boldsymbol{W}^T=\begin{pmatrix} 2 & 0 & 0 & 0 & 0 & 0 & 0 & 0 \\ 3 & 0 & 0 & 0 & 0 & 0 & 0 & 0 \\ 0 & 2 & 0 & 0 & 0 & 0 & 0 & 0 \\ 0 & 1 & 0 & 0 & 0 & 0 & 0 & 0 \\ 0 & 0 & -1 & 2 & 0 & 0 & 0 & 0 \\ 0 & 0 & 0 & 0 & 1 & 0 & 0 & 0 \\ 0 & 0 & 0 & 0 & 1 & 0 & 0 & 0 \\ 0 & 0 & 0 & 0 & 0 & -1.2 & 1 & 0 \end{pmatrix}, \qquad \text{(S6.27)}$$

with $f_5=W_{45}(g_4-g_4^*)-W_{35}(g_3-g_3^*)^3-\sigma_5(g_5-g^*)^3+\mu_5 g_5^*=2(g_4-2)-(g_3-2)^3-0.5(g_5-2)^3$ and $f_P=W_{6P}(g_6-g_6^*)-W_{7P}(g_7-g_7^*)^3-\sigma_P(g_P-g_P^*)^3+\mu_P g_P^*=(g_6-1)-1.2(g_7-1)^3-0.5(g_P-2)^3$; $\boldsymbol{g}^*=(1,1,2,2,2,1,1,2)^T$; and,

$$\begin{array}{llllllll} \sigma_I=1, & \sigma_2=0, & \sigma_3=0, & \sigma_4=0, & \sigma_5=0.5, & \sigma_6=0, & \sigma_7=0, & \sigma_P=0.5; \\ \mu_I=0, & \mu_2=1, & \mu_3=0.5, & \mu_4=0.5, & \mu_5=0, & \mu_6=1, & \mu_7=1, & \mu_P=0; \\ d_I=0, & d_2=250, & d_3=0, & d_4=0, & d_5=0, & d_6=0, & d_7=0, & d_P=0. \end{array} \qquad \text{(S6.28)}$$

The top-middle panel shows the pattern transformation produced by an 8-node hierarchic + over-Turing gene network. The parameter choice in this case is given by,

$$\boldsymbol{W}^T=\begin{pmatrix} 2 & 0 & 0 & 0 & 0 & 0 & 0 & 0 \\ 3 & 0 & 0 & 0 & 0 & 0 & 0 & 0 \\ 0 & 2 & 0 & 0 & 0 & 0 & 0 & 0 \\ 0 & 1 & 0 & 0 & 0 & 0 & 0 & 0 \\ 0 & 0 & -1 & 2 & 0 & 0 & 0 & 0 \\ 0 & 0 & 0 & 0 & 0.5 & 0 & 0.5 & -1 \\ 0 & 0 & 0 & 0 & 0 & 1 & 0 & 0 \\ 0 & 0 & 0 & 0 & 0 & 1 & 0 & 0 \end{pmatrix}, \qquad \text{(S6.28)}$$

with $f_5=W_{45}(g_4-g_4^*)-W_{35}(g_3-g_3^*)^3-\sigma_5(g_5-g_5^*)^3+\mu_5 g_5^*=2(g_4-2)-(g_3-2)^3-0.5(g_5-2)^3$; $\boldsymbol{g}^*=(1,1,2,2,2,1,1,1)^T$; and,

$$\begin{array}{llllllll} \sigma_I=1, & \sigma_2=0, & \sigma_3=0, & \sigma_4=0, & \sigma_5=0.5, & \sigma_6=1, & \sigma_7=1, & \sigma_P=0; \\ \mu_I=0, & \mu_2=1, & \mu_3=0.5, & \mu_4=0.5, & \mu_5=0, & \mu_6=0, & \mu_7=0, & \mu_P=1; \\ d_I=0, & d_2=4, & d_3=0, & d_4=0, & d_5=0, & d_6=0, & d_7=0, & d_P=5. \end{array} \qquad \text{(S6.29)}$$

The top-right panel shows the pattern transformation produced by an 8-node hierarchic + Turing gene network. The parameter choice in this case is given by,

$$
\boldsymbol{W}^T = \begin{pmatrix} 2 & 0 & 0 & 0 & 0 & 0 & 0 & 0 \\ 3 & 0 & 0 & 0 & 0 & 0 & 0 & 0 \\ 0 & 2 & 0 & 0 & 0 & 0 & 0 & 0 \\ 0 & 1 & 0 & 0 & 0 & 0 & 0 & 0 \\ 0 & 0 & -1 & 2 & 0 & 0 & 0 & 0 \\ 0 & 0 & 0 & 0 & 0.01 & 0 & 0.1255 & -1 \\ 0 & 0 & 0 & 0 & 0 & 1 & -0.5 & 0 \\ 0 & 0 & 0 & 0 & 0 & 0.09375 & 0 & 0 \end{pmatrix}, \tag{S6.30}
$$

with $f_5 = W_{45}(g_4 - g_*) - W_{35}(g_3 - g_3^*)^3 - \sigma_5(g_5 - g_5^*)^3 + \mu_5 g_5^* = 2(g_4 - 2) - (g_3 - 2)^3 - 2(g_5 - 0.5)^3$;
$\boldsymbol{g}^* = (1,1,2,2,0.5,1,1,4)^T$; and,

$$
\begin{array}{llllllll}
\sigma_I=1, & \sigma_2=0, & \sigma_3=0, & \sigma_4=0, & \sigma_5=2, & \sigma_6=1, & \sigma_P=1, & \sigma_8=0; \\
\mu_I=0, & \mu_2=1, & \mu_3=0.5, & \mu_4=0.5, & \mu_5=0, & \mu_6=0, & \mu_P=0, & \mu_8=0.25; \\
d_I=0, & d_2=4, & d_3=0, & d_4=0, & d_5=0, & d_6=0, & d_P=0.4, & d_8=0.4.
\end{array} \tag{S6.31}
$$

The middle-left panel shows the pattern transformation produced by a 6-node over-Turing + hierarchic gene network. The parameter choice in this case is given by,

$$
\boldsymbol{J} = \begin{pmatrix} 0 & 0.5 & -1 & 0 & 0 & 0 \\ 1 & 0 & 0 & 0 & 0 & 0 \\ 1 & 0 & 0 & 0 & 0 & 0 \\ 0 & 0 & 1 & 0 & 0 & 0 \\ 0 & 0 & 1 & 0 & 0 & 0 \\ 0 & 0 & 0 & 1 & -4 & 0 \end{pmatrix}, \tag{S6.32}
$$

with $f_P = W_{4P}(g_4 - g_4^*) - W_{5P}(g_5 - g_5^*)^3 - \sigma_P(g_P - g_P^*)^3 + \mu_P g_P^* = (g_4 - 2) - 4(g_5 - 2)^3 + 1$;
$\boldsymbol{g}^* = (1,0.6,1,2,2,1)^T$; and,

$$
\begin{array}{llllll}
\sigma_I=1, & \sigma_2=1.5, & \sigma_3=0, & \sigma_4=0, & \sigma_5=0, & \sigma_P=0; \\
\mu_I=0, & \mu_2=0, & \mu_3=1, & \mu_4=0.5, & \mu_5=0.5, & \mu_P=1; \\
d_I=0, & d_2=0, & d_3=5, & d_4=0, & d_5=0, & d_P=0.
\end{array} \tag{S6.33}
$$

The central panel shows the pattern transformation produced by a 6-node over-Turing + over-Turing gene network. The parameter choice in this case is given by,

$$
\boldsymbol{W}^T = \begin{pmatrix} 0 & 0.5 & -1 & 0 & 0 & 0 \\ 1 & 0 & 0 & 0 & 0 & 0 \\ 1 & 0 & 0 & 0 & 0 & 0 \\ 0 & 0 & 1 & 0 & 0.5 & -1 \\ 0 & 0 & 0 & 1 & 0 & 0 \\ 0 & 0 & 0 & 1 & 0 & 0 \end{pmatrix}; \tag{S6.34}
$$

$\boldsymbol{g}^* = (1,0.6,1,2,2,1)^T$; and,

$$\begin{array}{llllll} \sigma_I=1, & \sigma_2=1.5, & \sigma_3=0, & \sigma_4=0.5, & \sigma_5=0.5, & \sigma_P=0; \\ \mu_I=0, & \mu_2=0, & \mu_3=1, & \mu_4=0, & \mu_5=0, & \mu_P=1; \\ d_I=0, & d_2=0, & d_3=5, & d_4=0, & d_5=0, & d_P=10. \end{array} \tag{S6.35}$$

Panel B3 shows the pattern transformation produced by a 6-node over-Turing + Turing gene network. The parameter choice in this case is given by,

$$\boldsymbol{W}^T=\begin{pmatrix} 0 & 0.5 & -1 & 0 & 0 & 0 \\ 1 & 0 & 0 & 0 & 0 & 0 \\ 1 & 0 & 0 & 0 & 0 & 0 \\ 0 & 0 & 0.5 & 0 & 0.125 & -1 \\ 0 & 0 & 0 & 1 & -0.5 & 0 \\ 0 & 0 & 0 & 0.09375 & 0 & 0 \end{pmatrix}; \tag{S6.36}$$

$\boldsymbol{g}^*=(1,0.6,1,1,1,4)^T$; and,

$$\begin{array}{llllll} \sigma_I=1, & \sigma_2=1.5, & \sigma_3=0, & \sigma_4=1, & \sigma_P=1, & \sigma_6=0; \\ \mu_I=0, & \mu_2=0, & \mu_3=1, & \mu_4=0, & \mu_P=0, & \mu_6=0.25; \\ d_I=0, & d_2=0, & d_3=5, & d_4=0, & d_P=1, & d_6=1. \end{array} \tag{S6.37}$$

The bottom-left panel shows the pattern transformation produced by a 6-node Turing + hierarchic gene network. The parameter choice in this case is given by,

$$\boldsymbol{W}^T=\begin{pmatrix} 0 & 0.125 & -1 & 0 & 0 & 0 \\ 1 & -0.5 & 0 & 0 & 0 & 0 \\ 0.09375 & 0 & 0 & 0 & 0 & 0 \\ 0 & 1.5 & 0 & 0 & 0 & 0 \\ 0 & 1.5 & 0 & 0 & 0 & 0 \\ 0 & 0 & 0 & 1 & -16 & 0 \end{pmatrix}, \tag{S6.38}$$

with $f_P=W_{4P}(g_4-g_4^*)-W_{5P}(g_5-g_5^*)^3-\sigma_P(g_P-g_P^*)^3+\mu_P g_P^*=(g_4-1)-16(g_5-2)^3+1$; $\boldsymbol{g}^*=(1,1,4,1,2,1)^T$; and,

$$\begin{array}{llllll} \sigma_I=1, & \sigma_2=1, & \sigma_3=0, & \sigma_4=0, & \sigma_5=0, & \sigma_P=0; \\ \mu_I=0, & \mu_2=0, & \mu_3=0.25, & \mu_4=1, & \mu_5=0.5, & \mu_P=1; \\ d_I=0, & d_2=5, & d_3=5, & d_4=0, & d_5=0, & d_P=0. \end{array} \tag{S6.39}$$

The bottom-middle panel shows the pattern transformation produced by a 6-node Turing + over-Turing gene network. The parameter choice in this case is given by,

$$\boldsymbol{W}^T=\begin{pmatrix} 0 & 0.125 & -1 & 0 & 0 & 0 \\ 1 & -0.5 & 0 & 0 & 0 & 0 \\ 0.09375 & 0 & 0 & 0 & 0 & 0 \\ 0 & 0.5 & 0 & 0 & 0.5 & -1 \\ 0 & 0 & 0 & 1 & 0 & 0 \\ 0 & 0 & 0 & 1 & 0 & 0 \end{pmatrix}; \tag{S6.40}$$

$g^*=(1,1,4,2,2,1)^T$; and,

$$\begin{aligned}
&\sigma_I=1, \quad \sigma_2=1, \quad \sigma_3=0, \quad \sigma_4=0.5, \quad \sigma_5=0.5, \quad \sigma_P=0;\\
&\mu_I=0, \quad \mu_2=0, \quad \mu_3=0.25, \quad \mu_4=0, \quad \mu_5=0, \quad \mu_P=1;\\
&d_I=0, \quad d_2=1, \quad d_3=1, \quad d_4=0, \quad d_5=0, \quad d_P=2.
\end{aligned} \tag{S6.41}$$

Finally, the bottom-right panel shows the pattern transformation produced by a 6-node Turing + Turing gene network. The parameter choice in this case is given by,

$$W^T=\begin{pmatrix} 0 & 0.125 & -1 & 0 & 0 & 0\\ 1 & -0.5 & 0 & 0 & 0 & 0\\ 0.09375 & 0 & 0 & 0 & 0 & 0\\ 0 & 0.05 & 0 & 0 & 0.125 & -1\\ 0 & 0 & 0 & 1 & -0.5 & 0\\ 0 & 0 & 0 & 0.09375 & 0 & 0 \end{pmatrix}; \tag{S6.42}$$

$g^*=(1,1,4,1,1,4)^T$; and,

$$\begin{aligned}
&\sigma_I=1, \quad \sigma_2=1, \quad \sigma_3=0, \quad \sigma_4=1, \quad \sigma_P=1, \quad \sigma_6=0;\\
&\mu_I=0, \quad \mu_2=0, \quad \mu_3=0.25, \quad \mu_4=0, \quad \mu_P=0, \quad \mu_6=0.25;\\
&d_I=0, \quad d_2=2, \quad d_3=2, \quad d_4=0, \quad d_P=0.5, \quad d_6=0.5.
\end{aligned} \tag{S6.43}$$

*Figure S2 in the supporting information*

Figure S2 in the supporting information shows the pattern transformations that an 8-node hierarchic gene network produces on a spike initial pattern ($g^*=\mathbf{0}$, $x_0=25$, $A=(0.05,0,0,0)^T$, $L=2$) for several sets of model parameters. The clamping introduced in (S6.2) is considered to prevent negative concentrations. The parameter choices for panel C correspond to,

$$W^T=\begin{pmatrix} 1 & 0 & 0 & 0 & 0 & 0 & 0 & 0\\ 45 & 0 & 0 & 0 & 0 & 0 & 0 & 0\\ 40 & 0 & 0 & 0 & 0 & 0 & 0 & 0\\ 0 & 5 & -4.7 & 0 & 0 & 0 & 0 & 0\\ 100 & 0 & 0 & 0 & 0 & 0 & 0 & 0\\ 0 & 0 & 0 & -1.25 & 1.5 & 0 & 0 & 0\\ 0 & 0 & 0 & 0 & 0 & 2 & 0 & 0\\ 0 & 0 & 1.5 & W_{48} & 0 & -3 & 2 & 0 \end{pmatrix}, \tag{S6.44}$$

with $W_{48}\in\{-1.8 \text{ (blue)}, -0.3 \text{ (red)}\}$; and,

$$\begin{aligned}
&\sigma_1=1, \quad \sigma_2=0, \quad \sigma_3=0, \quad \sigma_4=0, \quad \sigma_5=0, \quad \sigma_6=0, \quad \sigma_7=0, \quad \sigma_8=0;\\
&\mu_1=0, \quad \mu_2=1, \quad \mu_3=1, \quad \mu_4=1, \quad \mu_5=1, \quad \mu_6=1, \quad \mu_7=1, \quad \mu_8=1;\\
&d_1=0, \quad d_2=2, \quad d_3=1, \quad d_4=0, \quad d_5=40, \quad d_6=0, \quad d_7=20, \quad d_8=0.
\end{aligned} \tag{S6.45}$$

Similarly, the parameter choices for panel D are given by,

$$W^T = \begin{pmatrix} 1 & 0 & 0 & 0 & 0 & 0 & 0 & 0 \\ 45 & 0 & 0 & 0 & 0 & 0 & 0 & 0 \\ 40 & 0 & 0 & 0 & 0 & 0 & 0 & 0 \\ 0 & 5 & W_{34} & 0 & 0 & 0 & 0 & 0 \\ 100 & 0 & 0 & 0 & 0 & 0 & 0 & 0 \\ 0 & 0 & 0 & -1.25 & 1.5 & 0 & 0 & 0 \\ 0 & 0 & 0 & 0 & 0 & 2 & 0 & 0 \\ 0 & 0 & 1.5 & W_{48} & 0 & -3 & 2 & 0 \end{pmatrix}, \tag{S6.46}$$

with $W_{34} \in \{-4.7 \text{ (blue)}, -4.9 \text{ (green)}\}$ and $W_{48} \in \{-1.8 \text{ (blue)}, -2.7 \text{ (green)}\}$; and,

$$\begin{aligned} &\sigma_1=1, \quad \sigma_2=0, \quad \sigma_3=0, \quad \sigma_4=0, \quad \sigma_5=0, \quad \sigma_6=0, \quad \sigma_7=0, \quad \sigma_8=0; \\ &\mu_1=0, \quad \mu_2=1, \quad \mu_3=1, \quad \mu_4=1, \quad \mu_5=1, \quad \mu_6=1, \quad \mu_7=1, \quad \mu_8=1; \\ &d_1=0, \quad d_2=2, \quad d_3=1, \quad d_4=0, \quad d_5=40, \quad d_6=0, \quad d_7=20, \quad d_8=0. \end{aligned} \tag{S6.47}$$

##### *Figure S3 in the supporting information*

Figure S3 in the supporting information shows the pattern transformations that a 2-node over-Truing network produces on a noise-homogeneous initial pattern ($\boldsymbol{g}^* = (1.25, 0.6)^T$, $A_i = 0.05$ for all $i$) and two spike-homogeneous initial patterns ($\boldsymbol{g}^* = (1.25, 0.6)^T$, $x_0 = 25$ (left), $x_0 = 0$ (right), $\boldsymbol{A} = (0.05, 0, 0, 0)^T$, $L = 2$). The parameter choices for all panels in the same and it is given by,

$$W^T = \begin{pmatrix} 1 & -1 \\ 2 & 0 \end{pmatrix}, \tag{S6.48}$$

and,

$$\begin{aligned} &\sigma_1=0.8, \quad \sigma_2=0; \\ &\mu_1=0, \quad \mu_2=1.5; \\ &d_1=0, \quad d_2=8. \end{aligned} \tag{S6.49}$$

##### *Figure S4 in the supporting information*

Figure S4 in the supporting information show the pattern transformations that a 3-node over-Turing gene network and a 3-node Turing gene network produce on a spike-homogeneous initial pattern ($x_0 = 25$ (left), $x_0 = 0$ (right), $\boldsymbol{A} = (0.05, 0, 0)^T$, $L = 2$). In panel D, the basal concentration is $\boldsymbol{g}^* = (1, 0.6, 1)^T$, and the choice of model parameters is given by,

$$W^T = \begin{pmatrix} 0 & 0.5 & -1 \\ 1 & 0 & 0 \\ 1 & 0 & 0 \end{pmatrix}, \tag{S6.50}$$

and,

$$\begin{aligned} &\sigma_1=1, \quad \sigma_2=1.5 \quad \sigma_3=0, \\ &\mu_1=0, \quad \mu_2=0, \quad \mu_3=1, \\ &d_1=0, \quad d_2=0, \quad d_3=d, \end{aligned} \tag{S6.51}$$

with $d \in \{0.5 \text{ (blue)}, 1 \text{ (green)}, 5 \text{ (red)}\}$.

In panel E, the basal concentration is $\boldsymbol{g}^* = (1,1,4)^T$, and the choice of model parameters is given by,

$$\boldsymbol{W}^T = \begin{pmatrix} 0 & 0.125 & -1 \\ 1 & -0.5 & 0 \\ 0.09375 & 0 & 0 \end{pmatrix}, \tag{S6.52}$$

and,

$$\begin{aligned} &\sigma_1=1, \quad \sigma_2=1, \quad \sigma_3=0, \\ &\mu_1=0, \quad \mu_2=0, \quad \mu_3=0.25, \\ &d_1=0, \quad d_2=d, \quad d_3=d, \end{aligned} \tag{S6.53}$$

with $d \in \{0.5 \text{ (blue)}, 1 \text{ (green)}, 2 \text{ (red)}\}$.

##### *Figure S5 in the supporting information*

Figure S5 in the supporting information shows the pattern transformations that a 2-node $L^+$ gene network, a 4-node $l^+L^-$ gene network and a 4-node $L^+L^-$ gene network produce on a spike ( $x_0=25$, $\boldsymbol{A}=(0.05,0,0,0)^T$, $L=2$), a spike-homogeneous ($x_0=25$, $\boldsymbol{A}=(0.05,0,0,0)^T$, $L=2$), and a noise-homogeneous ($A_i=0.05$ for all $i$) initial pattern. The clamping introduced in (S6.2) is considered in the case of spike initial patterns.

The choice of model parameters is different for each of the three types of gene regulatory networks, but identical regardless the initial pattern being considered. For the $L^+$ network the parameter choice is given by,

$$\boldsymbol{W}^T = \begin{pmatrix} 0 & 1 \\ 1 & 0 \end{pmatrix}, \tag{S6.54}$$

and,

$$\begin{aligned} &\sigma_I=1, \quad \sigma_P=0, \\ &\mu_I=0, \quad \mu_P=2, \\ &d_I=0, \quad d_P=10, \end{aligned} \tag{S6.55}$$

with $\boldsymbol{g}^*=\boldsymbol{0}$ for the spike initial pattern, and $\boldsymbol{g}^*=(1,0.5)^T$ for the spike-homogeneous and noise-homogeneous initial patterns.

For the $l^+L^-$ network, the parameter choice is given by,

$$W^T = \begin{pmatrix} 0 & 1 & 0 & 0 \\ 1 & 0 & 0 & 0 \\ 0 & 0.5 & 0 & -4 \\ 0 & 0 & 1 & 0 \end{pmatrix}, \quad \text{(S6.56)}$$

and,

$$\begin{aligned} &\sigma_I = 1, \quad \sigma_2 = 0, \quad \sigma_3 = 0, \quad \sigma_P = 0, \\ &\mu_I = 0, \quad \mu_2 = 2, \quad \mu_3 = 4, \quad \mu_P = 2, \\ &d_I = 0, \quad d_2 = 0, \quad d_3 = 0, \quad d_P = 10, \end{aligned} \quad \text{(S6.57)}$$

with $g^* = \mathbf{0}$ for the spike initial pattern, and $g^* = (1, 0.5, 0.25, 0.5)^T$ for the spike-homogeneous and noise-homogeneous initial patterns.

For the L$^+$L$^-$ network, the parameter choice is give by,

$$W^T = \begin{pmatrix} 0 & 1 & 0 & 0 \\ 1 & 0 & 0 & 0 \\ 0 & 0.5 & 0 & -4 \\ 0 & 0 & 1 & 0 \end{pmatrix}, \quad \text{(S6.58)}$$

and,

$$\begin{aligned} &\sigma_I = 1, \quad \sigma_2 = 0, \quad \sigma_3 = 0, \quad \sigma_P = 0, \\ &\mu_I = 0, \quad \mu_2 = 2, \quad \mu_3 = 4, \quad \mu_P = 2, \\ &d_I = 0, \quad d_2 = 10, \quad d_3 = 0, \quad d_P = 10, \end{aligned} \quad \text{(S6.59)}$$

with $g^* = \mathbf{0}$ for the spike initial pattern, and $g^* = (1, 0.5, 0.25, 0.5)^T$ for the spike-homogeneous and noise-homogeneous initial patterns.

*Figure S6 in the supporting information*

Figure S6 in the supporting information shows the pattern transformations that a 3-node L$^-$ gene network, a 5-node L$^-$l$^+$ gene network and a 5-node L$^-$L+ gene network produce on a spike ($x_0 = 25$, $A = (0.05, 0, 0, 0)^T$, $L = 2$), a spike-homogeneous ($x_0 = 25$, $A = (0.05, 0, 0, 0)^T$, $L = 2$), and a noise-homogeneous ($A_i = 0.05$ for all $i$) initial pattern. The clamping in (S6.2) is considered in the case of spike initial patterns.

The choice of model parameters is different for each of the three types of gene regulatory networks, but identical regardless the initial pattern being considered. For the L$^-$ network the parameter choice is given by,

$$W^T = \begin{pmatrix} 1 & 0 & 0 \\ 0.5 & 0 & -6 \\ 0 & 0.5 & 0 \end{pmatrix}, \quad \text{(S6.60)}$$

and,

$$\begin{aligned}&\sigma_I=1, \quad \sigma_2=0, \quad \sigma_P=0,\\ &\mu_I=0, \quad \mu_2=4, \quad \mu_P=2,\\ &d_I=0, \quad d_2=0, \quad d_P=10,\end{aligned} \tag{S6.61}$$

with $\boldsymbol{g}^*=\mathbf{0}$ for the spike initial pattern, and $\boldsymbol{g}^*=(1,0.25,0.5)^T$ for the spike-homogeneous and noise-homogeneous initial patterns.

For the I⁺L⁻ network, the parameter choice is given by,

$$\boldsymbol{W}^T=\begin{pmatrix}1&0&0&0&0\\0.5&0&-4&0&0\\0&1&0&0&0\\0&0&1&0&1\\0&0&0&1&0\end{pmatrix}, \tag{S6.62}$$

and,

$$\begin{aligned}&\sigma_I=1, \quad \sigma_2=0, \quad \sigma_3=0, \quad \sigma_4=0, \quad \sigma_P=0,\\ &\mu_I=0, \quad \mu_2=2, \quad \mu_3=4, \quad \mu_4=4, \quad \mu_P=2,\\ &d_I=0, \quad d_2=0, \quad d_3=10, \quad d_4=0, \quad d_P=0,\end{aligned} \tag{S6.63}$$

with $\boldsymbol{g}^*=\mathbf{0}$ for the spike initial pattern, and $\boldsymbol{g}^*=(1,0.5,0.25,0.25,0.5)^T$ for the spike-homogeneous and noise-homogeneous initial patterns.

For the L⁺L⁻ network, the parameter choice is give by,

$$\boldsymbol{W}^T=\begin{pmatrix}1&0&0&0&0\\0.5&0&-4&0&0\\0&1&0&0&0\\0&0&1&0&1\\0&0&0&1&0\end{pmatrix}, \tag{S6.64}$$

and,

$$\begin{aligned}&\sigma_I=1, \quad \sigma_2=0, \quad \sigma_3=0, \quad \sigma_4=0, \quad \sigma_P=0,\\ &\mu_I=0, \quad \mu_2=2, \quad \mu_3=4, \quad \mu_4=4, \quad \mu_P=2,\\ &d_I=0, \quad d_2=0, \quad d_3=10, \quad d_4=0, \quad d_P=10,\end{aligned} \tag{S6.65}$$

with $\boldsymbol{g}^*=\mathbf{0}$ for the spike initial pattern, and $\boldsymbol{g}^*=(1,0.5,0.25,0.25,0.5)^T$ for the spike-homogeneous and noise-homogeneous initial patterns.

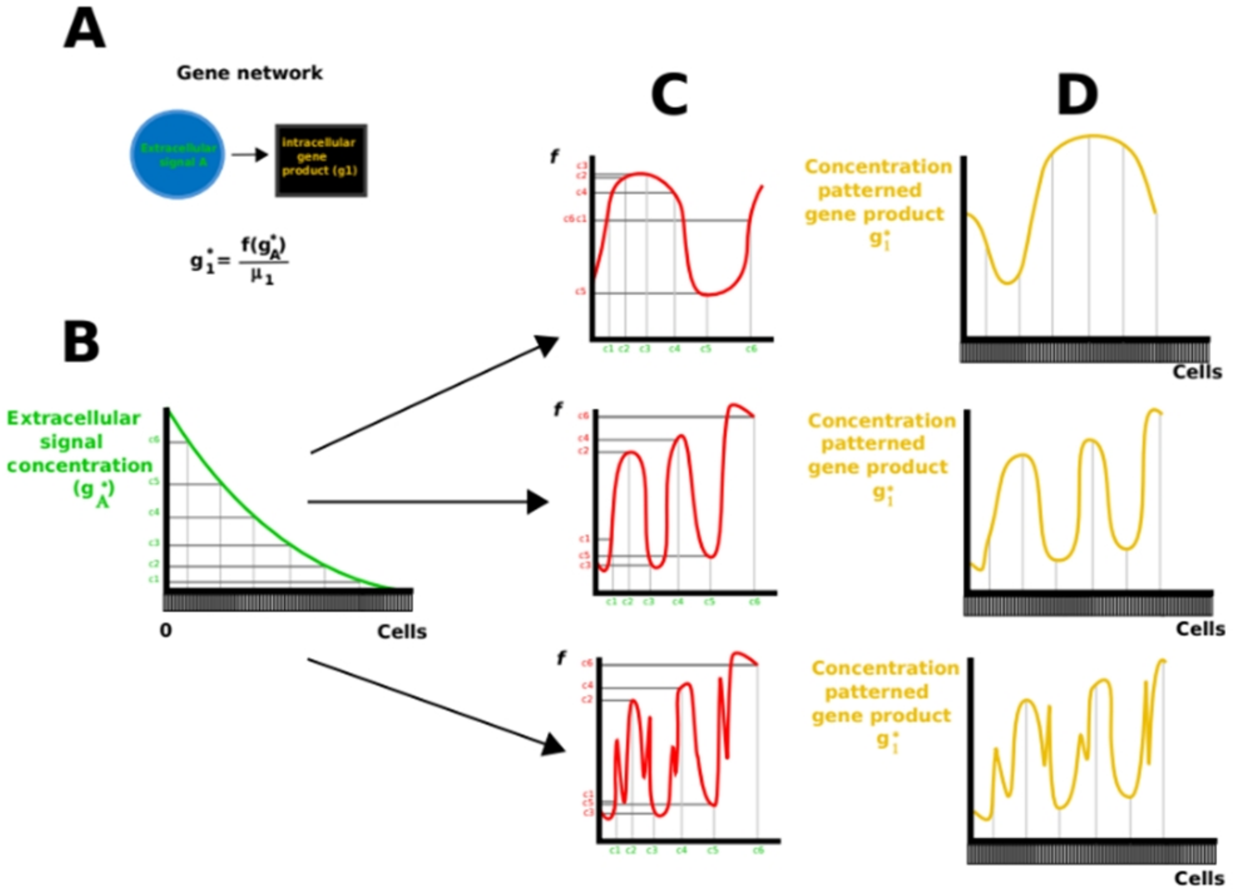


**Figure S1. If function *f* did not satisfy R5, then a two gene network could lead to any resulting pattern in which concentrations are finite. (A)** Gene network topology. An extracellular signal activating an intracellular gene product. $g_A^*$ stands for the concentration of the extracelllular signal in the steady state and $g_1^*$ for that of the intracellular gene product. **(B)** The extracellular signal is secreted from the boundary of an array of cells (x-axis), leading to a gradient in the concentration of the concentration of the signal. Labels c1 to c6 are shown to facilitate the interpretation of the figures but we assume *f* can be understood as continuous. **(C)** Different example functions *f* are plotted. In the x-axis we represent the different concentrations of the signal and in the y-axis the concentration of the intracellular gene product arising from each concentration of the signal. Notice that none of the example functions *f* satisfy R5, i.e. the function is not monotonously growing with the concentration of the signal **(D)** The resulting patterns arising from each *f*. By choosing an adequate *f*, any resulting pattern can be achieved.

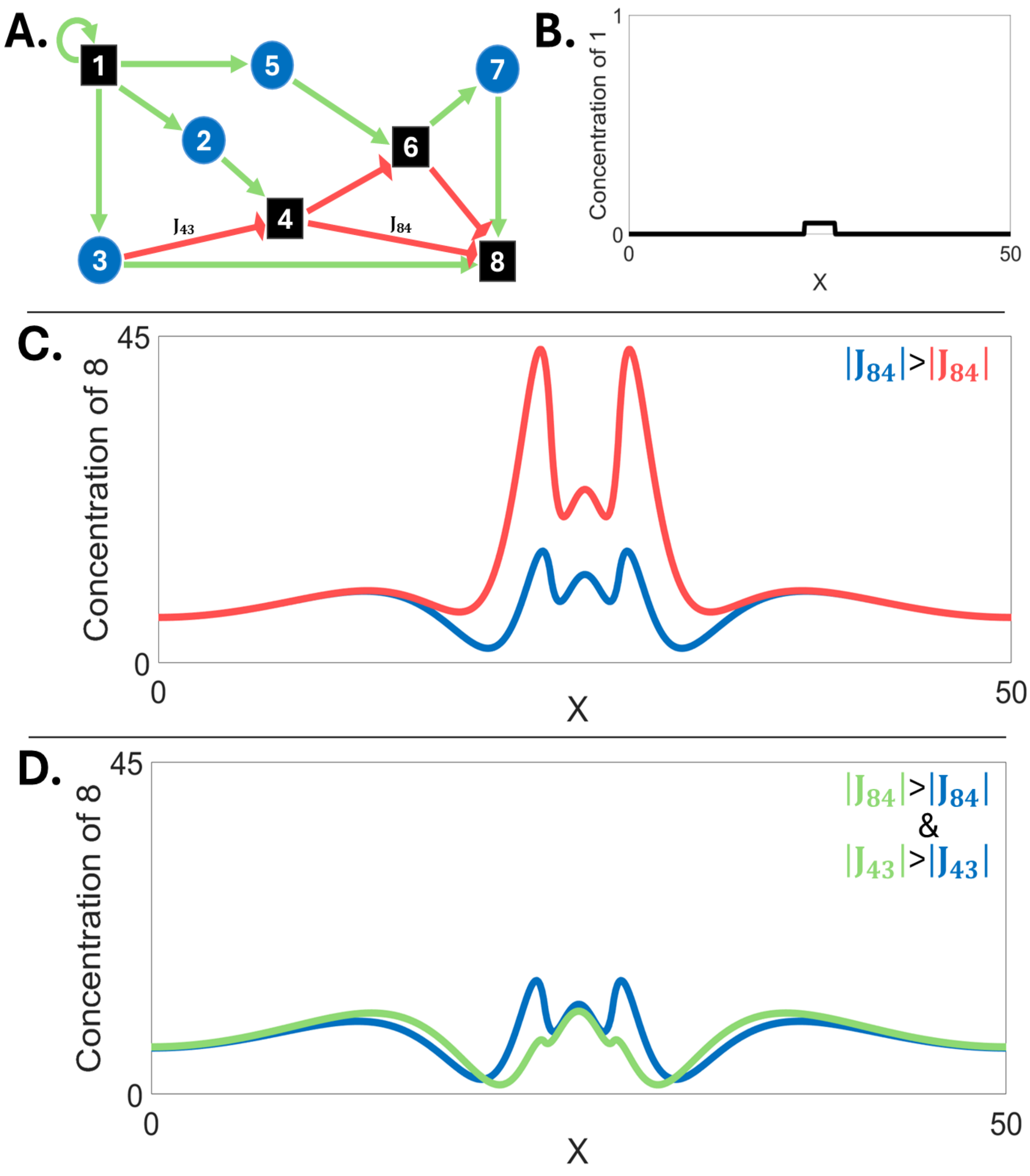


**Figure S2: Example H gene network where by changing the parameters it is possible to vary the height of the different concentration peaks. (A)** Gene network topology. **(B)** Spike initial pattern. **(C-D)** The blue, red and green plots show the concentration of gene product 8 for different combinations of parameter values. As we can see, the height of the concentration peaks of 8 around the initial spike can be independently tuned. Simulations were run using a Forward-Euler algorithm on the Maini-Miura model (see section S6 for parameter values).

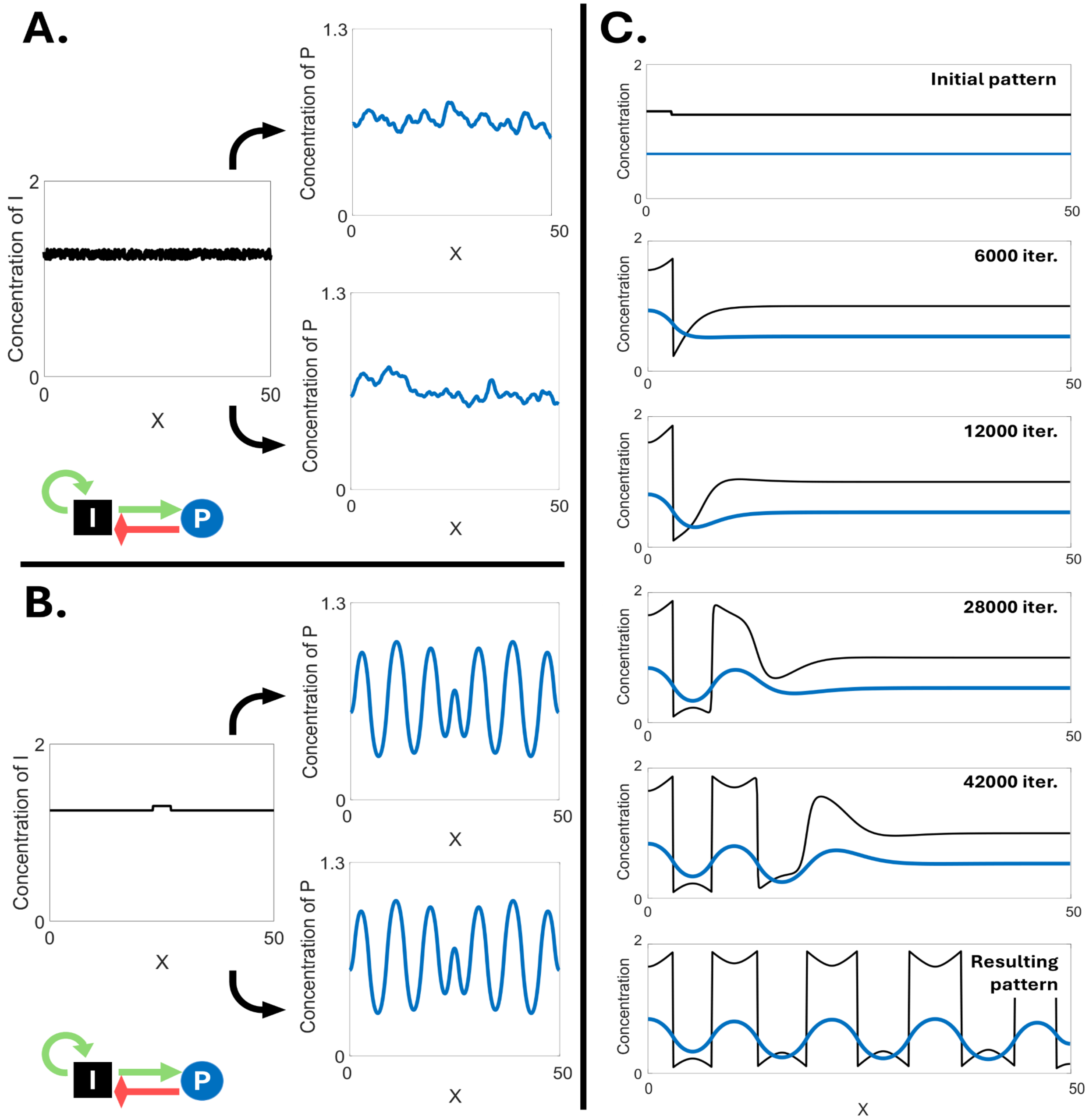


**Figure S3: Pattern formation in over-Turing gene networks. (A)** Noise-homogeneous initial patterns are transformed into amplified noise patterns. Even with the same choices of model parameters, the amplified noise resulting patterns are different every time. **(B)** Spike-homogeneous initial patterns lead to periodical patterns. If the choice of model parameters is the same, the final pattern is the same for each transformation. **(C)** The peaks of concentration formed from a spike-homogeneous initial pattern are produced one by one. The whole process resembles a traveling wave that moves away from the spike and, at the same time, freezes whenever a new peak is formed. Network colors and shapes as in Figure 2. $P$stands for the gene product plotted as resulting pattern while $I$ stands for the gene product in the initial pattern. Simulations as in Figure S2 (see section S6 for parameter values).

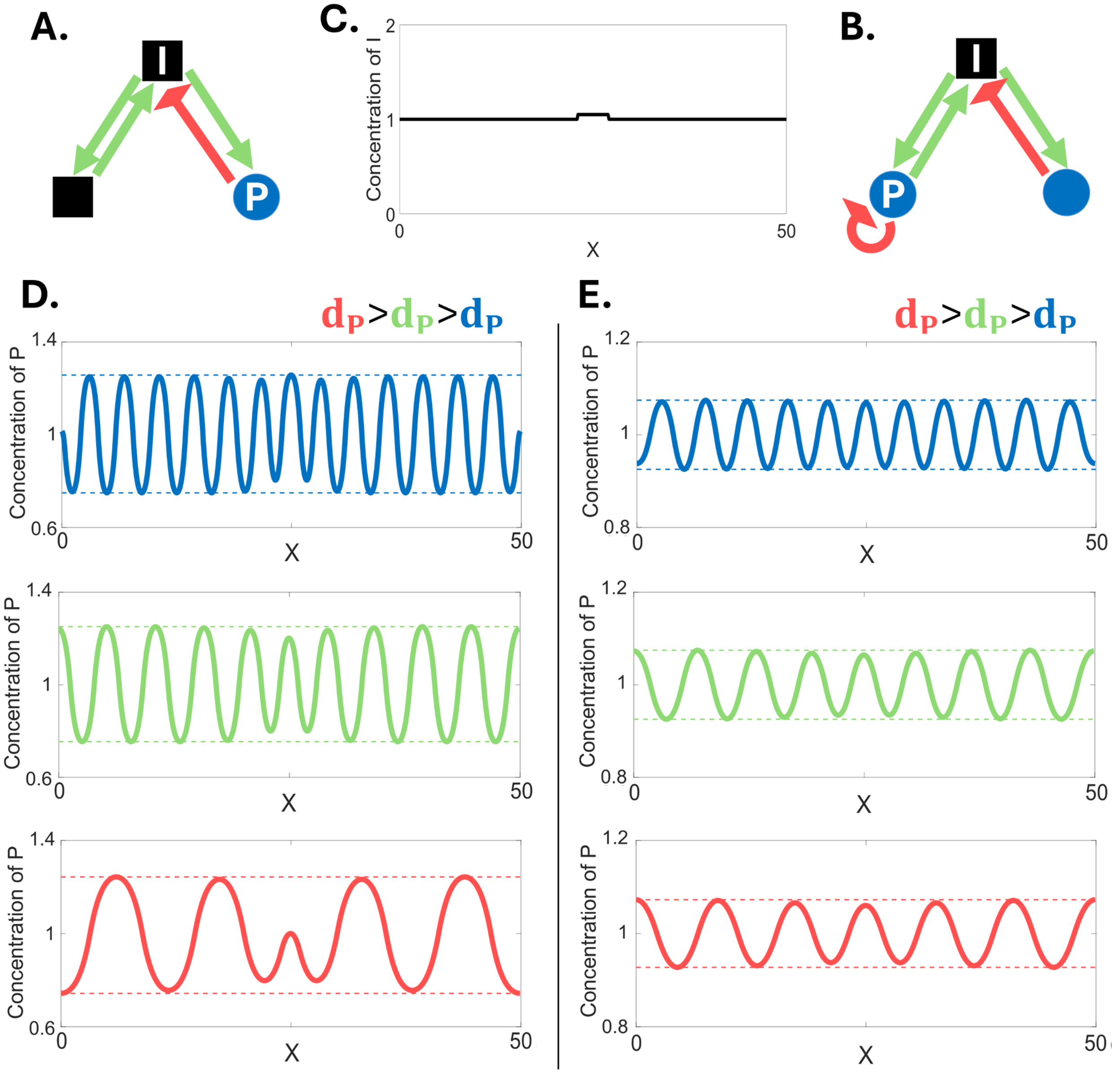


**Figure S4: Variational properties of emergent networks. (A-B)** An over-Turing (left) and Turing (right) gene networks produce a very similar periodical pattern from a spike-homogeneous initial pattern. A change in any model parameter (such as the diffusivity of an extracellular signal) alters the height and position of all peaks at the same time. **(C)** Spike initial pattern. **(D-E)**. Network colors and shapes as in Figure S1. $P$ stands for the gene product plotted as resulting pattern while $I$ stands for the gene product in the initial pattern. Simulations as in Figure S3 (see section S6 for parameter values).

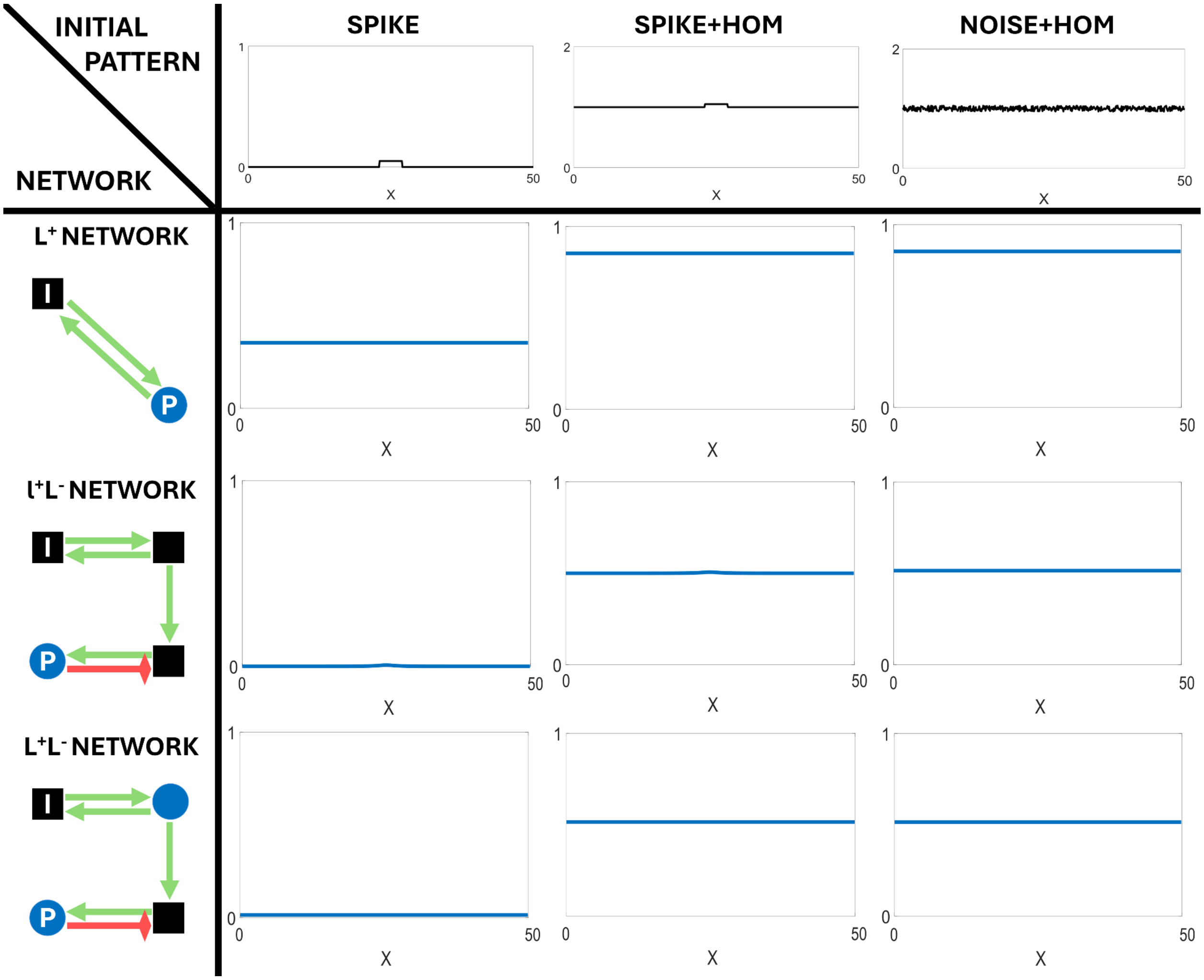


**Figure S5: $L^+$, $I^+L^-$ and $L^+L^-$ networks do not produce non-trivial pattern transformations.** The first column depicts simple examples of each type of these gene network topologies. The upper row shows the three initial patterns. Intermediate panels show each type of possible resulting pattern arising from each combination of initial pattern and gene network topology. Note that all pattern transformations are trivial. Network colors and shapes as in Figure S1. The y-axis in all plots represents gene product concentrations. $P$ stands for the gene product plotted as resulting pattern while $I$ stands for the gene product in the initial pattern. Simulations as in Figure S3 (see section S6 for parameter values).

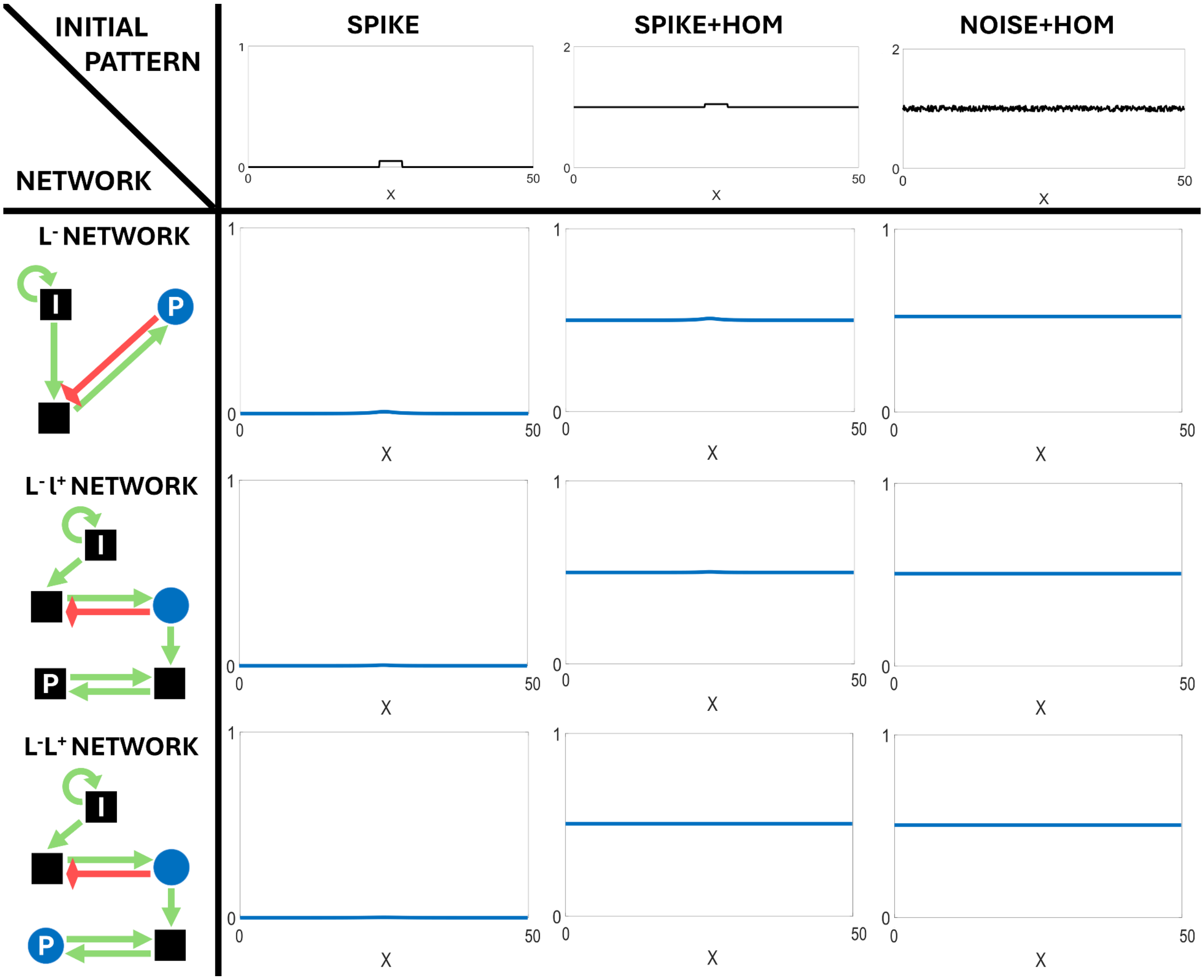


**Figure S6: L⁻, L⁻l⁺ and L⁻L⁺ networks do not produce non-trivial pattern transformations.** The first column depicts simple examples of each type of these gene network topologies. The upper row shows the three initial patterns. Intermediate panels show each type of possible resulting pattern arising from each combination of initial pattern and gene network topology. Note that all pattern transformations are trivial. Network colors and shapes as in Figure S1. The y-axis in all plots represents gene product concentrations. $P$ stands for the gene product plotted as resulting pattern while $I$ stands for the gene product in the initial pattern. Simulations as in Figure S3 (see section S6 for parameter values).
